# Genetic dissection of persiathiacin biosynthesis defines hierarchical P450 oxidations and reveals a more potent antitubercular intermediate

**DOI:** 10.64898/2026.07.09.737402

**Authors:** Felaine A. Sumang, Maxwell T. Stevens, Warwick J. Britton, Jeff Errington, Yousef Dashti

**Affiliations:** School of Medical Sciences, Faculty of Medicine and Health, The University of Sydney, Sydney NSW 2006, Australia; Tuberculosis Research Program at the Centenary Institute, The University of Sydney, Sydney NSW 2006, Australia; Department of Clinical Immunology, Royal Prince Alfred Hospital, Sydney NSW 2050, Australia; Sydney Infectious Diseases Institute, The University of Sydney, Sydney NSW 2006, Australia

**Keywords:** RiPP, Antibiotic, Cytochrome P450, *M. tuberculosis*, MRSA

## Abstract

Thiopeptides are ribosomally synthesized and post-translationally modified peptides (RiPPs) that form complex bioactive scaffolds through extensive enzymatic tailoring. The polyglycosylated thiopeptides persiathiacins, exhibit potent activity against multidrug-resistant *Mycobacterium tuberculosis* (Mtb) and methicillin-resistant *Staphylococcus aureus* (MRSA). The persiathiacin biosynthetic gene cluster encodes six cytochrome P450 (CYP) enzymes, but the logic of their oxidative modifications was unknown. Here, we establish a protoplast-based genetic system for *Actinokineospora* and systematically assign functions to all P450s. We demonstrate that PerX hydroxylates the central thiazole, PerV installs the third indole–core crosslink required for macrocyclization, and PerT, not PerU, catalyses indole *N*-hydroxylation. Combined gene inactivation and metabolite profiling reveal a hierarchical enzymatic sequence leading to the mature scaffold prior to sugar installation. Notably, the intermediate accumulating in the *ΩperX* mutant exhibits enhanced anti-*M. tuberculosis* potency compared to persiathiacin A (IC_50_ = 0.07 vs 1.5 µg mL^−1^). These results define the enzymatic logic and temporal organization of persiathiacin biosynthesis, providing a conceptual framework for rational diversification of complex thiopeptide natural products.

## Introduction

The structural and functional diversification of RiPPs arises from complex enzymatic tailoring reactions that convert simple precursor peptides into architecturally elaborate bioactive molecules. Among these, thiopeptides constitute one of the most structurally intricate RiPP families, distinguished by extensive heterocyclization, macrocyclization, and oxidative crosslinking reactions that generate rigid, densely functionalized scaffolds with potent antibacterial activity.^1-5^ Despite this potency, the clinical development of thiopeptides has been hindered by poor aqueous solubility and limited gastrointestinal absorption. Consequently, only thiostrepton and nosiheptide have reached application, primarily in topical formulations or as growth promoters in animal feed.^3^

The polyglycosylated thiopeptide antibiotics persiathiacin A (**1**) and B (**2**) (Figure 1A), produced by *Actinokineospora* sp. UTMC 2448, exhibit potent activity against *M. tuberculosis* and MRSA.^6,7^ Their activity is comparable to structurally related thiopeptides nocathiacin I (**3**) and nosiheptide (**4**) (Figure 1A), indicating that extensive glycosylation does not compromise antibacterial activity and highlights opportunities for biosynthetic engineering to generate novel (poly)glycosylated thiopeptides with improved pharmacological properties. The persiathiacin biosynthetic gene cluster (*per*) shares close homology with the *noc* and *nos* BGCs involved in the production and maturation of nocathiacin I and nosiheptide (Figure 1B).^8,9^ Biosynthesis begins with ribosomal production of the precursor peptide PerM (**5**), comprising an N-terminal leader peptide and a C-terminal core containing residues extensively modified in the mature scaffold (Figure 1C).^1,6,10,11^ Core maturation starts with cyclodehydration of cysteine residues by the YcaO cyclodehydratase complex PerG/PerH, followed by oxidation of the resulting thiazolines to thiazoles by the dehydrogenase PerF. Selective dehydration of serine and threonine residues is proposed to be catalysed by PerD and PerE.^6^

**Figure 1.**
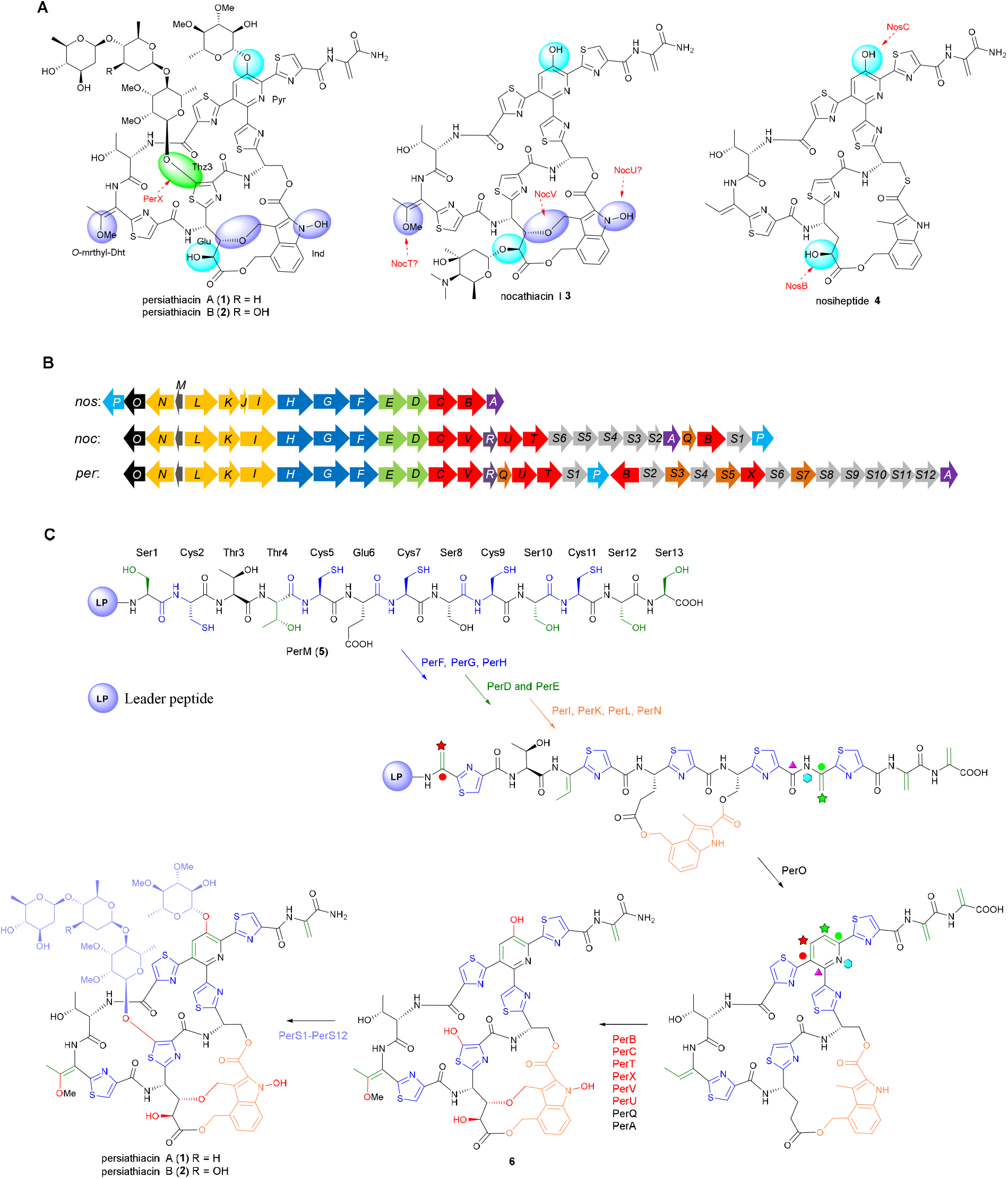
**(A)** Structures of persiathiacins A (**1**) and B (**2**), nocathiacin I (**3**), and nosiheptide (**4**). Sites modified by cytochrome P450 enzymes encoded in each BGC are color-coded: cyan circles indicate hydroxyls installed by conserved CYP homologues in all three pathways (NosB/NocB/PerB and NosC/NocC/PerC); purple ovals mark oxidative modifications present in *noc* and *per* clusters but absent in *nos* (NocT/PerT, NocV/PerV, and NocU/PerU); a green circle highlights the hydroxyl introduced by PerX, unique to the persiathiacin BGC. **(B)** Comparative organization of the persiathiacin (*per*), nocathiacin I (*noc*), and nosiheptide (*nos*) BGCs. Homologous genes are shown in matching colours. **(C)** Proposed biosynthetic pathway of persiathiacin based on prior knowledge. This schematic summarizes the functional assignments and order of oxidative tailoring steps inferred from sequence homology, heterologous assays, and chemical logic; the experimentally revised pathway established in this study is presented below.

Installation of the 3,4-dimethylindolic acid (DMIA) moiety, derived from tryptophan, is mediated by PerI, PerK, PerL, and PerN. Macrocycle formation and construction of the central pyridine core occur concomitantly with leader peptide elimination, via a formal [4+2] cycloaddition catalysed by PerO, a homologue of NosO and NocO. The scaffold is further matured by PerA, which catalyses C-terminal amide formation, together with multiple cytochrome P450 enzymes responsible for late-stage hydroxylations and oxidative crosslinking to generate **6**. Finally, a suite of sugar biosynthetic enzymes and glycosyltransferases encoded by *perS1*–*perS12* assembles and installs four highly modified 6-deoxysugars, producing the distinctive polyglycosylated scaffold of persiathiacins.^6^

The persiathiacin biosynthetic gene cluster encodes six P450 enzymes, an unusually large complement for a single RiPP pathway, raising questions about how multiple oxidative enzymes are temporally and functionally coordinated. Initial functional assignments, based on homologues in the nosiheptide and nocathiacin pathways, and chemical logic, proposed roles in glutamic acid hydroxylation (PerB), pyridine hydroxylation (PerC), thiazole hydroxylation (PerX), indole *N*-hydroxylation (PerU), oxidative crosslink formation (PerV), and dehydrothreonine hydroxylation (PerT), the latter proposed to precede methylation (Figure 1A).^6^ However, these assignments remained speculative, particularly for PerX, whose function was inferred from heterologous assays employing nosiheptide as a surrogate substrate.^6^

Here, we establish a protoplast-based genetic platform for *Actinokineospora* sp. UTMC 2448 and apply it to systematically dissect the functions of persiathiacin-associated P450 enzymes in their native biosynthetic context. Using targeted gene inactivation combined with comprehensive metabolite profiling, we define the specific oxidative transformations catalysed by each P450. We confirm that PerX hydroxylates the thiazole, identify PerV as the enzyme responsible for installing the third indole–core peptide crosslink required for macrocyclization, and reassign indole *N*-hydroxylation from PerU to PerT. Analysis of pathway intermediates further delineates the temporal sequence of oxidative tailoring events. Together, these findings reveal how multiple P450 enzymes are hierarchically coordinated to program oxidative diversification of the persiathiacin scaffold, establishing a framework for the rational engineering of thiopeptide natural products.

## Results and discussion

### Genetic validation of PerX as the thiazole-hydroxylating cytochrome P450

The *per* cluster encodes six cytochrome P450 enzymes: PerB, PerC, PerT, PerU, PerV, and PerX. PerB and PerC have homologues in both *nos* and *noc*, whereas PerT, PerU, and PerV are found in *noc* but absent from *nos*. Notably, PerX is unique to the *per* cluster and was thus proposed to catalyse a persiathiacin-specific oxidative transformation. Bioinformatic analysis and chemical logic suggested that PerX hydroxylates the central thiazole (Thz3), generating the attachment site for the trisaccharide moiety (Figure 1).

In our previous study, direct genetic interrogation of *perX* was precluded by limited genetic tools for *Actinokineospora*. To probe its function, we first assayed recombinant His_6_-tagged PerX *in vitro* using nosiheptide (**4**) as a surrogate substrate. Reactions were conducted with spinach ferredoxin, ferredoxin reductase, and NADPH to reconstitute electron transfer. LC– HRMS analysis revealed a +16 Da mass shift, consistent with monooxygenation of the nosiheptide scaffold.^6^ To obtain sufficient material for NMR characterization, *perX* was overexpressed under the constitutive *ermE\** promoter in the native nosiheptide producer *Streptomyces actuosus* ATCC25421. LC–HRMS of culture extracts confirmed production of the same oxygenated derivative observed *in vitro*. However, isolation of sufficient material for full NMR characterization proved unsuccessful.^6^ These results supported a role for PerX in scaffold oxidation, but definitive assignment of the hydroxylation site remained elusive.

To enable direct *in vivo* functional analysis, we established a protoplast-based transformation system for *Actinokineospora* sp. UTMC 2248. A disruption construct was generated from pSET152 by replacing the *Φ*C31 integrase with a 2 kb internal fragment of *perX*, designed to yield single-crossover insertional inactivation.^12,13^ To minimize polar effects on downstream genes, a strong constitutive promoter *kasOp\** was introduced at the 3′ end of the fragment.^14^ Initial attempts at intergeneric conjugation from *E. coli* ET12567/pUZ8002 were unsuccessful. We therefore adopted a protoplast transformation approach.^15^ Lysozyme treatment in protoplast buffer generated spherical protoplasts, confirming efficient cell wall removal (Figure 2A). PEG-mediated transformation yielded apramycin-resistant colonies after 4 days. One transformant, designated *Actinokineospora* sp. UTMC 2248-*ΩperX*, was selected for metabolic analysis.

**Figure 2.**
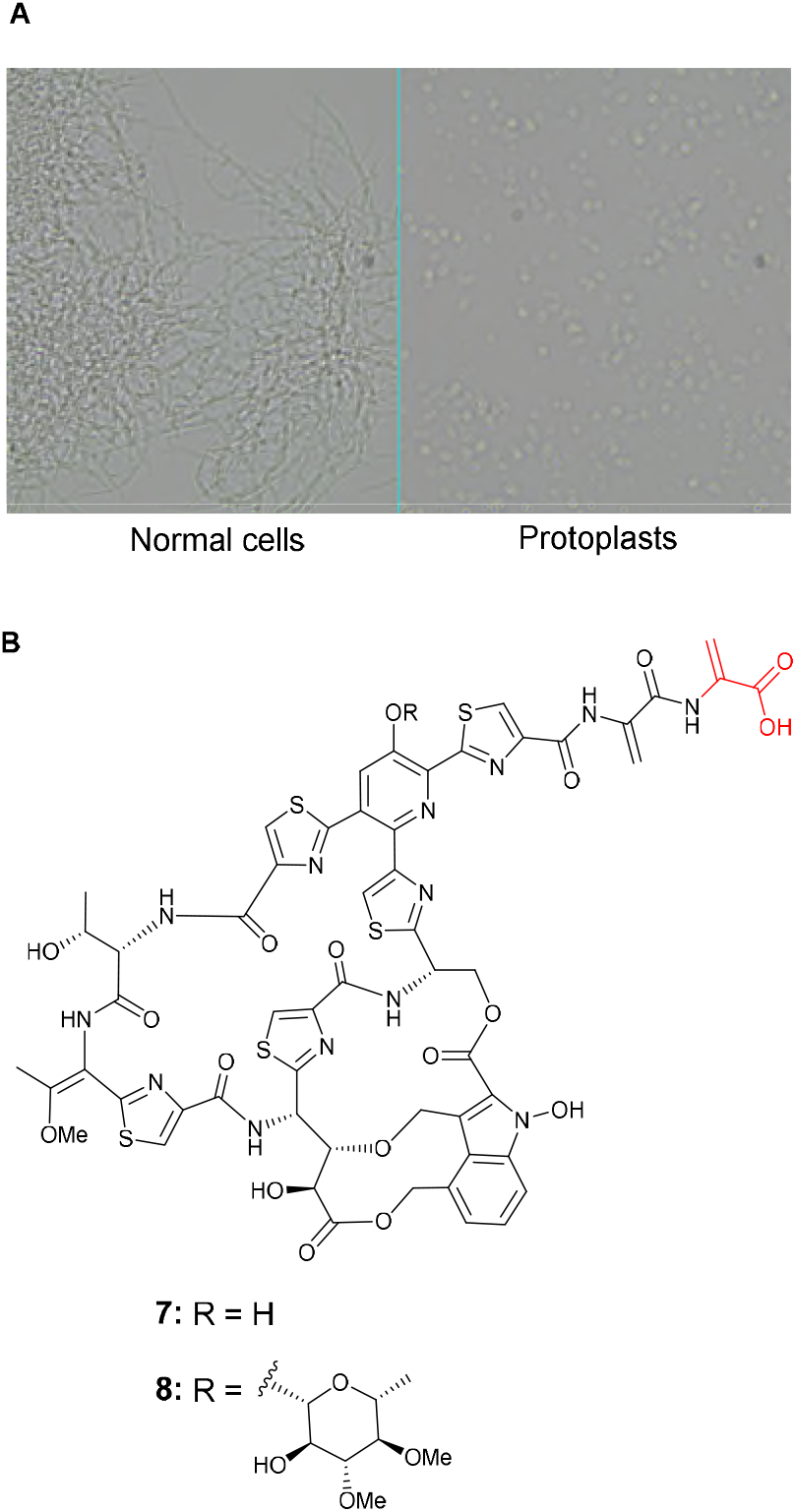
**(A)** Microscopic image of *Aktinokineospora* sp. UTMC 2248 normal cells compared to protoplasts generated after 30 minutes of treatment with lysozyme in protoplast buffer. **(B)** Structures of metabolites **7** and **8** accumulated in culture extract of *Actinokineospora* sp. UTMC 2248-*ΩperX*.

LC–HRMS comparison of wild-type and mutant extracts revealed complete abolition of persiathiacin A and B production in *ΩperX*. Instead, two new metabolites **7** and **8** (Figure 2B) were detected, with molecular formulae consistent with deshydroxy, minimally glycosylated persiathiacin intermediates. Large-scale fermentation enabled purification of the major compound **7** for full structural characterization. Metabolite **7** lacked all anomeric proton signals in the ^1^H NMR spectrum, indicating absence of sugar residues. 1D and 2D NMR analysis (Table S3 and Figures S1, S7, and S15-S19) combined with HR-MS data (Figure S7) confirmed a core structure resembling persiathiacin, with two key differences: (i) the Thz3 unit remained unmodified, as indicated by the Thz3-H5 resonance at *δ*_*H*_ 8.25 and HMBC correlations to Thz3-C4 (*δ*_*C*_ 150.6) and the adjacent carbonyl (*δ*_*C*_ 160.5); (ii) an additional dehydroalanine (Dha) residue was present at the C-terminus, as indicated by characteristic sp^2^ methylene protons at *δ*_*H*_ 5.78 and 6.23, which show HSQC correlations to *δ*_*C*_ 104.9. The molecular formulae C_63_H_59_N_13_O_22_S_5_ of minor metabolite **8** supports the same aglycone bearing a single deoxysugar on the pyridine ring, consistent with fragmentation patterns yielding aglycone **7** (Figure S8).

Notably, the second Dha residue is normally removed during persiathiacin maturation by PerA. The accumulation of **7** and **8** in the *ΩperX* mutant indicates that these intermediates are poor substrates for PerA, suggesting that Thz3 hydroxylation (and possibly glycosylation at this position) precedes PerA-mediated processing. Furthermore, the higher abundance of **7** relative to **8** suggests that the non-hydroxylated scaffold is a poor substrate for the glycosyltransferase responsible for pyridine-ring modification.

### Catalysis of the ether linkage between DMIA and the core peptide by PerV CYP precedes macrocyclization

Having established a protoplast-based genetic system for *Actinokineospora* sp. UTMC 2248, we next interrogated the remaining CYP enzymes encoded within the *per* biosynthetic gene cluster. Among these, PerV shares homology with NocV, a CYP proposed to catalyze formation of the ether linkage between DMIA and the core peptide during nocathiacin biosynthesis. *In vivo* heterologous expression studies implicated *nocV* in ether bond formation in a genetically tractable nosiheptide-producing strain; however, *in vitro* assays with purified NocV and nosiheptide intermediates failed to generate the ether-linked product, leaving the timing of this transformation unresolved.^16^

To define the role of PerV in persiathiacin biosynthesis, we constructed a disruption plasmid containing an ∼2 kb internal fragment of *perV* and introduced it into wild-type *Actinokineospora* sp. UTMC 2248 via protoplast-mediated transformation. The resulting mutant strain, *Actinokineospora* sp. UTMC 2248-*ΩperV*, was subjected to comparative metabolic profiling against the wild type. LC–HRMS analysis revealed complete loss of the production of persiathiacins A and B in the *ΩperV* mutant, and accumulation of a new metabolite **9** (Figure 3) with the molecular formula C_53_H_49_N_13_O_15_S_5_. Large-scale fermentation enabled isolation of this compound, and comprehensive NMR analysis (Table S4 and Figures S2, S9, and S20-S24) established its structure. As anticipated, **9** lacks the ether linkage between DMIA and the peptide core. Strikingly, the characteristic macrocyclic and pyridine rings were also absent. Thus, disruption of PerV prevents not only ether formation but also macrocyclization.

**Figure 3.**
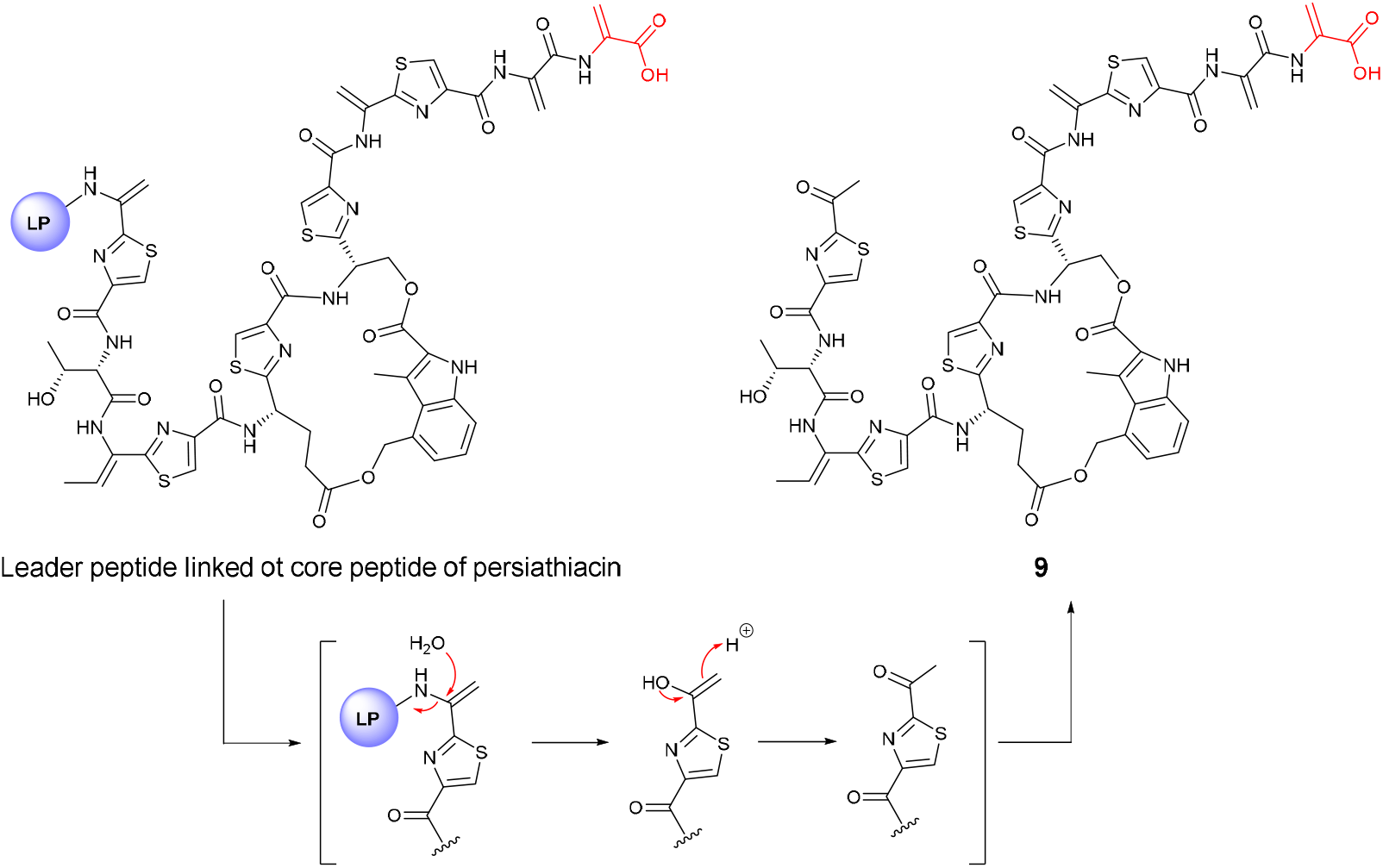
Structure of compound **9** accumulated in the culture extract of *Actinokineospora* sp. UTMC 2248-*ΩperV*. The proposed hydrolytic cleavage of the leader peptide and subsequent keto–enol tautomerization to the more stable ketone form are shown.

In persiathiacin biosynthesis, macrocyclization is proposed to proceed via a formal [4+2] cycloaddition catalysed by PerO, generating the central pyridine core concomitant with leader peptide removal. The failure of macrocycle formation in **9** indicates that installation of the DMIA–peptide ether bridge is a prerequisite for PerO activity. We therefore conclude that PerV-catalysed ether formation precedes, and likely enables, PerO-mediated macrocyclization. This ordering is consistent with prior studies of NocV, in which incubation with fully cyclized nosiheptide substrates did not yield ether-linked products,^16^ suggesting that ether formation does not occur after macrocyclization.

Notably, compound **9** contains an N-terminal ketone. This feature is consistent with hydrolytic removal of the leader peptide, followed by keto–enol tautomerization to the thermodynamically favoured ketone (Figure 3). In addition, **9** lacks the oxidative tailoring modifications normally installed by the remaining P450 enzymes encoded in the *per* cluster. These observations indicate that the linear, non-etherified intermediate, whether free or transiently associated with the leader peptide, is not a competent substrate for downstream CYP-catalysed transformations.

Rather, macrocyclization appears to establish the conformational framework required for late-stage oxidative tailoring.

Consistent with intermediates **7** and **8** identified in the *ΩperX* mutant, compound **9** retains the second C-terminal Dha residue. Collectively, these findings define a revised temporal sequence for persiathiacin (and likely nocathiacin) biosynthesis: (i) PerV installs the DMIA–peptide ether bridge; (ii) PerO subsequently catalyses pyridine formation and macrocyclization; and (iii) late-stage oxidative tailoring and Dha removal proceed only after macrocycle assembly. This work clarifies the ambiguity surrounding the timing of ether formation in thiopeptide biosynthesis and establishes PerV (and likely NocV) as a key gatekeeper enzyme controlling entry into the macrocyclization phase.

Despite the requirement for the DMIA–core ether bridge in PerO and NocO-dependent macrocyclization, the homologous enzyme NosO from the nosiheptide pathway catalyses pyridine formation in its absence, indicating functional divergence within this enzyme family across different biosynthetic contexts.

### Revising the function of PerU: *N*-hydroxylation of the indolic moiety is catalysed by PerT

PerU and PerT are two additional cytochrome P450 enzymes encoded within the *per* biosynthetic gene cluster that share homology with NocU and NocT from the nocathiacin pathway. In the *noc* system, NocU has been proposed to catalyse *N*-hydroxylation of the indolic nitrogen based on *in vivo* heterologous expression in nosiheptide-producing strain *S. octuosus* SL5001, which yielded a hydroxylated nosiheptide analogue. Structural assignment by NMR and MS/MS led to the conclusion that hydroxylation occurred at the indole nitrogen.^17^ By analogy, PerU was therefore presumed to catalyse indole *N*-hydroxylation in persiathiacin biosynthesis, whereas PerT (the NocT homolog) was proposed to hydroxylate the dehydrothreonine (Dht) residue prior to *O*-methylation by PerQ to generate the *O*-methyl-Dht moiety.

To test this model directly, we first disrupted *perT*. Unexpectedly, LC–HRMS analysis of the *ΩperT* mutant revealed production of a metabolite (**10**) (Figure 4) whose molecular formula differed from persiathiacin A by loss of a single oxygen atom. Large-scale fermentation and NMR characterization (Table S5 and Figures S3, S10, and S25-S29) demonstrated that the *O*-methyl-Dht moiety remained intact, excluding a role for PerT in Dht hydroxylation. Instead, pronounced changes were observed in the indole region. Specifically, the ^13^C chemical shifts of Ind-C3 (*δ*_*C*_ 116.5), Ind-C3a (*δ*_*C*_ 123.9), and Ind-C7 (*δ*_*C*_ 116.0) were significantly downfield relative to persiathiacin A (*δ*_*C*_ 109.8, 119.5, and 112.1, respectively). Moreover, the indolic NH proton (*δ*_*H*_ 10.18) exhibited clear ^*2*^*J* and ^*3*^*J* HMBC correlations to Ind-C2, Ind-C7a, Ind-C3, and Ind-C3a, consistent with an unsubstituted indole nitrogen. Together, these data unequivocally demonstrate that the *N*-hydroxyl group is absent in the *ΩperT* metabolite. We therefore conclude that PerT, rather than PerU, catalyses indole *N*-hydroxylation.

**Figure 4.**
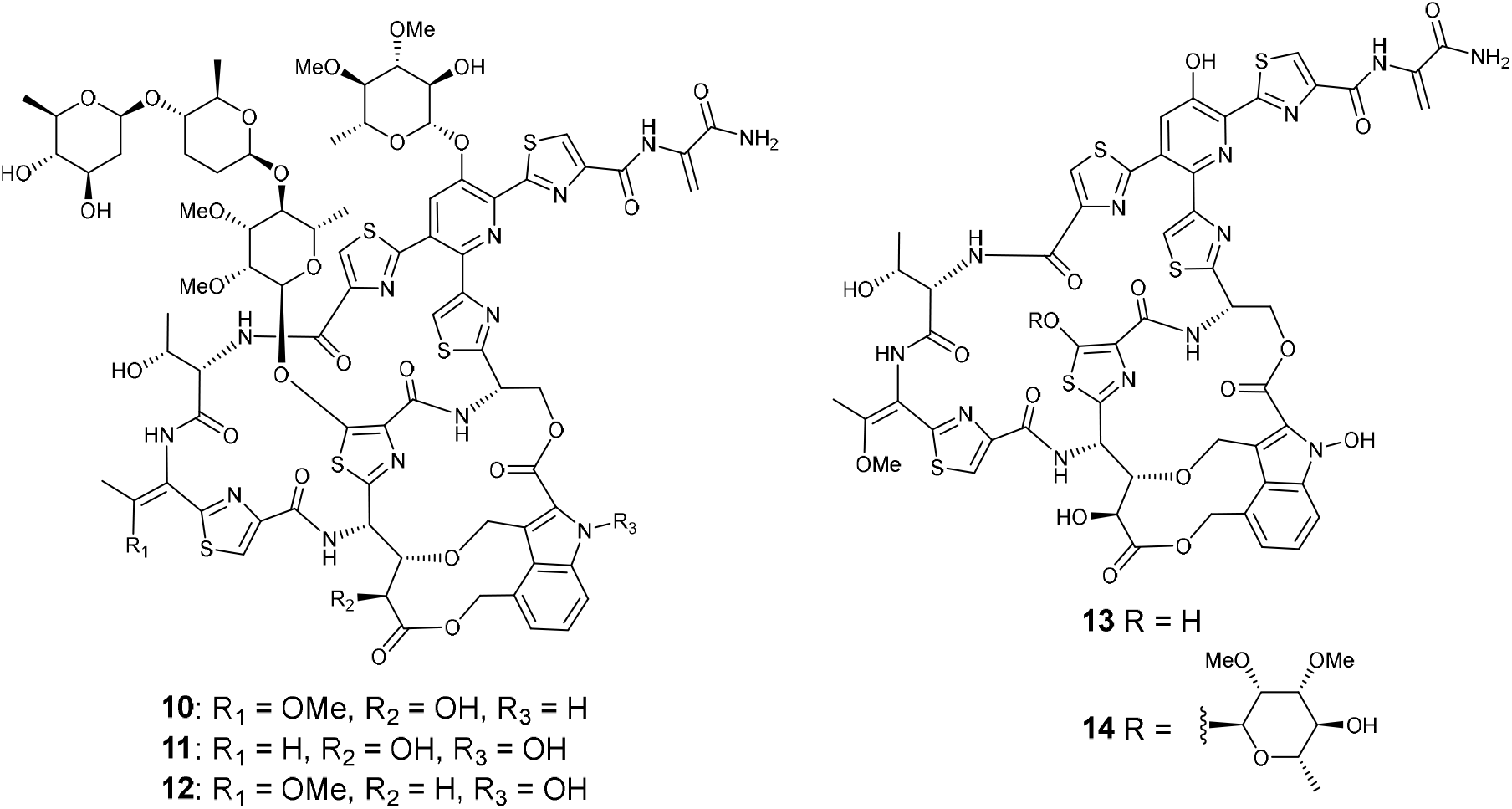
Structures of metabolites **10**–**12** isolated from the CYP deletion mutants *ΩperT, ΩperU*, and *ΩperB*, respectively, and structures **13** and **14** from the glycosyltransferase deletion mutant *ΩperS4*.

This unexpected finding prompted re-examination of the original functional assignment of NocU.^17^ Comparison of the reported NMR data for the NocU-derived hydroxylated nosiheptide analogue (recorded in THF-*d*_*8*_) with literature data for nosiheptide (recorded in DMSO-*d*_6_)^18^ revealed nearly identical chemical shifts in the indole region despite the use of different solvents. Furthermore, the Dht methyl resonance, expected to appear as a doublet, was reported as a singlet.^17^ These inconsistencies call into question the structural assignment of the hydroxylated product and suggest that the role of NocU may have been misinterpreted.

We next disrupted the *perU* gene. This deletion abolished production of persiathiacins A and B and yielded a minor metabolite whose molecular formula (C_79_H_89_N_13_O_29_S_5_) was consistent with loss of a methoxy substituent. Although repeated scale-up fermentations did not provide sufficient material for complete NMR characterization, HRMS data (Figure S11) supported assignment of this compound as the desmethoxy analogue **11** (Figures 4). These findings indicate that PerU catalyses hydroxylation of the Dht residue, which is subsequently methylated by PerQ to furnish the *O*-methyl-Dht moiety. These results revise the functional assignments of the two P450 enzymes in the *per* pathway: PerT catalyses indole *N*-hydroxylation, whereas PerU mediates Dht hydroxylation prior to *O*-methylation. This reassignment overturns the biosynthetic logic previously inferred from the nocathiacin system and underscores the limitations of homology-based functional annotation in the absence of direct genetic and structural validation.

The remaining cytochrome P450 enzymes encoded within the *per* biosynthetic gene cluster, PerB and PerC, share close homology with NosB/NosC from the nosiheptide pathway and NocB/NocC from the nocathiacin system.^6^ In nosiheptide biosynthesis, NosB catalyses hydroxylation at C4 of the glutamic acid residue, whereas NosC mediates hydroxylation of the pyridine ring.^19^ The corresponding enzymes in the *noc* pathway were similarly validated to hydroxylate these positions.^20,21^

We generated an *ΩperB* mutant to determine whether PerB performs an analogous role in persiathiacin assembly. LC–MS analysis revealed accumulation of a metabolite (**12**) (Figure 4) consistent with deshydroxy-persiathiacin A. Purification and comprehensive NMR characterization (Table S3 and Figures S4, S12, and S30-S34) demonstrated the absence of the C4 hydroxyl group on the glutamic acid residue. In the ^13^C NMR spectrum, C4 appeared as a methylene carbon at *δ*_*C*_ 37.9, with corresponding ^1^H signals at *δ*_*H*_ 2.44 and 2.69. HMBC correlations from these protons to C2 (*δ*_*C*_ 52.2), C3 (*δ*_*C*_ 79.2), and C5 (*δ*_*C*_ 172.0) of the glutamic acid residue unambiguously established dehydroxylation at this position. These results confirm that PerB catalyses C4 hydroxylation of the glutamate residue in persiathiacin biosynthesis. Based on the functionally validated homologs NosC and NocC and the roles of other CYPs in the *per* cluster, the remaining CYP, PerC, is assigned to pyridine hydroxylation.

### Temporal hierarchy of CYP-catalysed and late-stage transformations

Integration of the CYP mutant phenotypes provides insight into the temporal sequence of oxidative tailoring events. The *ΩperV* mutant uniquely accumulated a linear intermediate lacking both the DMIA–peptide ether linkage and the macrocyclic pyridine core, demonstrating that PerV operates prior to PerO-mediated macrocyclization. In contrast, deletion of *perB, perT*, or *perU* yielded fully macrocyclized, glycosylated analogues lacking specific oxidative modifications. These findings establish that PerV acts early, whereas PerB, PerT, and PerU function after macrocycle formation.

The *ΩperB* and *ΩperT* mutants accumulated deshydroxy analogues at titres comparable to native persiathiacins, indicating that removal of the glutamate C4 hydroxyl or indole *N*-hydroxyl group does not substantially impair downstream enzymatic processing. These modifications therefore appear to fine-tune scaffold functionality rather than govern pathway progression. In contrast, the *ΩperU* mutant produced a fully glycosylated desmethoxy analogue at significantly reduced yield. The absence of an open-chain intermediate analogous to that observed in *ΩperV* argues against a pre-macrocyclization role for PerU. Instead, these data support a model in which PerU acts after macrocyclization but is nevertheless important for maintaining efficient substrate recognition and pathway flux.

Deletion of *perX* resulted in accumulation of unglycosylated and monoglycosylated metabolites **7** and **8**, which retain the C-terminal bisDha motif, confirming that they are bona fide intermediates. Notably, fully oxidized scaffolds were observed in this background, indicating that CYP-mediated oxidative tailoring occurs prior to glycosylation. Together, these results establish that oxidative modifications are largely completed before sugar installation.

In our previous work, *in vitro* assays demonstrated that the glycosyltransferase PerS4 catalyses attachment of a deoxysugar to the hydroxyl group of the pyridine ring of nosiheptide as a surrogate substrate. To check this, we made an *ΩperS4* mutant and found that this strain accumulated fully oxidized metabolites **13** and **14** (Figures 4 and S13-S14). The lack of complete glycosylation of **13** confirmed that CYP-catalysed oxidative tailoring proceeds independently of glycosylation. Furthermore, detection of only the monoglycosylated intermediate **14** in the *ΩperS4* indicates that trisaccharide assembly at Thz3 requires prior installation of the pyridine-linked sugar. These results support a hierarchical model of glycosylation in which initial sugar attachment to both Thz3 and the pyridine ring establishes the structural context required for subsequent extension. We also deleted the putative methyltransferase *perS5*, which led to production of multiple thiopeptide analogues with altered glycosylation patterns (Figure S5). These observations indicate that deoxysugars are methylated prior to transfer, and that methylation influences the fidelity of downstream glycosylation events.

Analysis of multiple CYP and glycosylation mutants was used to define the timing and substrate requirements of PerA, the enzyme responsible for removal of the second C-terminal Dha residue. In the *ΩperX* and *ΩperV* mutants, the accumulated intermediates retained the bisDha motif, indicating that PerA does not act on substrates lacking Thz3 hydroxylation (**7** and **8**) or on **9**, which lacks the macrocycle and the DMIA–peptide ether linkage. These intermediates likely adopt altered conformations compared to the mature scaffold. In contrast, *ΩperB, ΩperT*, and *ΩperU* mutants produced metabolites **10**–**12** in which the second Dha residue has been efficiently removed. Moreover, in the *ΩperS4* mutant, Dha cleavage occurred despite incomplete glycosylation of **13** and **14**. Together, these results demonstrate that PerA activity is independent of glycosylation but strongly dependent on prior formation of the correct macrocyclic architecture. We conclude that PerA recognizes a conformationally mature scaffold and functions as a late-stage processing enzyme in persiathiacin biosynthesis.

### Revised biosynthetic model for persiathiacin

Our genetic and structural analyses, together with the recently reported post-translational modification logic of nosiheptide,^22^ define a revised and experimentally validated biosynthetic framework for persiathiacin assembly (Figure 5) that reveals a tightly ordered hierarchy of post-translational modifications. Biosynthesis begins with ribosomal production of the precursor peptide PerM (**5**), followed by canonical thiopeptide core maturation. **(i)** The PerG/PerH complex catalyses cyclodehydration of cysteine residues, and PerF oxidizes the resulting thiazolines to thiazoles, installing five thiazole rings to yield pentathiazolyl intermediate **15. (ii)** The radical SAM enzyme PerL generates 3-methylindolic acid (MIA), which is activated by PerI and transferred by the α/β-hydrolase fold protein PerK to Ser8 of **15** via O-MIAylation to afford **16. (iii)** The dehydratase pair PerDE processes Ser1, Thr4, Ser10, Ser12, and Ser13 to generate the penta-dehydrated intermediate **17. (iv)** The radical SAM enzyme PerN catalyses C1-unit insertion into MIA to form DMIA and simultaneously mediates its conjugation to Glu6 through ester bond formation, yielding **18. (v)** PerV then installs a third bridge between DMIA and Glu6 via ether bond formation to generate **19**, the substrate for **(vi)** pyridine synthase PerO, which forms the 26-membered macrocycle through a formal [4+2] cycloaddition followed by aromatization. **(vii)** The resulting thiopeptide intermediate **20** subsequently undergoes oxidative tailoring by cytochrome P450 enzymes PerBCTXU and methylation by PerQ to produce **6. (viii)** Finally, the C-terminal Dha is removed by PerA through C-N cleavage, and the scaffold is decorated with methylated deoxysugars generated by PerS1–PerS12.

**Figure 5.**
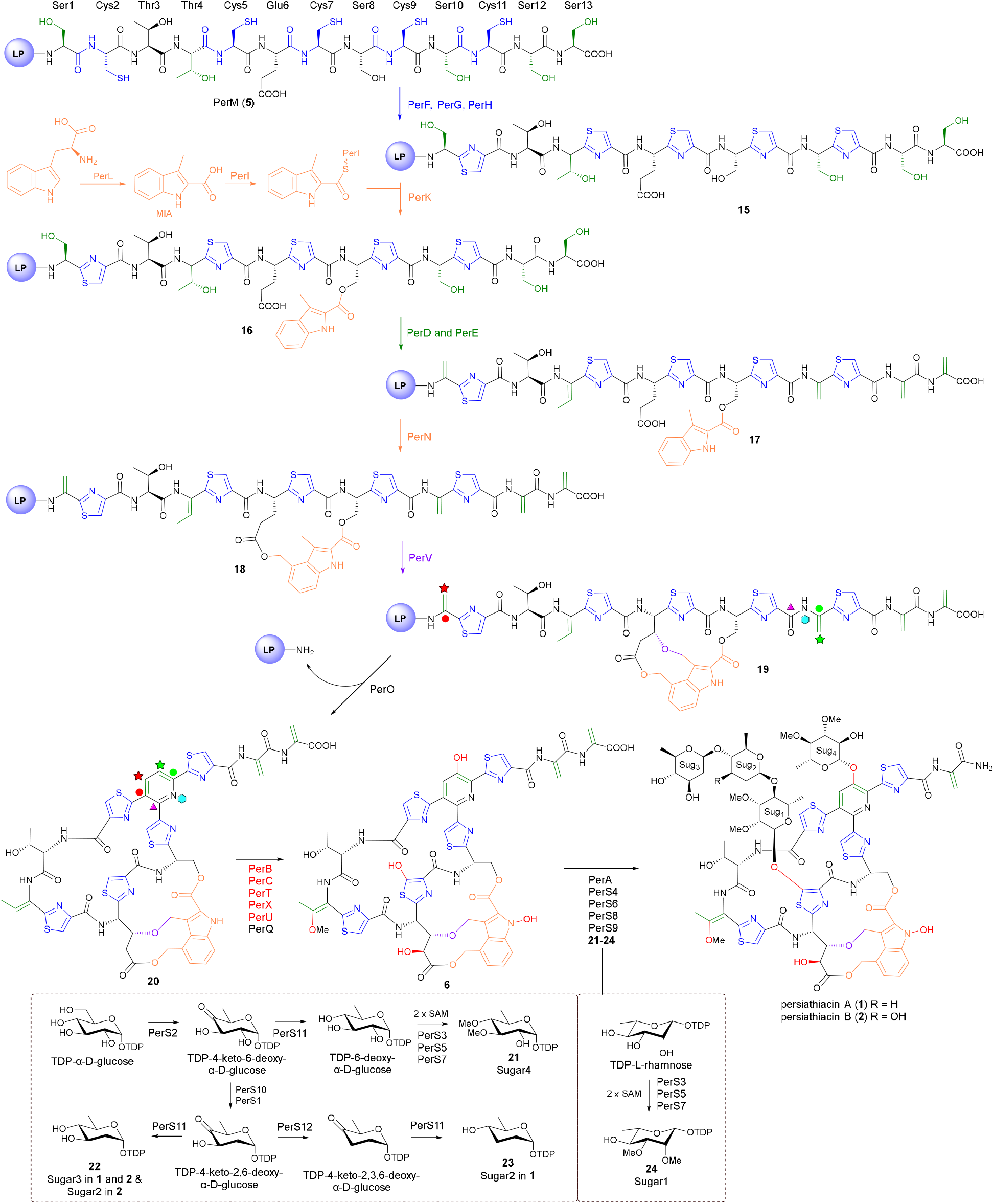
Revised biosynthesis of persiathiacin. The persiathiacin precursor peptide encoded by *perM* comprises an N-terminal leader peptide (LP) and a C-terminal core peptide (structure shown). Post-translational modifications catalysed by enzymes encoded within the *per* biosynthetic gene cluster convert the core peptide into the mature thiopeptide scaffold **6** in a hierarchical manner discussed in the main text. The biosynthesis of TDP-sugars **21–24** is shown in the dashed box. Methyltransferases PerS3, PerS5, and PerS7 *O*-methylate L-rhamnose and 6-deoxy-D-glucose prior to their attachment to the scaffold. PerS4 installs **21** on the pyridine ring, while PerS6, PerS8, and PerS9 decorate the scaffold with **24, 22**, and **23** to produce persiathiacin A (**1**) or with **24** and two units of **22** to produce persiathiacin B (**2**).

Our data demonstrate that oxidative ether bond formation between DMIA and the peptide backbone, catalysed by PerV, constitutes a pivotal early event that gates entry into macrocyclization. In the absence of PerV, the pathway stalls at a linear intermediate incapable of undergoing PerO-mediated pyridine formation. PerV-dependent crosslinking therefore establishes the conformational prerequisites for the formal [4+2] cycloaddition catalysed by PerO, which constructs the central pyridine core and completes macrocycle formation.

Following macrocyclization, a coordinated series of P450-catalysed oxidative transformations diversifies the scaffold. Functional reassignment of the CYP enzymes revealed that PerT catalyses indole *N*-hydroxylation, overturning prior homology-based predictions, whereas PerU hydroxylates the dehydrothreonine residue to enable subsequent *O*-methylation by PerQ. PerB installs the C4 hydroxyl group of the glutamate residue, PerC hydroxylates the pyridine ring, and PerX performs thiazole hydroxylation at Thz3. Importantly, accumulation of fully oxidized but unglycosylated intermediates in *ΩperX* and *ΩperS4* backgrounds demonstrated that oxidative tailoring is largely completed prior to sugar installation. Together, these findings define oxidative maturation as a post-macrocyclization but pre-glycosylation phase in which specific P450-mediated modifications contribute to conformational tuning of the scaffold.

Glycosylation follows a similarly ordered, hierarchical programme. The *perS1*–*perS12* locus encodes the enzymes responsible for deoxysugar biosynthesis and their sequential installation onto the mature thiopeptide core, as detailed in our previous study (Figure 5).^6^ Genetic disruption of the methyltransferase PerS5 resulted in accumulation of multiple glycosylated congeners (Figure S5), demonstrating that methylation of deoxysugar donors precedes glycosyl transfer. TDP-6-deoxy-α-D-glucose and TDP-L-rhamnose are regioselectively *O*-methylated by the methyltransferases PerS3, PerS5, and PerS7 en route to donors **21** and **24** prior to glycosyl transfer (Figure 5). The precise methylation sites for PerS3, PerS5, and PerS7 have not yet been determined. These methylated donors, together with D-olivose (**22**) and D-amicetose (**23**), are subsequently installed onto the persiathiacin scaffold by the glycosyltransferases PerS4, PerS6, PerS8, and PerS9.

Our data identified PerS4 as responsible for installation of the pyridine-linked sugar, a priming event required for efficient elaboration of the trisaccharide appended at Thz3. Thus, glycosylation is not a stochastic decorating process but a programmed sequence in which the initial sugar attachment establishes the structural and kinetic framework for subsequent glycosyltransferase-mediated extension at Thz3.

PerA mediates the final maturation of the persiathiacin scaffold by removing the second C-terminal dehydroalanine residue. Genetic studies indicated that its activity is independent of glycosylation yet strictly contingent upon the macrocycle adopting the correct conformation. Consequently, PerA acts on a conformationally mature intermediate, functioning as a late-stage enzyme that finalizes the persiathiacin scaffold.

Overall, this revised pathway reveals a biosynthetic logic in which **(i)** PerV-dependent ether formation enables macrocyclization; **(ii)** macrocyclization establishes the structural platform for P450-mediated oxidative diversification; **(iii)** oxidative maturation precedes hierarchical glycosylation of premethylated deoxysugars; and **(iv)** PerA selectively acts on a conformationally mature intermediate, with activity independent of glycosylation. This work redefines the functional roles and temporal coordination of six cytochrome P450 enzymes within a single RiPP pathway and illustrates how oxidative transformations shape conformational maturation to ensure faithful downstream enzymatic processing.

### Persiathiacin biosynthetic intermediate show superior antimycobacterial activity

As persiathiacin A exhibits activity against both drug-susceptible and drug-resistant clinical strains of *M. tuberculosis*, we evaluated the purified persiathiacin analogues obtained in this study (excluding the minor metabolite **8** and **11**) against *M. tuberculosis* H37Rv using the resazurin microtiter assay (Figure 6A).^23^ The linear analogue **9** completely lost activity, consistent with studies showing that disruption of the macrocyclic scaffold abolishes binding to ribosomal protein L11 and impairs inhibition of translation.^24^ Notably, compound **7** showed significantly enhanced potency, with an IC_50_ of 0.07 µg mL^−1^ compared to 1.5 µg mL^−1^ for persiathiacin A, representing an approximately 21-fold improvement in activity. Despite its markedly enhanced potency against *M. tuberculosis*, compound **7** retained anti-MRSA activity similar to that of **1**. The increase in activity against *M. tuberculosis* may be associated with the presence of the second dehydroalanine (Dha) residue, although its precise contribution remains unclear. However, intracellular assays of **1** and **7** against *M. tuberculosis* H37Rv lux in THP-1 macrophages showed similar activity (Figure 6B), suggesting that the degree of macrophage penetration may restrict their intracellular potency. Time kill kinetics of the compounds against *M. tuberculosis* strain H37Rv lux were assessed over 28 days at increasing concentrations of the compounds that inhibited regrowth of the mycobacteria (Figure 6C).^25^ The regrowth of *M. tuberculosis* H37Rv lux occurred at a lower concentration (625 nM) for **7** than for **1**, and the remaining compounds displayed activity comparable to **1**.

**Figure 6.**
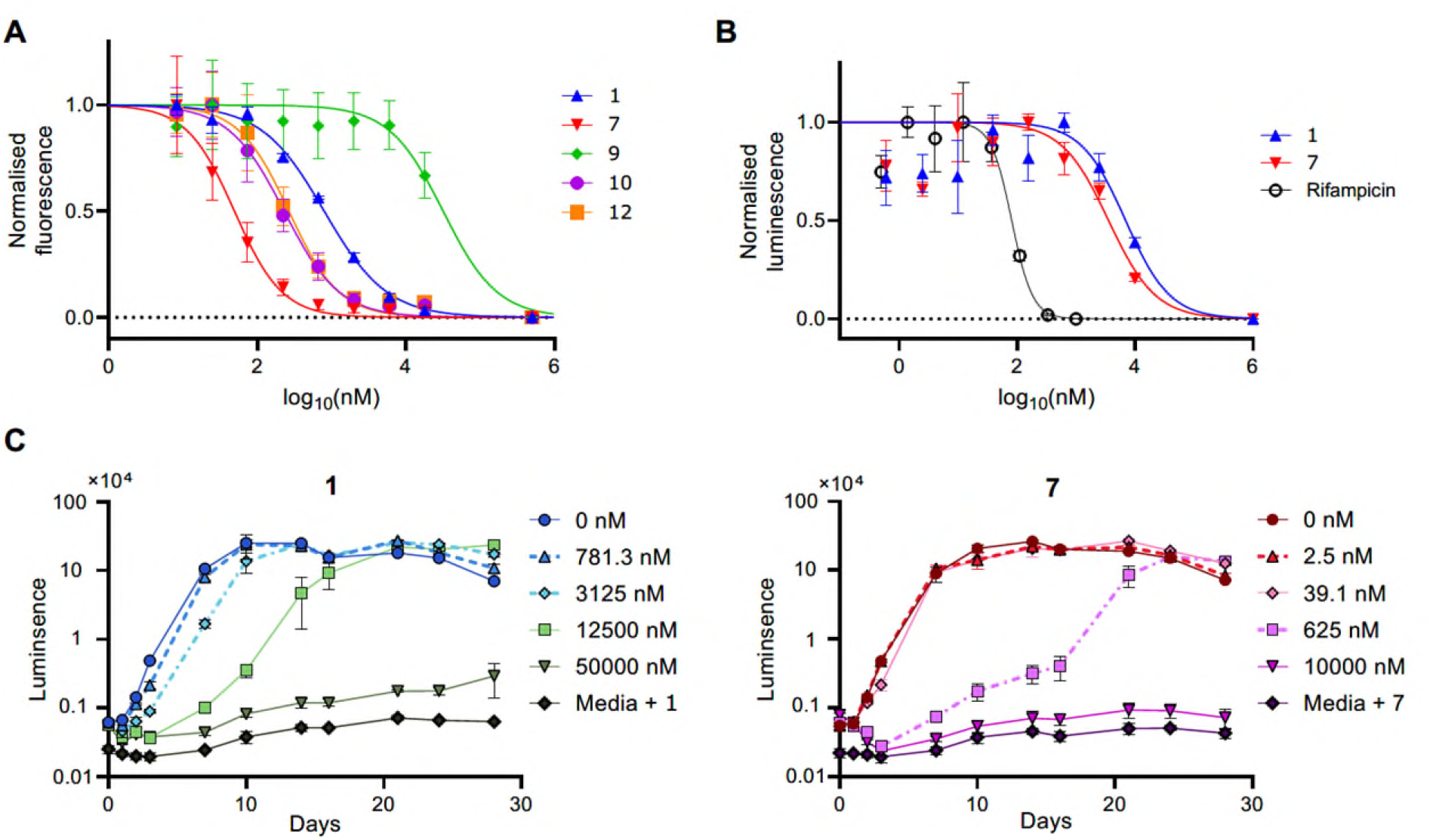
Antimycobacterial activity of persiathiacin analogues. **(A)** *In vitro* activity of purified persiathiacin analogues against *M. tuberculosis* H37Rv measured using the resazurin microtiter assay. Compound **7** showed enhanced potency relative to persiathiacin A (**1**), whereas the linear analogue **9** was inactive. **(B)** Intracellular activity of **1** and **7** against *M. tuberculosis* H37Rv lux in THP-1 macrophages. **(C)** Time–kill kinetics of compounds **1** and **7** against *M. tuberculosis* H37Rv lux.

## Conclusions

In this study, we establish genetically tractable platform for *Actinokineospora* sp. UTMC 2248 and use it to systematically define the functions of all cytochrome P450 enzymes encoded within the persiathiacin biosynthetic gene cluster. Through targeted gene inactivation and structural characterization of pathway intermediates, we delineate a tightly ordered sequence of oxidative transformations and revise a homology-based assignment. Our data demonstrate that PerV-dependent ether formation gates entry into macrocyclization, that PerT (rather than PerU) catalyses indole *N*-hydroxylation, and that oxidative tailoring occurs largely after macrocycle assembly but prior to glycosylation. Together, these findings reveal a hierarchical biosynthetic logic in which macrocyclization establishes the conformational framework required for downstream oxidative diversification, sugar installation, and late-stage PerA-mediated maturation.

Biological evaluation of pathway intermediates demonstrates that macrocycle formation is essential for antimycobacterial activity, underscoring its role in ribosomal engagement. Notably, the *ΩperX*-derived intermediate **7** displays an approximately 21-fold increase in potency against *M. tuberculosis* H37Rv relative to persiathiacin A, identifying the second dehydroalanine residue as an important determinant of antimycobacterial activity. This was associated with increased activity of intermediate **7** in time-kill assays relative to the other compounds. Although this potency advantage is not retained in an intracellular macrophage infection model, the discovery of a biosynthetic intermediate with enhanced *in vitro* activity highlights the potential of pathway interrogation to reveal cryptic analogues with altered biological properties. Importantly, intermediate **7** contains a carboxylic acid–bearing moiety within the macrocyclic tail that provides a versatile handle for chemical diversification. Because modifications to this region of thiopeptides are often well tolerated, this functional group offers a promising entry point for semisynthetic derivatisation aimed at improving physicochemical and pharmacological properties.

Collectively, this work provides a comprehensive and experimentally validated blueprint for persiathiacin biosynthesis. By elucidating the functional logic and temporal choreography of oxidative tailoring in a complex thiopeptide pathway, our study establishes a foundation for rational pathway engineering and late-stage diversification toward next-generation thiopeptide antibiotics with improved antibacterial potency and drug-like properties.

## Acknowledgements

This work was supported by an ARC Australian Laureate Fellowship (grant number FL210100071) to JE and NHMRC grant (APP2038610) to WJB. YD acknowledges support from the Sydney Infectious Diseases Institute. FAS is supported by the DOST-SEI Foreign Graduate Scholarship Program. We thank Biswaranjan Mohanty (Sydney Analytical) and Sam Lee (Sydney Mass Spectrometry) for NMR and mass spectrometry support.

## Supporting Information

Supporting Information Available free of charge: Materials and methods; primers used in this study; NMR and MS data and.

