## Supplementary Information for "Genetic dissection of persiathiacin biosynthesis defines hierarchical P450 oxidations and reveals a more potent antitubercular intermediate"

| **Contents** | **Page** |
|---|---|
| Materials and Methods | S2-S7 |
| Table S1. Sequences of primers used for constructing vectors. | S8 |
| Table S2. Sequences of primers used to validate the insertion site. | S8 |
| Figure S1. Chemical structure of compound **7** with numbering as shown in Table S3. | S9 |
| Table S3. Chemical shifts and HMBC correlations of **7**. | S9-S10 |
| Figure S2. Chemical structure of compound **9** with numbering as shown in Table S3. | S11 |
| Table S4. Chemical shifts and HMBC correlations of **9**. | S11-S12 |
| Figure S3. Chemical structure of compound **10** with numbering as shown in Table S3. | S13 |
| Table S5. Chemical shifts and HMBC correlations of **10**. | S13-S15 |
| Figure S4. Chemical structure of compound **12** with numbering as shown in Table S3. | S15 |
| Table S6. Chemical shifts and HMBC correlations of **12**. | S16-S17 |
| Figure S5. Comparative LC-MS profiles of extracts from wild-type *Actinokineospora* sp. UTMC 2448 and the *Actinokineospora* sp. UTMC 2448 *ΩperS5* mutant. | S18 |
| Figure S6. Time–kill kinetics of rifampicin and isoniazid against *M. tuberculosis*. | S18 |
| Figures S7-S14. High resolution mass spectrometry profiles of compounds **7-14**. | S19-S21 |
| Figure S15-S19: 1D/2D NMR spectra of **7**. | S22-S26 |
| Figure S20-S24: 1D/2D NMR spectra of **9**. | S27-S31 |
| Figure S25-S29: 1D/2D NMR spectra of **10**. | S32-S36 |
| Figure S30-S34: 1D/2D NMR spectra of **12**. | S37-S41 |
| Figure S35-S38: 1D/2D NMR spectra of **13**. | S42-S45 |
| Figure S39-S42: 1D/2D NMR spectra of **14**. | S46-S49 |

### Materials and Methods

**General Experimental Procedures:** LC-HR-MS analysis was performed on a Thermo Vanquish UHPLC system equipped with a Proshell 120 EC-C18 column (2.1 × 100 mm, 1.9 μm) and connected to an Orbitrap IQ-X Tribrid mass spectrometer. An Agilent 1260 Infinity II preparative HPLC, fitted with a semi-preparative reverse-phase C18 Betasil column (21.2 × 150 mm), was used for the purification of persiathiacin analogues. NMR spectra were recorded on a Bruker 600 or 800 MHz spectrometer equipped with a TCI cryoprobe at 25 °C. Compounds were dissolved in a mixture of $CDCl_3$-$CD_3OD$ (9:1) or DMSO-$d_6$ for NMR spectroscopic analyses. The $^1$H and $^{13}$C NMR chemical shifts were referenced to the solvent peaks at $\delta_H$ 7.26 and $\delta_C$ 77.16 for $CDCl_3$ or to the DMSO-$d_6$ solvent peaks at $\delta_H$ 2.50 and $\delta_C$ 39.52. All HPLC and LC-MS experiments were performed with a MeCN-$H_2O$ gradient solvent system. Millipore Milli-Q $H_2O$ and HPLC grade solvents were used for chromatography.

**Insertional Mutagenesis by Protoplast Transformation:** Insertional mutagenesis by protoplast transformation was used to inactivate target genes in *Actinokineospora* sp. UTMC 2248. The pSET-*kasOp*-perX* vector was initially constructed by assembling three DNA fragments: *(i)* a 3.5 kb fragment from pSET152 containing the origin of replication, apramycin resistance cassette, and *traJ* gene; *(ii)* a 2 kb internal fragment of *perX*; and *(iii)* the *kasOp** promoter,[1] inserted upstream of the amplified *perX* fragment to minimise polar effects on downstream genes within the *per* BGC following insertional mutagenesis. The three fragments were assembled using the NEBuilder system according to the manufacturer's instructions.

The resulting pSET-*kasOp*-perX* construct was subsequently used as the backbone for generation of additional mutagenesis vectors. These constructs were generated by two-piece assembly consisting of: *(i)* a linearised pSET-*kasO*p* backbone derived from pSET-*kasO*p*-*perX*, and (ii) an internal fragment of the target gene amplified with 20 bp overlaps matching the pSET-*kasO*p* backbone. Primer pairs used for amplification are listed in Table S1. All resulting pSET-based constructs were verified by DNA sequencing.

Protoplast preparation and buffer compositions were adapted from MacNeil and Klapko.[2] *Actinokineospora* sp. UTMC 2248 was cultured in YEME medium (3 g/L yeast extract, 3 g/L malt extract, 5 g/L peptone, and 10 g/L glucose) supplemented with 10% (w/v) sucrose to promote dispersed growth, and incubated at 30 °C for 2 days. The culture was passed through a 70 µm nylon cell strainer to remove large aggregates. The filtrate was centrifuged and the pellet washed twice with 0.3 M sucrose.

Cells were resuspended in 7 mL of P buffer containing 2 mg/mL lysozyme and incubated at 30 °C for 15 min to generate protoplasts. P buffer (50 mL) consisted of 40 mL sucrose solution (103 g/L sucrose, 0.25 g/L $K_2SO_4$, and 10.12 g/L $MgCl_2 \cdot 6H_2O$), 5 mL $CaCl_2 \cdot 2H_2O$ solution, 5 mL TES buffer (2-[Tris(hydroxymethyl)methylamino]ethanesulfonic acid), and 500 µL 0.5% (w/v) $KH_2PO_4$. Protoplast formation was confirmed microscopically using 5 µL of the suspension. Protoplasts were subsequently collected by centrifugation at 2000 × g for 15 min and resuspended in 1 mL P buffer. Prepared protoplasts could be stored at 4 °C for up to 1–2 weeks without noticeable loss of quality.

For transformation, protoplasts were diluted 1:10 in P buffer and mixed with 2–4 µg plasmid DNA, followed by addition of 200 µL transformation buffer. The transformation mixture was spread onto two pre-dried R2YE agar plates and incubated at 30 °C. After overnight incubation, one plate from each transformation was overlaid with 1 mL sterile deionised water containing apramycin to a final concentration of 30 µg/mL, while the second plate was left untreated to assess protoplast regeneration efficiency.

Transformation plates were incubated at 30 °C for 5–10 days. Colonies obtained from apramycin-overlaid plates were streaked onto R2YE agar supplemented with 30 µg/mL apramycin, followed by re-streaking onto YEME agar containing 50 µg/mL apramycin. Resistant colonies were subsequently cultured in liquid YEME supplemented with apramycin at 30 °C for 2 days. Genomic DNA was extracted using the Wizard® Genomic DNA Purification Kit (Promega) according to the manufacturer's instructions.

Chromosomal insertion of the pSET-based constructs was verified by PCR using primer pairs targeting the left and right flanking regions of the target gene. In each case, one primer annealed outside the target gene region and the second primer annealed within the pSET152 backbone. Primer pairs used for verification of insertion mutants are listed in Table S2. To assess the effect of insertional mutagenesis on persiathiacin production, mutant and wild-type strains were cultivated on solid ISP2 medium (4 g/L glucose, 4 g/L yeast extract, 10 g/L malt extract, 2 g/L $CaCO_3$, 15 g/L Bacto agar) for 7 days. Agar cultures were cut into small pieces, extracted with EtOAc, and analysed by UHPLC–HRMS.

**Large Scale Production, Extraction, and HPLC Purification of the Metabolites:** Wild type *Actinokineospora* sp. UTMC 2448 and generated mutants were grown on 1- or 2 L solid ISP2 medium for 7 days at 30 °C. The agar cultures were chopped and extracted with EtOAc. The extract was dried on a rotary evaporator and Genevac, then resuspended in 900 µL DMSO for direct injection to HPLC using reverse-phase C18 column. The column was eluted with 5% acetonitrile for 5 min, then a linear gradient from 5 to 100% acetonitrile was applied over 45 min, and the column was eluted for an additional 10 min with 100% acetonitrile. The flow rate was 12 mL/min. Sixty fractions were collected in 1 min increments over 60 min. Compound **7** was mainly eluted in fractions 36-37. This intermediate was distributed in many fractions till last fraction of the HPLC. Compounds **9**, **10**, **12**, **13**, and **14** were eluted in fractions 35, 38, 39, 39, 37, respectively (Note that these compounds obtained from separate HPLC runs from separate extracts).

*(7)*: $^1$H NMR (600 MHz, $CDCl_3$:$CD_3OD$ (9:1)) and $^{13}$C NMR (150 MHz, $CDCl_3$:$CD_3OD$ (9:1)), see Table S3; HRESIMS *m*/*z* 1336.1685 $[M + H]^+$ (calcd for $C_{55}H_{46}N_{13}O_{18}S_5$, 1336.1682).

*(8)*: HRESIMS *m*/*z* 1510.2576 $[M + H]^+$ (calcd for $C_{63}H_{60}N_{13}O_{22}S_5$, 1510.2574).

*(9)*: $^1$H NMR (600 MHz, DMSO-$d_6$) and $^{13}$C NMR (150 MHz, DMSO-$d_6$), see Table S4; HRESIMS *m*/*z* 1280.2147 $[M + H]^+$ (calcd for $C_{54}H_{50}N_{13}O_{15}S_5$, 1280.2147).

*(10)*: $^1$H NMR (600 MHz, $CDCl_3$:$CD_3OD$ (9:1)) and $^{13}$C NMR (150 MHz, $CDCl_3$:$CD_3OD$ (9:1)), see Table S5; HRESIMS *m*/*z* 1858.4724 $[M + H]^+$ (calcd for $C_{80}H_{92}N_{13}O_{29}S_5$, 1858.4722).

*(11)*: HRESIMS *m*/*z* 1844.4565 $[M + H]^+$ (calcd for $C_{79}H_{90}N_{13}O_{29}S_5$, 1844.4565).

*(12)*: $^1$H NMR (600 MHz, $CDCl_3$:$CD_3OD$ (9:1)) and $^{13}$C NMR (150 MHz, $CDCl_3$:$CD_3OD$ (9:1)), see Table S6; HRESIMS *m*/*z* 1858.4723 $[M + H]^+$ (calcd for $C_{80}H_{92}N_{13}O_{29}S_5$, 1858.4722).

*(13)*: HRESIMS *m*/*z* 1282.1566 $[M + H]^+$ (calcd for $C_{52}H_{44}N_{13}O_{17}S_5$, 1282.1576).

*(14)*: HRESIMS *m*/*z* 1456.2458 $[M + H]^+$ (calcd for $C_{60}H_{58}N_{13}O_{21}S_5$, 1456.2468).

**Resazurin assay:** *Mycobacterium tuberculosis* H37Rv was cultured in Middlebrook 7H9 media (Becton Dickinson, Franklin Lakes, USA), supplemented with 0.2% v/v glycerol (Sigma Aldrich, Burlington, USA), 5 g/L bovine serum albumin (BSA; Moregate Biotech, Bulimba, Australia), 2 g/L D-glucose (Sigma Aldrich), 4 mg/L catalase (Sigma Aldrich), and 0.02% v/v tyloxapol (Sigma Aldrich). Cultures were passaged twice weekly, maintaining $OD_{600}$ below 0.8. To assay compound activity, *M. tuberculosis* H37Rv was cultured to approximately $OD_{600}$ = 0.5 in 7H9 medium in the logarithmic growth phase. From this culture, 7H9 medium was inoculated at $OD_{600}$ = 0.005 in flat-bottom 96-well plates, 200 µl/well. Test compounds were titrated in 7H9 medium and added directly to plated cultures, then incubated at 37 °C. After 7 days, 10 µL resazurin solution (0.025% w/v, Sigma Aldrich) was added to each well (0.00125% w/v final concentration) and incubated at 37 °C for 24 h. Luminescence was read using a POLARstar Omega instrument (BMG Labtech); excitation at 550 nm, emission at 590 nm, bottom read.

**Time-kill assay:** The luminescent *M. tuberculosis* H37Rv lux strain was used as previously published.[3] H37Rv lux was cultured in 7H9 medium with 50 µg/ml kanamycin (Sigma Aldrich) to approximately $OD_{600}$ = 0.5, within the logarithmic growth phase. The culture was centrifuged at 6000 rcf for 5 min and resuspended in 7H9 medium containing 1% DMSO (7H9/DMSO). The bacterial suspension was sonicated for 30 s in a Grant XUBA water bath (44 kHz, 35 W), and diluted to 0.01 $OD_{600}$ in 7H9/DMSO. Compounds were diluted to 2X concentrations in 7H9/DMSO at concentrations chosen to span a full dose response for each compound. In clear-bottom flat white 96-well plates (Greiner Bio-One, Kremsmünster, Austria), 100 µL of diluted compounds was added to 100 µL of diluted bacteria; 1X drug and 0.005 $OD_{600}$ bacteria final concentrations. Edge wells were not used, and were filled with sterile water. Plates were incubated in a humidified 37 °C incubator for 28 days and luminescence read at regular intervals. Luminescence was measured using a POLARstar Omega plate reader, bottom read, 5 second exposure.

**Intracellular assay:** Human THP-1 cells (ATCC TIB-202) were cultured in RPMI-1640 medium (Gibco, Waltham, USA) with 10% v/v foetal bovine serum (HyClone, Marlborough, USA) and 50 µM 2-mercaptoethanol (Sigma Aldrich). Cultures were passaged twice weekly, maintained between 2 x $10^5$ and 1 x $10^6$ cells/ml.

To examine the effect of test compounds during intracellular *M. tuberculosis* growth, THP-1 cells were seeded in clear-bottom flat white 96-well plates, 6 x $10^4$ cells in 200 µl. Cells were incubated overnight in a 37 °C humidified incubator, 5% $CO_2$. During this incubation, 100 ng/mL phorbol 12-myristate 13-acetate (Sigma Aldrich) was added to THP-1 monocytes to differentiate them to macrophages. Compounds were diluted in RPMI medium supplemented with 0.1% DMSO (RPMI/DMSO). H37Rv *lux* culture in logarithmic phase was washed, suspended in RPMI medium, and sonicated for 30 s as above. Culture media was removed from macrophage cells and H37Rv *lux* added at a multiplicity of infection of 1. Plates were incubated for 4 h at 37 °C before wells were washed 3 times with RPMI media, and diluted compounds were added in RPMI/DMSO. Plates were incubated for 5 days post-infection. Luminescence was measured using a POLARstar Omega plate reader, bottom read, 5 second exposure.

**Table S1.** Sequences of primers used for construction vectors used for insertional mutagenesis.

| Construct/DNA fragment | Primer Name | Sequences |
|---|---|---|
| pSET-*kasOp**-*perX* | UTMC-*perX*-F | GACCTGCAGGCATGCAAGCTGAGTTCTACCGCGACCCGCA |
| | UTMC-*perX*-R | CAGGCTTCCCGGGTGTCTCGCCGGAAGAACGACCGGAACG |
| | pSET-perX-F | CGTTCCGGTCGTTCTTCCGGCGAGACACCCGGGAAGCCTG |
| | pSET-perX-R | TGCGGGTCGCGGTAGAACTCAGCTTGCATGCCTGCAGGTC |
| | *kasOp**-F | CTAGTTGTTCACATTC |
| | *kasOp**-R | TATGAACTCCCCCAG |
| Blunt end for pSET-*kasOp**[a] | pSET-F | CGAGACACCCGGGAAGCCTG |
| | *kasOp**-pSET-R | CGATTATGAACTCCCCCAGTCCT |
| pSET-*kasOp**-*perB* | *kasOp**-*perB*-F | ACTGGGGGAGTTCATAATCGAACCGGCGCACCGTCCGGCC |
| | UMTC-*perB*-R | CAGGCTTCCCGGGTGTCTCGCGGTCGCCCCCGTCACCCTG |
| pSET-*kasOp**-*perU* | *kasOp**-*perU*-F | ACTGGGGGAGTTCATAATCGGGCCTGCTGCCCGATCCGTA |
| | UMTC-*perU*-R | CAGGCTTCCCGGGTGTCTCGGCGGACAGCATGTGCCCCAG |
| pSET-*kasOp**-*perT* | *kasOp**-*perT*-F | ACTGGGGGAGTTCATAATCGACCGGCCAGTGGGTCGTCCG |
| | UMTC-*perT*-R | CAGGCTTCCCGGGTGTCTCGCCCGCAGCACCTGCGAGAAG |
| pSET-*kasOp**-*perC* | *kasOp**-*perC*-F | ACTGGGGGAGTTCATAATCGCGTTCGCCATGCCCGAGGAG |
| | UMTC-*perC*-R | CAGGCTTCCCGGGTGTCTCGGCCCTCCTGCATCTCGGCCC |
| pSET-*kasOp**-*perV* | *kasOp**-*perV*-F | ACTGGGGGAGTTCATAATCGGCGCTGCGCGACCCGCACAT |
| | UMTC-*perV*-R | CAGGCTTCCCGGGTGTCTCGTGTGCGTGTCGATCCGCACG |
| pSET-*kasOp**-*pers4* | *kasOp**-*perS4*-F | ACTGGGGGAGTTCATAATCGCCGCGGCGGCGGGGCACGAC |
| | UMTC-*perS4*-R | CAGGCTTCCCGGGTGTCTCGCGCAGTGGCCTAGCGTACGC |
| pSET-*kasOp**-*pers5* | *kasOp**-*perS5*-F | ACTGGGGGAGTTCATAATCGGAGTACGCCGAGTCCGTCGC |
| | UMTC-*perS5*-R | CAGGCTTCCCGGGTGTCTCG GCGACGGACTCGGCGTACTC |

[a] pSET-*kasOp**-*perX* construct used to a template to generate following other constructs

**Table S2.** Sequences of primers used to validate the insertion site within the genome of *Actinokineospora* sp. UTMC 2448.

| Target gene | Primer Name | Sequences |
|---|---|---|
| Primers on the pSET152 backbone | check-1R | CTCTAGAGTCGACCTGCAGC |
| | check-2F | GCCAGGTGCGAATAAGGGAC |
| *perX* | check-*perX*-1F | AAGTTCTACGCCACC |
| | check-*perX*-2R | TTGTCGACGATCGCC |
| *perB* | check-*perB*-1F | GAAGAAGCAATTGGC |
| | check-*perB*-2R | CACCCATGCTCGAAT |
| *perU* | check-*perU*-1F | AGTGGGAGATCGACA |
| | check-*perU*-2R | GCTCGAAGAACTGCG |
| *perT* | check-*perT*-1F | GCTGTTCGAGGAGTT |
| | check-*perT*-2R | GGACATAGACGGCGT |
| *perC* | check-*perC*-1F | TTCCTCGGCTTCCTC |
| | check-*perC*-2R | TGATGATGTGCGGGT |
| *perS4* | check-*perS4*-1F | CGTGATCATCGACGA |
| | check-*peS4*-2R | GCAGCCGATGAGGAT |
| *perS5* | check-*perS5*-1F | TTCGAGCGGAACTAC |
| | check-*perS5*-2R | CGATGTGGGCGAAGC |

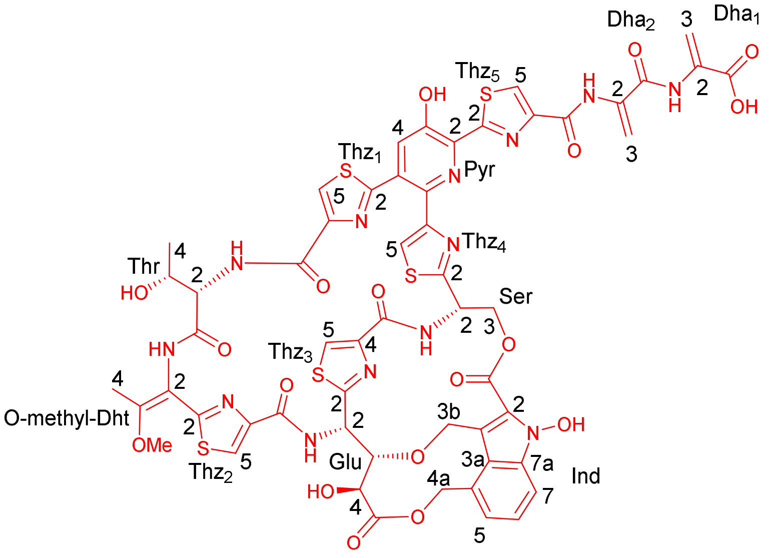


**7**

**Figure S1.** Chemical structure of compound **7** with numbering as shown in Table S3.

**Table S3.** $^{1}H$ (600 MHz) and $^{13}C$ (150 MHz) chemical shifts and HMBC correlations of **7** recorded in $CDCl_3$-$CD_3OD$ (9:1).

| Position | | $\delta_C$ | $\delta_H$ | HMBC |
|---|---|---|---|---|
| Dha1 | C=O | 169.3 | | |
| | C2 | 135.6 | | |
| | C3 | 104.9 | H3a: 5.78, brs<br>H3b: 6.32, brs | Dha1-(C=O)<br>Dha1-(C2, C=O) |
| | NH | | Exchanged/not observed | |
| Dha2 | C=O | 161.3 | | |
| | C2 | 133.8 | | |
| | C3 | 103.0 | H3a: 5.57, brs<br>H3b: 6.59, brs | Dha2-(C=O)<br>Dha2-(C2, C=O) |
| | NH | | Exchanged/not observed | |
| Thz5 | C2 | 166.4 | | |
| | C4 | 149.3 | | |
| | C5 | 124.6 | 8.20, s | Thz5-(C2, C4, C=O) |
| | C=O | 160.5 | | |
| Pyr | C2 | 134.7 | | |
| | C3 | 151.4 | | |
| | C4 | 126.9 | 7.66, s | Pyr-(C2, C3, C6), Thz-(C2) |
| | C5 | 128.5 | | |
| | C6 | 144.0 | | |
| Thz1 | C2 | 165.5 | | |
| | C4 | 149.2 | | |
| | C5 | 125.5 | 8.29, s | Thz1-(C2, C4, C=O) |
| | C=O | 161.5 | | |
| Thr | NH | | Exchanged/not observed | |
| | C=O | 167.2 | | |
| | C2 | 56.2 | 4.26, d, $J$ = 4.2 | Thr-(C3, C4, C=O), Thz1-(C=O) |
| | C3 | 64.4 | 2.9, m | Thr-(C=O) |
| | C4 | 18.1 | 1.43, d, $J$ = 6.0 | Thr-(C2, C3) |

| | | | | |
|---|---|---|---|---|
| O-methyl-Dht | NH | | Exchanged/not observed | |
| | C2 | 110.8 | | |
| | C3 | 159.0 | | |
| | C4 | 13.7 | 1.87, s | Dht-(C2, C3), Thz2-(C2) |
| | OMe | 55.8 | 3.80, s | Dht-(C3) |
| Thz2 | C2 | 162.1 | | |
| | C4 | 145.8 | | |
| | C5 | 124.9 | 7.97, s | Thz2-(C2, C4, C=O) |
| | C=O | 161.2 | | |
| Glu | NH | | 8.32, d, *J* = 9.8 | Glu-(C2), Thz2-(C=O) |
| | C2 | 48.9 | 6.03, d, *J* = 9.8 | Glu-(C3) |
| | C3 | 83.4 | 3.67, d, 11.4 | Glu-(C4), Ind (C3b) |
| | C4 | 67.6 | 4.17, *ol* | Glu-(C2, C3, C=O) |
| | C=O | 174.1 | | |
| Thz3 | C2 | 169.3 | | |
| | C4 | 150.6 | | |
| | C5 | 126.6 | 8.28, s | Thz3-(C2, C4) |
| | C=O | 160.5 | | |
| Ser | NH | | 8.12, d, *J* = 11.0 | Thz3-(C=O) |
| | C2 | 48.4 | 5.68, dd, *J* = 5.9, 11.0 | Thz3-(C=O), Thz4-(C2) |
| | C3 | 64.4 | H3a: 4.42, d, *J* = 11.4<br>H3b: 5.29, dd, *J* = 5.9, 11.4 | Thz4-(C2), Ind-(C=O)<br>Ser-(C2), Thz4-(C2), Ind-(C=O) |
| Thz4 | C2 | 169.1 | | |
| | C4 | 155.0 | | |
| | C5 | 120.3 | 7.67, s | Thz4-(C2, C4), Pyr-(C6) |
| Ind | C=O | 161.3 | | |
| | C2 | 127.0 | | |
| | C3 | 109.8 | | |
| | C3a | 119.3 | | |
| | C3b | 65.9 | H3ba: 4.19, *ol*<br>H3bb: 4.86, d, *J* = 10.8 | Ind-(C2, C3, C3a), Glu-(C3)<br>Ind-(C2, C3, C3a) |
| | C4 | 127.9 | | |
| | C4a | 68.5 | H4aa: 4.99, d, *J* = 12.5<br>H4ab: 5.95, d, *J* = 12.5 | Ind-(C3a, C4, C5)<br>Ind-(C3a, C4, C5), Glu-(C=O) |
| | C5 | 123.3 | 7.13, d, *J* = 7.0 | Ind-(C3a, C4a, C7) |
| | C6 | 125.3 | 7.38, dd, *J* = 6.8, 8.4 | Ind-(C4, C7a) |
| | C7 | 112.3 | 7.79, d, *J* = 8.5 | Ind-(C3a, C5) |
| | C7a | 135.5 | | |
| | N-OH | | Exchanged/not observed | |

* overlapped

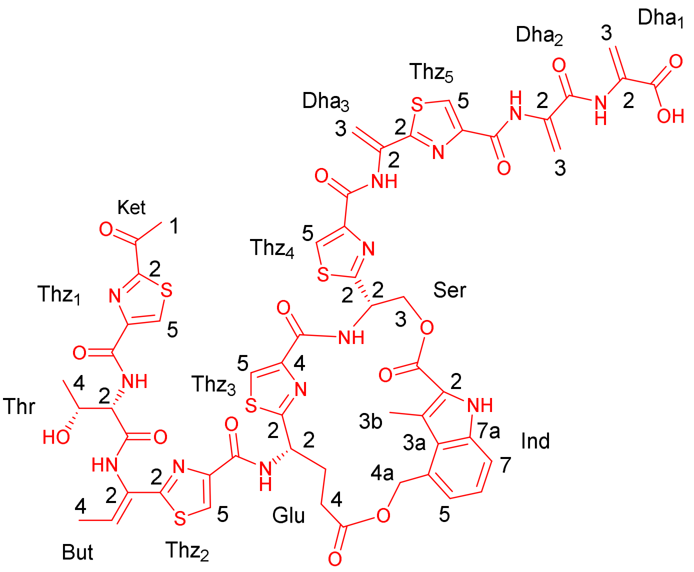

**9**

**Figure S2.** Chemical structure of compound **9** with numbering as shown in Table S4.

**Table S4.** $^{1}$H (600 MHz) and $^{13}$C (125 MHz) chemical shifts and HMBC correlations of **9** recorded in DMSO-$d_6$.

| Position | | $\delta_C$ | $\delta_H$ | HMBC |
|---|---|---|---|---|
| Dha1 | C=O | 164.6 | | |
| | C2 | 134.7 | | |
| | C3 | 104.0 | H3a: 5.74, brs<br>H3b: 5.98, brs | Dha1-(C=O)<br>Dha1-(C2, C=O) |
| | NH | | 9.67, s | Dha1-(C=O), Dha2-(C=O) |
| Dha2 | C=O | 162.0 | | |
| | C2 | 134.3 | | |
| | C3 | 104.4 | H3a: 5.79, *ol**<br>H3b: 6.59, d, $J$ = 1.5 | Dha2-(C2, C=O)<br>Dha2-(C2, C=O) |
| | NH | | 9.88, s | Dha2-(C3, C=O), Thz5-(C4, C=O) |
| Thz5 | C2 | 165.3 | | |
| | C4 | 148.5 | | |
| | C5 | 127.2 | 8.54, s | Thz5-(C2, C4, C=O) |
| | C=O | 158.6 | | |
| Dha3 | C2 | 133.5 | | |
| | C3 | 104.5 | H3a: 5.79, *ol*<br>H3b: 6.44, d, $J$ = 1.6 | Dha3-(C2), Thz5-(C2)<br>Dha3-(C2), Thz5-(C2) |
| | NH | | 10.31, s | Dha3-(C3), Thz4-(C4), Thz5-(C2) |
| Ket | C1 | 25.8 | 2.71, s | Ket (C=O), Thz1-(C2) |
| | C=O | 191.2 | | |
| Thz1 | C2 | 166.6 | | |
| | C4 | 150.7 | | |
| | C5 | 131.4 | 8.76, s | Thz1-(C2, C4, C=O) |
| | C=O | 160.1 | | |
| Thr | NH | | 8.17, d, $J$ = 8.4 | Thz1-(C4, C=O), Thr-(C2, C3, C=O) |
| | C=O | 169.5 | | |
| | C2 | 58.6 | 4.59, d, $J$ = 3.7, 8.4 | Thr-(C3, C4, C=O), Thz1-(C=O) |

| | | | | |
|---|---|---|---|---|
| | C3 | 66.5 | 4.33, m | Thr-(C=O) |
| | C4 | 20.8 | 1.22, d, *J* = 6.3 | Thr-(C2, C3) |
| But | NH | | 9.80, s | But-(C2, C3), Thr-(C=O), Thz2-(C2) |
| | C2 | 128.6 | | |
| | C3 | 126.4 | 6.81, q, *J* = 7.1 | But-(C2, C4), Thz2-(C2) |
| | C4 | 13.3 | 1.77, d, *J* = 7.1 | But-(C2, C3), Thz2-(C2) |
| Thz2 | C2 | 167.6 | | |
| | C4 | 149.1 | | |
| | C5 | 124.6 | 8.28, s | Thz2-(C2, C4, C=O) |
| | C=O | 161.0 | | |
| Glu | NH | | 9.24, d, *J* = 8.7 | Thz2-(C=O), Glu-(C2, C3) |
| | C2 | 50.9 | 5.37, m | Glu-(C3, C4), Thz3-(C2) |
| | C3 | 29.1 | 2.45, m<br>2.81, m | Glu-(C2, C=O) |
| | C4 | 31.5 | 2.41, m | Glu-(C2, C3, C=O) |
| | C=O | 171.6 | | |
| Thz3 | C2 | 174.3 | | |
| | C4 | 148.0 | | |
| | C5 | 125.6 | 8.39, s | Thz3-(C2, C4, C=O) |
| | C=O | 160.1 | | |
| Ser | NH | | 8.72, d, *J* = 8.2 | Ser-(C2, C3), Thz3-(C=O) |
| | C2 | 50.3 | 6.00, m | Ser-(C3), Thz3-(C=O), Thz4-(C2) |
| | C3 | 66.5 | H3a: 4.65, dd, *J* = 3.6, 11.3<br>H3b: 5.15, dd, *J* = 2.6, 11.2 | Ser-(C2), Thz4-(C2), Ind-(C=O)<br>Thz4-(C2), Ind-(C=O) |
| Thz4 | C2 | 172.4 | | |
| | C4 | 149.1 | | |
| | C5 | 125.8 | 8.42, s | Thz4-(C2, C4, C=O) |
| | C=O | 158.9 | | |
| Ind | C=O | 161.9 | | |
| | C2 | 123.9 | | |
| | C3 | 118.1 | | |
| | C3a | 125.6 | | |
| | C3b | 12.6 | 2.59, s | Ind-(C2, C3, C3a) |
| | C4 | 129.3 | | |
| | C4a | 65.9 | 5.39, m | Ind-(C3a, C4, C5), Glu-(C=O) |
| | C5 | 123.1 | 7.06, d, *J* = 7.1 | Ind-(C3a, C4a, C7) |
| | C6 | 124.3 | 7.18, dd, *J* = 7.0, 8.4 | Ind-(C4, C5, C7a) |
| | C7 | 113.9 | 7.41, d, *J* = 8.3 | Ind-(C3a, C5) |
| | C7a | 137.5 | | |
| | N-H | | 11.79, s | Ind-(C2, C3, C3a, C7a) |

* overlapped

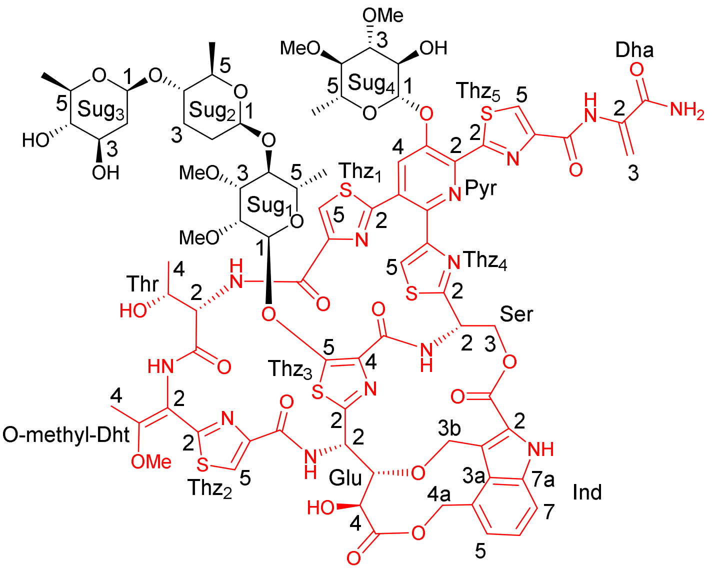


**10**

**Figure S3.** Chemical structure of compound **10** with numbering as shown in Table S5.

**Table S5.** $^{1}H$ (600 MHz) and $^{13}C$ (150 MHz) chemical shifts and HMBC correlations of **10** recorded in $CDCl_3$-$CD_3OD$ (9:1).

| Position | | $\delta_C$ | $\delta_H$ | HMBC |
|---|---|---|---|---|
| Dha | C=O | 166.5 | | |
| | C2 | 133.5 | | |
| | C3 | 104.9 | H3a: 5.55, brs<br>H3b: 6.53, brs | Dha-( C2, C=O)<br>Dha-(C2, C=O) |
| | NH | | 9.88, s | Dha-(C3, C=O), Thz5-(C=O) |
| Thz5 | C2 | 166.6 | | |
| | C4 | 151.1 | | |
| | C5 | 127.3 | 8.21, s | Thz5-(C2, C4, C=O) |
| | C=O | 159.7 | | |
| Pyr | C2 | 139.5 | | |
| | C3 | 149.3 | | |
| | C4 | 126.9 | 7.92, s | Pyr-(C2, C3, C6), Thz1-(C2) |
| | C5 | 128.8 | | |
| | C6 | 144.2 | | |
| Thz1 | C2 | 164.5 | | |
| | C4 | 150.4 | | |
| | C5 | 124.2 | 8.35, s | Thz1-(C2, C4, C=O) |
| | C=O | 159.4 | | |
| Thr | NH | | 7.39, *ol** | Thz1-(C=O), Thr-(C=O) |
| | C=O | 168.0 | | |
| | C2 | 55.5 | 4.26, *ol* | Thr-(C3, C=O), Thz1-(C=O) |
| | C3 | 63.6 | 2.02, m | Thr-(C=O) |
| | C4 | 17.7 | 1.29, d, *J* = 6.0 | Thr-(C2, C3) |
| O-methyl-Dht | NH | | 7.39, *ol* | Dht-(C3), Thr-(C=O) |
| | C2 | 110.6 | | |
| | C3 | 158.6 | | |
| | C4 | 13.7 | 1.92, s | Dht-(C2, C3), Thz2-(C2) |
| | OMe | 55.5 | 3.82, s | Dht-(C3) |

| | | | | |
|---|---|---|---|---|
| Thz2 | C2 | 161.8 | | |
| | C4 | 146.0 | | |
| | C5 | 124.3 | 8.00, s | Thz2-(C2, C4, C=O), Dht-(C2) |
| | C=O | 161.6 | | |
| Glu | NH | | 8.27, d, *J* = 9.4 | Glu-(C2), Thz2-(C=O) |
| | C2 | 49.3 | 5.83, d, *J* = 10.0 | Glu-(C3), Thz2-(C=O), Thz3-(C2) |
| | C3 | 82.2 | 3.68, *ol* | Glu-(C4, C=O), Thz3-(C2), Ind (C3b) |
| | C4 | 67.8 | 4.22, d, *J* = 9.5 | Glu-(C2, C3, C=O) |
| | C=O | 174.7 | | |
| Thz3 | C2 | 154.0 | | |
| | C4 | 131.1 | | |
| | C5 | 160.4 | | |
| | C=O | 161.2 | | |
| Ser | NH | | 7.69, *ol* | Ser-(C2), Thz3-(C=O) |
| | C2 | 50.6 | 5.73, m | |
| | C3 | 62.8 | H3a: 4.62, d, *J* = 11.5<br>H3b: 5.33, dd, *J* = 4.7, 11.7 | Thz4-(C2), Ind-(C=O)<br>Ser-(C2), Thz4-(C2), Ind-(C=O) |
| Thz4 | C2 | 169.1 | | |
| | C4 | 154.6 | | |
| | C5 | 120.5 | 7.68, s | Thz4-(C2, C4), Pyr-(C6) |
| Ind | C=O | 161.4 | | |
| | C2 | 126.5 | | |
| | C3 | 116.5 | | |
| | C3a | 123.9 | | |
| | C3b | 64.5 | H3ba: 4.19, d, *J* = 10.1<br>H3bb: 5.24, *ol* | Ind-(C2, C3, C3a), Glu-(C3)<br>Ind-(C2, C3, C3a) |
| | C4 | 127.3 | | |
| | C4a | 68.5 | H4aa: 5.01, d, *J* = 12.3<br>H4ab: 6.05, d, *J* = 12.3 | Ind-(C3a, C4, C5), Glu-(C=O)<br>Ind-(C3a, C4, C5), Glu-(C=O) |
| | C5 | 123.5 | 7.15, d, *J* = 7.1 | Ind-(C3a, C4a, C6, C7) |
| | C6 | 124.8 | 7.36, m | Ind-(C4, C5, C7, C7a) |
| | C7 | 116.0 | 7.74, d, *J* = 8.4 | Ind-(C3a, C5) |
| | C7a | 136.5 | | |
| | N-H | | 10.18, s | Ind-(C2, C3, C3a, C7a) |
| Sug1 | C1 | 101.7 | 5.48, brs | Thz3-(C5), Sug1-(C3, C5) |
| | C2 | 76.0 | 4.07, m | Sug1-(C1, C3, C4, C2-OMe) |
| | C2-OMe | 59.1 | 3.55, s | Sug1-(C2) |
| | C3 | 79.9 | 3.68, m | Sug1-(C1, C5, C3-OMe) |
| | C3-OMe | 57.8 | 3.48, s | Sug1-(C3) |
| | C4 | 76.8 | 3.70, m | Sug1-(C2, C5, C6), Sug2-(C1) |
| | C5 | 70.0 | 3.68, m | Sug1-(C1, C3, C4, C6) |
| | C6 | 17.6 | 1.35, d, *J* = 5.1 | Sug1-(C4, C5) |
| Sug2 | C1 | 102.1 | 4.67, dd, *J* = 2.1, 9.0 | Sug1-(C4), Sug2-(C2, C5) |
| | C2 | 30.9 | H2a: 1.46, m<br>H2b: 1.85, m | Sug2-(C1, C3, C4) |
| | C3 | 29.8 | H3a: 1.51, m<br>H3b: 2.15, m | Sug2-(C1, C4) |
| | C4 | 80.6 | 3.13, m | Sug2-(C5, C6), Sug3-(C1) |
| | C5 | 73.8 | 3.30, dd, *J* = 6.1, 9.1 | Sug2-(C1, C3, C6) |
| | C6 | 17.9 | 1.17, d, *J* = 6.1 | Sug2-(C4, C5) |
| Sug3 | C1 | 100.9 | 4.48, dd, *J* = 2.0, 9.8 | Sug2-(C4), Sug3-(C2, C3, C5) |

| | | | | |
|---|---|---|---|---|
| | C2 | 38.9 | H2a: 1.53, m<br>H2b: 2.13, m | Sug3-(C1, C3, C4) |
| | C3 | 71.2 | 3.51, *ol* | Sug3-(C1, C5) |
| | C4 | 77.0 | 2.97, t, 8.9 | Sug3-(C2, C5, C6) |
| | C5 | 71.6 | 3.20, dd, *J* = 6.2, 9.2 | Sug3-(C1, C4, C6) |
| | C6 | 17.5 | 1.26, d, *J* = 6.2 | Sug3-(C4, C5) |
| Sug4 | C1 | 100.8 | 5.24, *ol* | Pyr-(C3), Sug4-(C3, C5) |
| | C2 | 69.5 | 4.25, *ol* | Sug4-(C1, C3) |
| | C3 | 84.0 | 3.36, m | Sug4-(C2, C4, C3-OMe) |
| | C3-OMe | 58.2 | 3.56, s | Sug4-(C3) |
| | C4 | 77.6 | 3.53, m | Sug4-(C2, C4-OMe) |
| | C4-OMe | 61.7 | 3.60, s | Sug4-(C4) |
| | C5 | 71.2 | 3.81, m | Sug4-(C1, C3, C4, C6) |
| | C6 | 16.5 | 1.41, d, *J* = 6.4 | Sug4-(C4, C5) |

* overlapped

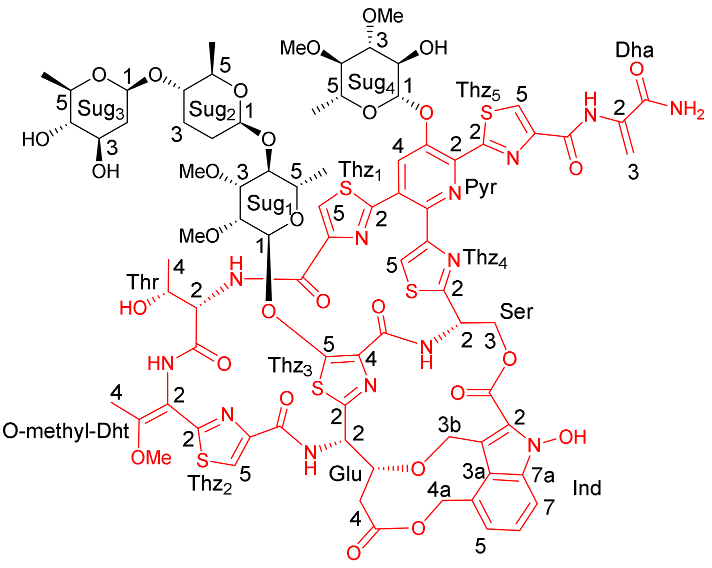


**12**

**Figure S4.** Chemical structure of compound **12** with numbering as shown in Table S6.

**Table S6.** $^{1}H$ (600 MHz) and $^{13}C$ (150 MHz) chemical shifts and HMBC correlations of **12** recorded in $CDCl_3$-$CD_3OD$ (9:1).

| Position | | $\delta_C$ | $\delta_H$ | HMBC |
|---|---|---|---|---|
| Dha | C=O | 167.0 | | |
| | C2 | 133.7 | | |
| | C3 | 104.9 | H3a: 5.6, brs<br>H3b: 6.5, brs | Dha-(C=O)<br>Dha-(C2, C=O) |
| | NH | | 9.94, s | Dha-(C3, C=O), Thz5-(C4, C=O) |
| Thz5 | C2 | 166.7 | | |
| | C4 | 151.3 | | |
| | C5 | 127.3 | 8.21, s | Thz5-(C2, C4, C=O) |
| | C=O | 160.1 | | |
| Pyr | C2 | 139.9 | | |
| | C3 | 149.1 | | |
| | C4 | 126.9 | 7.80, s | Pyr-(C2, C3, C6), Thz-(C2) |
| | C5 | 128.4 | | |
| | C6 | 145.0 | | |
| Thz1 | C2 | 165.6 | | |
| | C4 | 149.2 | | |
| | C5 | 125.1 | 8.28, s | Thz1-(C2, C4, C=O) |
| | C=O | 161.8 | | |
| Thr | NH | | 7.96, brs | Thz1-(C=O) |
| | C=O | 167.5 | | |
| | C2 | 56.4 | 4.29, *ol** | Thr-(C3, C4, C=O), Thz1-(C=O) |
| | C3 | 64.9 | 2.99, m | Thr-(C=O) |
| | C4 | 17.9 | 1.40, d, *J* = 6.5 | Thr-(C2, C3) |
| O-methyl-Dht | NH | | 8.401, s | Dht-(C3), Thr-(C=O) |
| | C2 | 111.0 | | |
| | C3 | 158.6 | | |
| | C4 | 13.8 | 1.90, s | Dht-(C2, C3), Thz2-(C2) |
| | OMe | 55.9 | 3.82, s | Dht-(C3) |
| Thz2 | C2 | 161.9 | | |
| | C4 | 146.1 | | |
| | C5 | 124.3 | 7.99, s | Thz2-(C2, C4, C=O), Dht-(C2) |
| | C=O | 161.6 | | |
| Glu | NH | | 8.29, brs | Glu-(C2, C3), Thz2-(C=O) |
| | C2 | 52.2 | 5.41, dd, *J* = 1.7, 9.8 | Glu-(C3, C4), Thz3-(C2) |
| | C3 | 79.2 | 4.12, dt, *J* = 2.2, 11.7 | Glu-(C4, C=O), Thz3-(C2), Ind (C3b) |
| | C4 | 37.9 | H4a: 2.44, m<br>H4b: 2.69, m | Glu-(C2, C3, C=O)<br>Glu-(C2, C3, C=O) |
| | C=O | 172.0 | | |
| Thz3 | C2 | 154.3 | | |
| | C4 | 130.9 | | |
| | C5 | 160.1 | | |
| | C=O | 160.3 | | |
| Ser | NH | | 7.92, d, *J* = 10.9 | Ser-(C2, C3), Thz3-(C4, C=O) |
| | C2 | 48.6 | 5.67, dd, *J* = 6.0, 10.9 | Ser-(C3), Thz3-(C=O), Thz4-(C2) |
| | C3 | 64.4 | H3a: 4.40, d, *J* = 11.3<br>H3b: 5.26, dd, *J* = 6.0, 11.3 | Ser-(C2), Thz4-(C2), Ind-(C=O)<br>Ser-(C2), Thz4-(C2), Ind-(C=O) |
| Thz4 | C2 | 169.9 | | |
| | C4 | 154.7 | | |
| | C5 | 121.3 | 7.79, s | Thz4-(C2, C4), Pyr-(C6) |

| | | | | |
|---|---|---|---|---|
| Ind | C=O | 161.5 | | |
| | C2 | 127.1 | | |
| | C3 | 110.1 | | |
| | C3a | 119.5 | | |
| | C3b | 65.9 | H3ba: 4.28, *ol*<br>H3bb: 4.93, d, *J* = 10.5 | Ind-(C2, C3, C3a), Glu-(C3)<br>Ind-(C2, C3, C3a) |
| | C4 | 128.4 | | |
| | C4a | 67.8 | H4aa: 4.98, d, *J* = 12.6<br>H4ab: 5.85, d, *J* = 12.4 | Ind-(C3a, C4, C5), Glu-(C=O)<br>Ind-(C3a, C4, C5), Glu-(C=O) |
| | C5 | 123.5 | 7.15, d, *J* = 7.1 | Ind-(C3a, C4a, C6, C7) |
| | C6 | 125.1 | 7.39, dd, *J* = 6.9, 8.4 | Ind-(C4, C5, C7, C7a) |
| | C7 | 112.0 | 7.78, *ol* | Ind-(C3a, C5) |
| | C7a | 135.4 | | |
| | N-OH | | 10.52, s | Ind-(C2) |
| Sug1 | C1 | 102.0 | 5.45, d, *J* = 1.8 | Thz3-(C5), Sug1-(C3, C5) |
| | C2 | 76.0 | 4.04, dd, *J* = 1.9, 3.3 | Sug1-(C1, C3, C4, C2-OMe) |
| | C2-OMe | 59.5 | 3.53, s | Sug1-(C2) |
| | C3 | 80.2 | 3.64, m | Sug1-(C1, C5, C3-OMe) |
| | C3-OMe | 58.0 | 3.46, s | Sug1-(C3) |
| | C4 | 77.2 | 3.68, *ol* | Sug1-(C2, C5, C6), Sug2-(C1) |
| | C5 | 70.2 | 3.63, *ol* | Sug1-(C1, C3, C4, C6) |
| | C6 | 17.8 | 1.35, d, *J* = 5.5 | Sug1-(C4, C5) |
| Sug2 | C1 | 102.2 | 4.66, dd, *J* = 2.2, 9.1 | Sug1-(C4), Sug2-(C2, C5) |
| | C2 | 30.9 | H2a: 1.46, m<br>H2b: 1.84, m | Sug2-(C1, C3, C4) |
| | C3 | 30.0 | H3a: 1.50, m<br>H3b: 2.12, m | Sug2-(C1, C4) |
| | C4 | 80.6 | 3.11, m | Sug2-(C5, C6), Sug3-(C1) |
| | C5 | 74.2 | 3.29, m | Sug2-(C1, C3, C6) |
| | C6 | 18.1 | 1.16, d, *J* = 6.2 | Sug2-(C4, C5) |
| Sug3 | C1 | 101.1 | 4.47, dd, *J* = 2.0, 9.7 | Sug2-(C4), Sug3-(C2, C3, C5) |
| | C2 | 39.1 | H2a: 1.50, m<br>H2b: 2.10, m | Sug3-(C1, C3, C4) |
| | C3 | 71.5 | 3.48, *ol* | Sug3-(C1, C5) |
| | C4 | 77.2 | 2.96, m | Sug3-(C2, C5, C6) |
| | C5 | 71.8 | 3.18, m | Sug3-(C1, C4, C6) |
| | C6 | 17.8 | 1.25, d, *J* = 6.1 | Sug3-(C4, C5) |
| Sug4 | C1 | 100.6 | 5.18, d, *J* = 7.7 | Pyr-(C3), Sug4-(C3, C5) |
| | C2 | 69.6 | 4.2, t, *J* = 8.8 | Sug4-(C1, C3) |
| | C3 | 84.3 | 3.24, d, *J* = 9.5 | Sug4-(C2, C4, C3-OMe) |
| | C3-OMe | 58.2 | 3.50, s | Sug4-(C3) |
| | C4 | 77.7 | 3.46, *ol* | Sug4-(C2, C4-OMe) |
| | C4-OMe | 61.9 | 3.57, s | Sug4-(C4) |
| | C5 | 71.3 | 3.79, m | Sug4-(C1, C3, C4, C6) |
| | C6 | 16.8 | 1.36, d, *J* = 6.5 | Sug4-(C4, C5) |

* overlapped

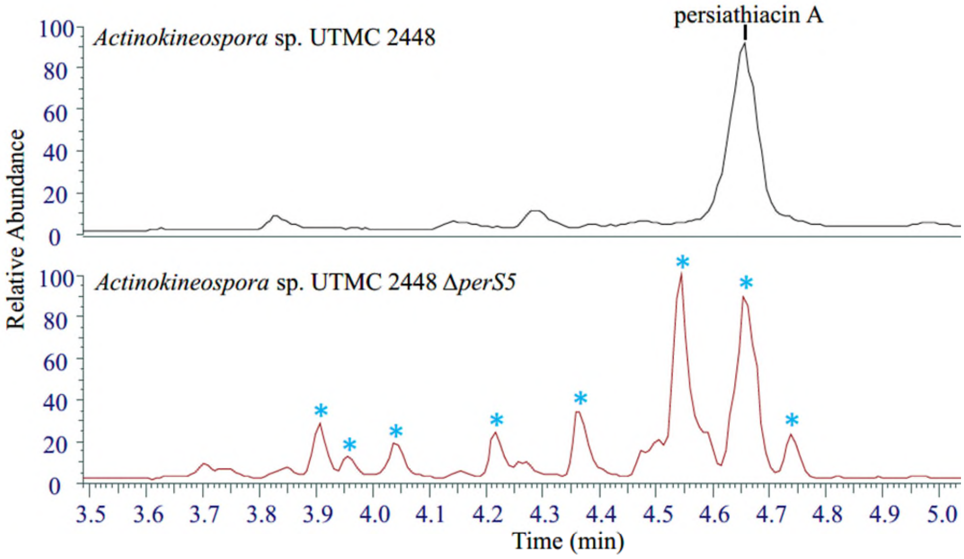


**Figure S5.** Comparative LC-MS profiles of extracts from wild-type *Actinokineospora* sp. UTMC 2448 and the *perS5* deleted mutant. Blue stars indicate novel glycosylated analogues.

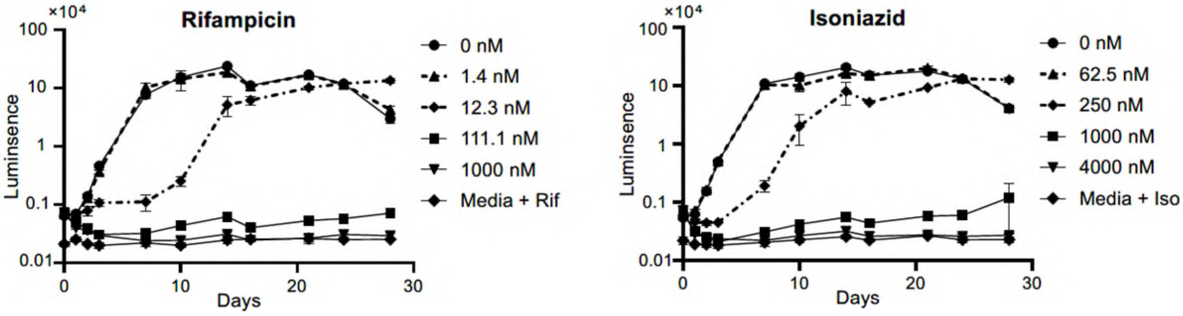


**Figure S6.** Time–kill kinetics of rifampicin and isoniazid against *M. tuberculosis* H37Rv lux.

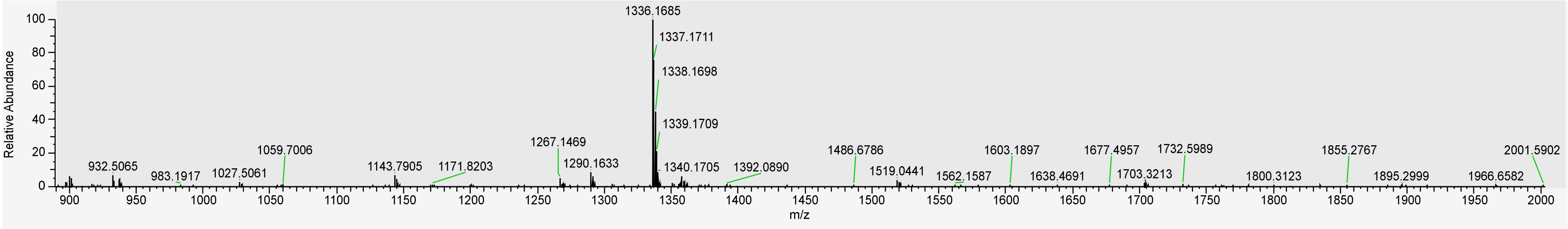

**Figure S7.** High resolution mass spectrometry profiles of compound **7**.

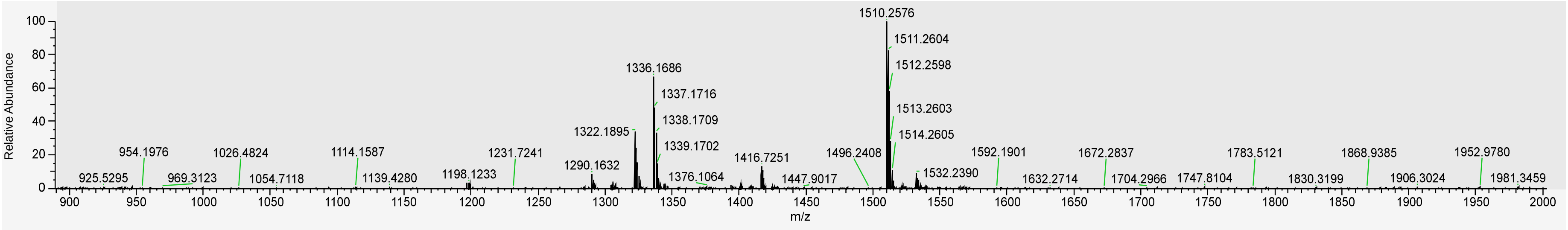

**Figure S8.** High resolution mass spectrometry profiles of compound **8**.

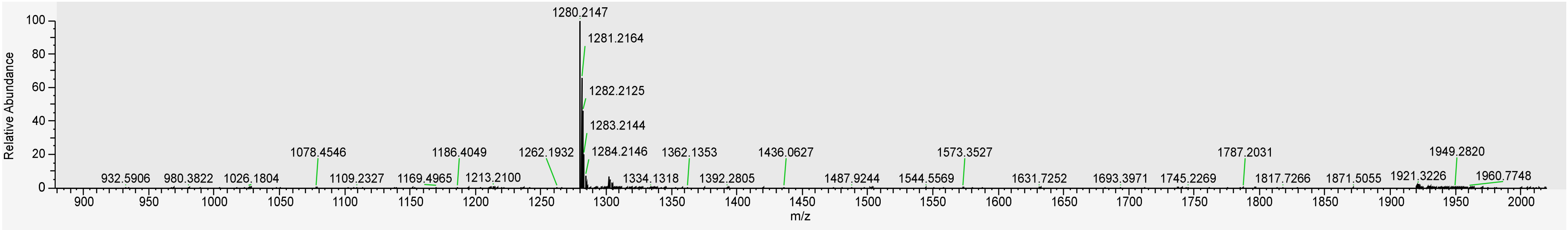

**Figure S9.** High resolution mass spectrometry profiles of compound **9**.

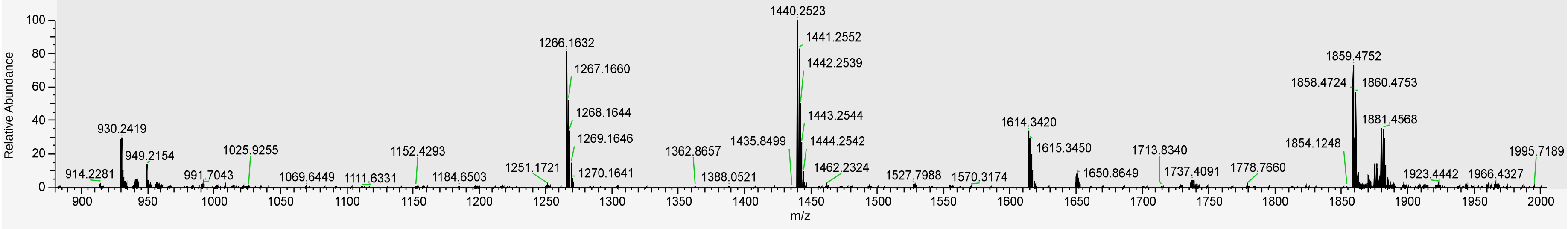


**Figure S10.** High resolution mass spectrometry profiles of compound **10**.

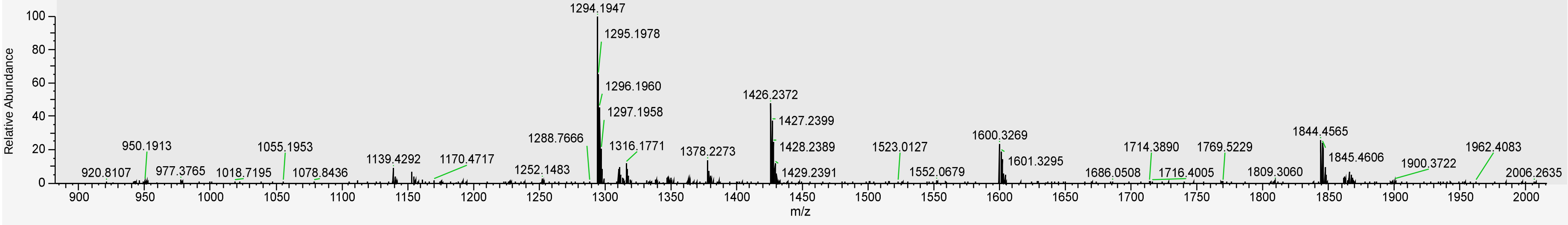


**Figure S11.** High resolution mass spectrometry profiles of compound **11**.

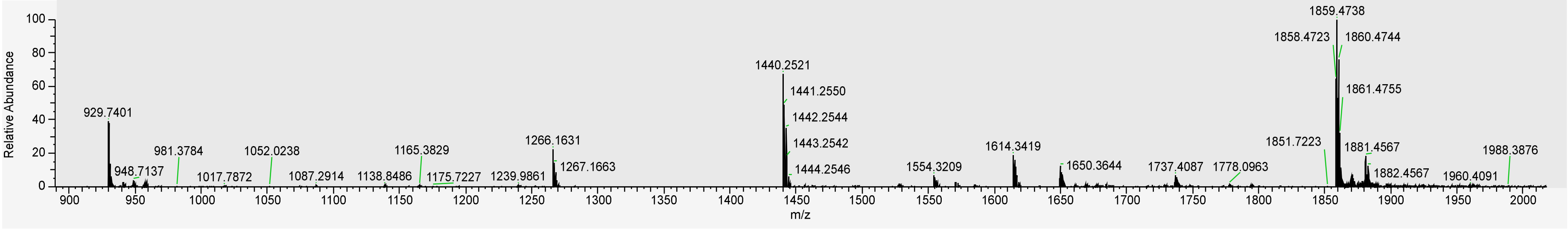


**Figure S12.** High resolution mass spectrometry profiles of compound **12**.

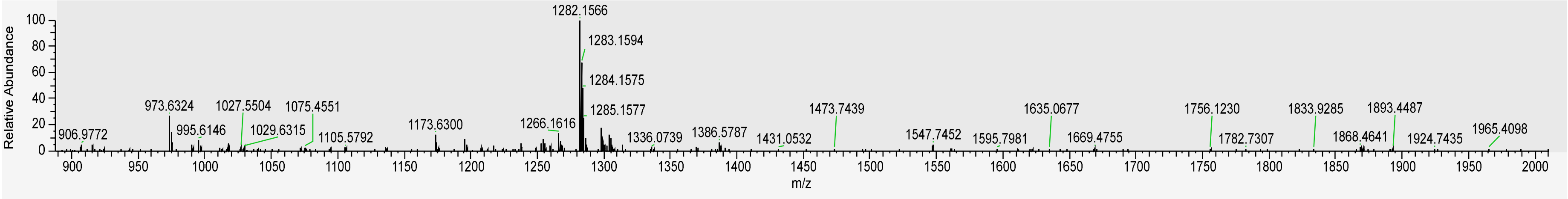


**Figure S13.** High resolution mass spectrometry profiles of compound **13**.

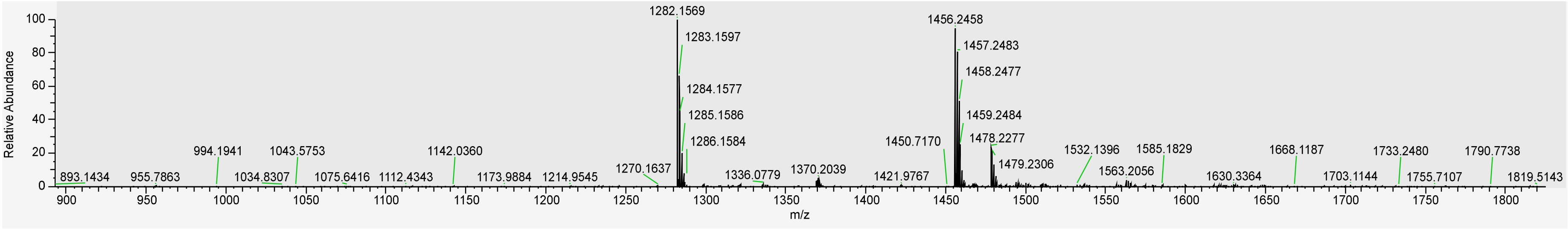


**Figure S14.** High resolution mass spectrometry profiles of compound **14**.

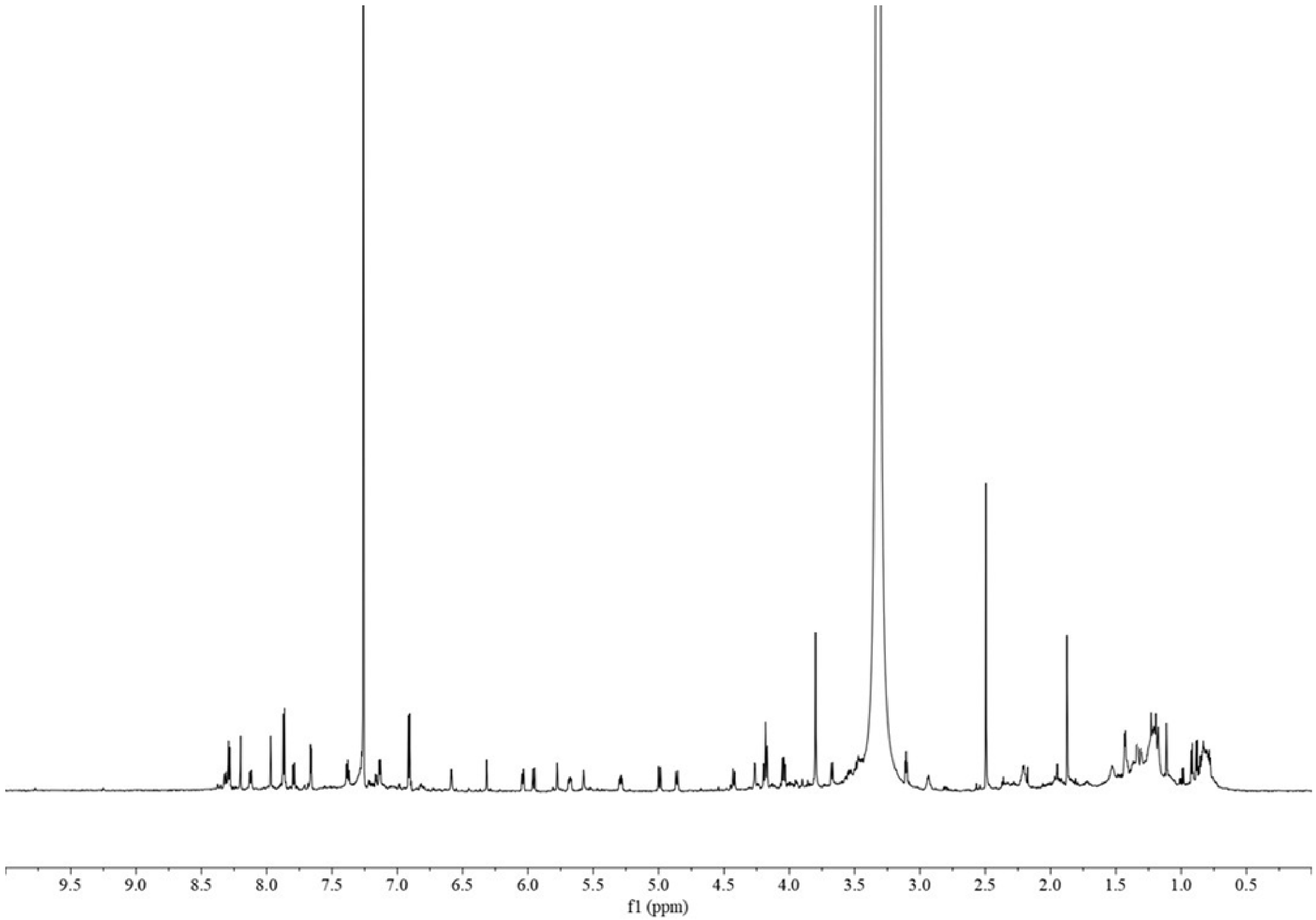


**Figure S15.** Proton spectrum of compound **7** in $CDCl_3$:$CD_3OD$ (9:1).

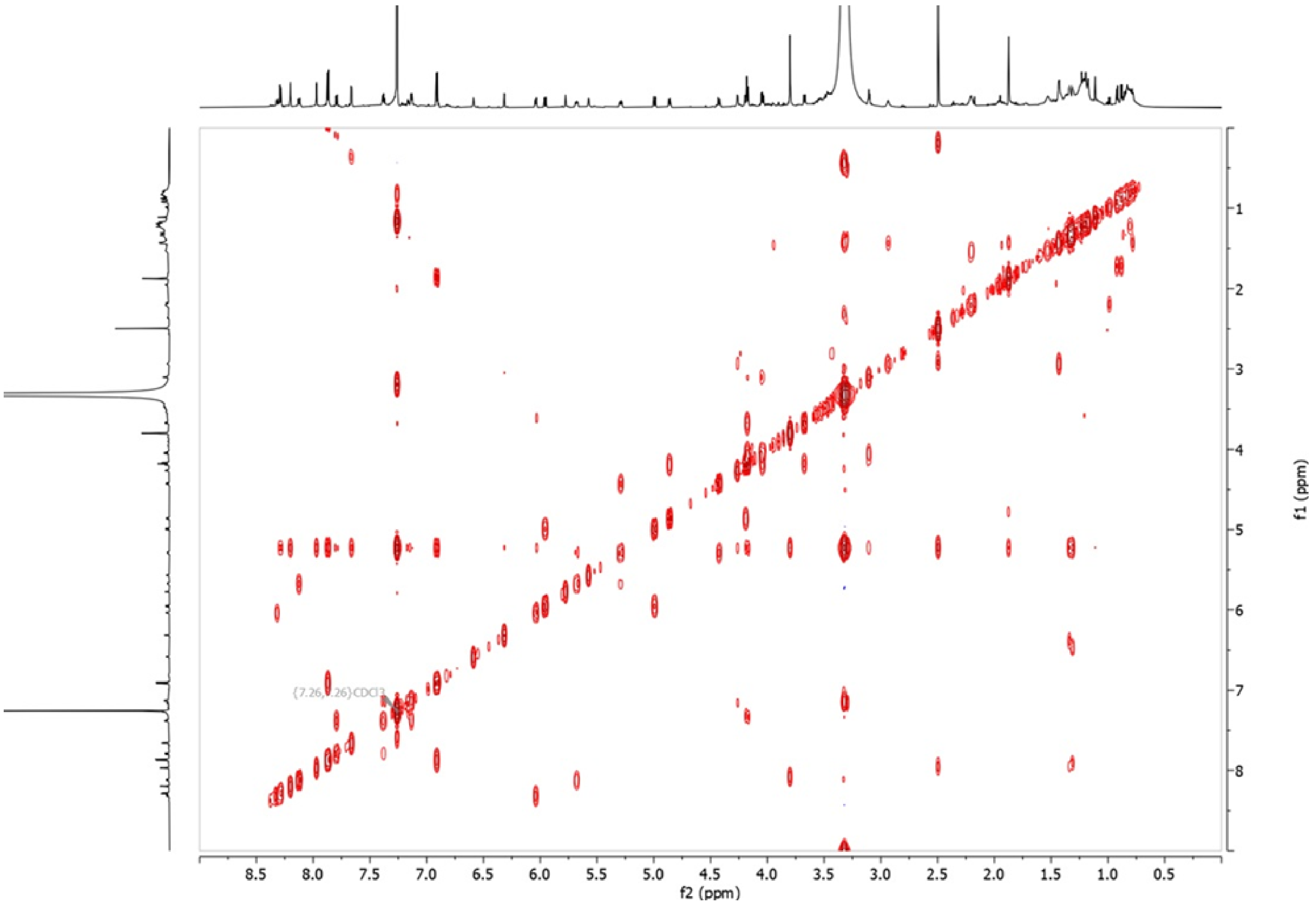


**Figure S16.** COSY spectrum of compound **7** in $CDCl_3$:$CD_3OD$ (9:1).

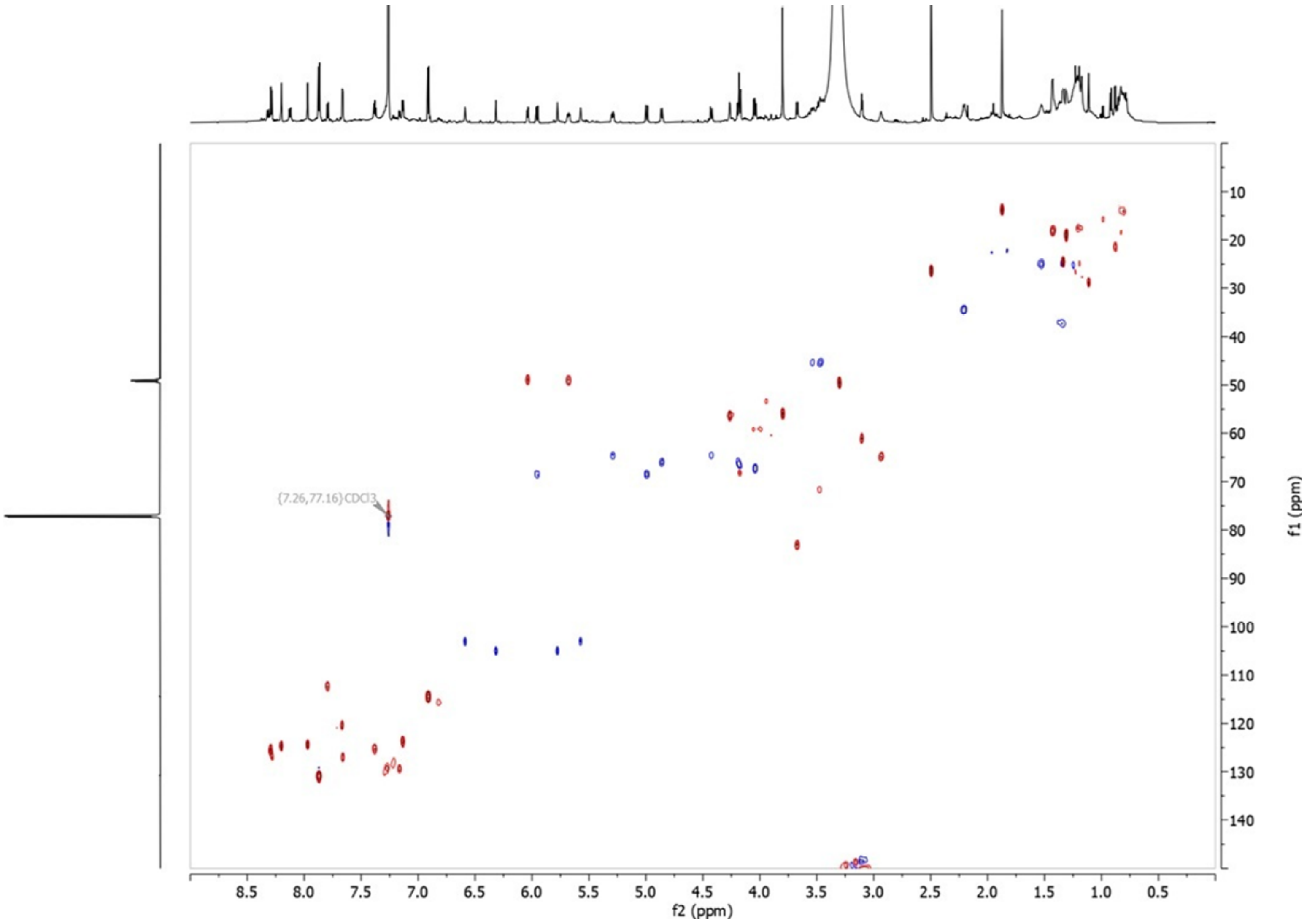


**Figure S17.** HSQC spectrum of compound **7** in $CDCl_3$:$CD_3OD$ (9:1).

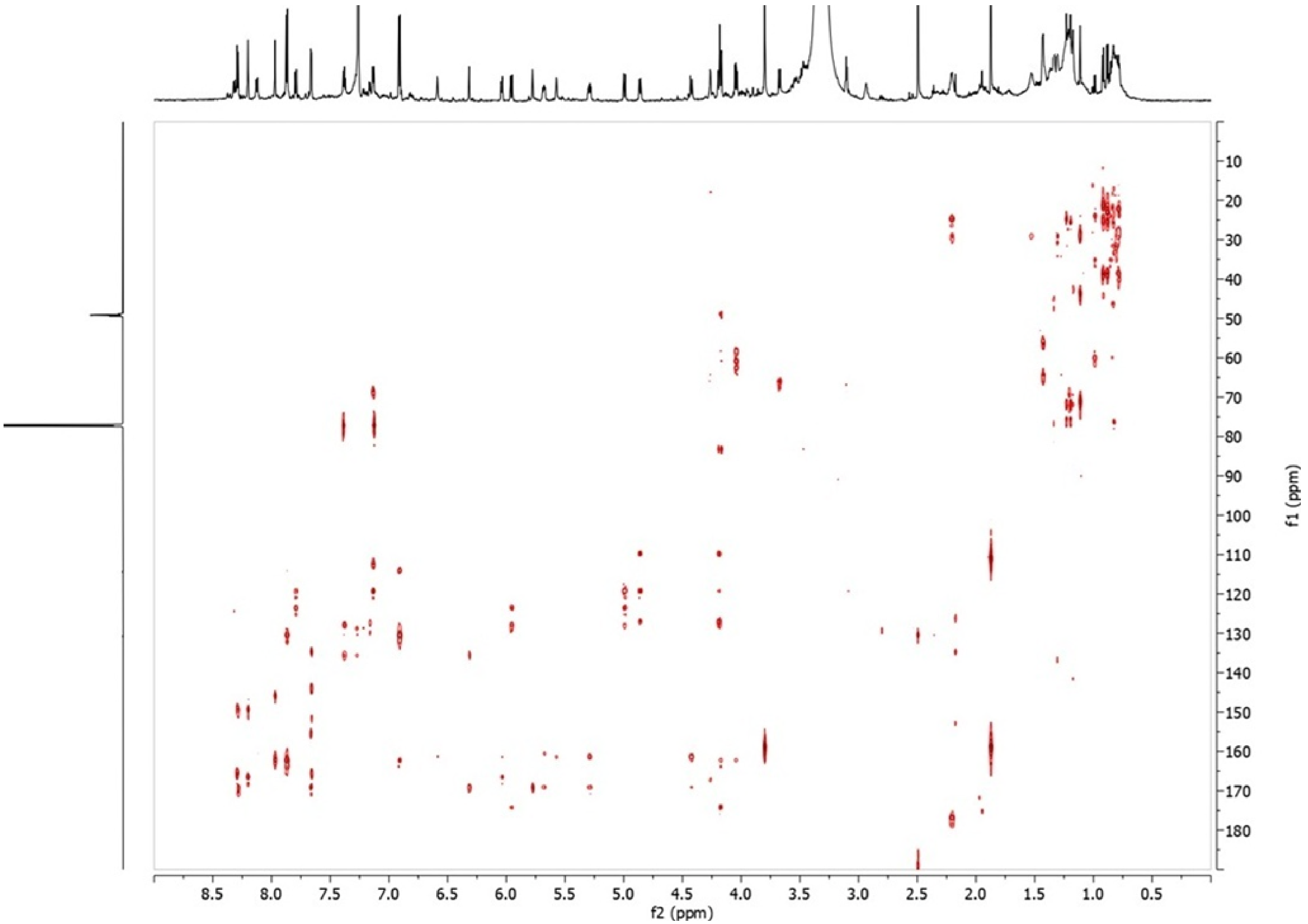


**Figure S18.** HMBC spectrum of compound **7** in $CDCl_3$:$CD_3OD$ (9:1).

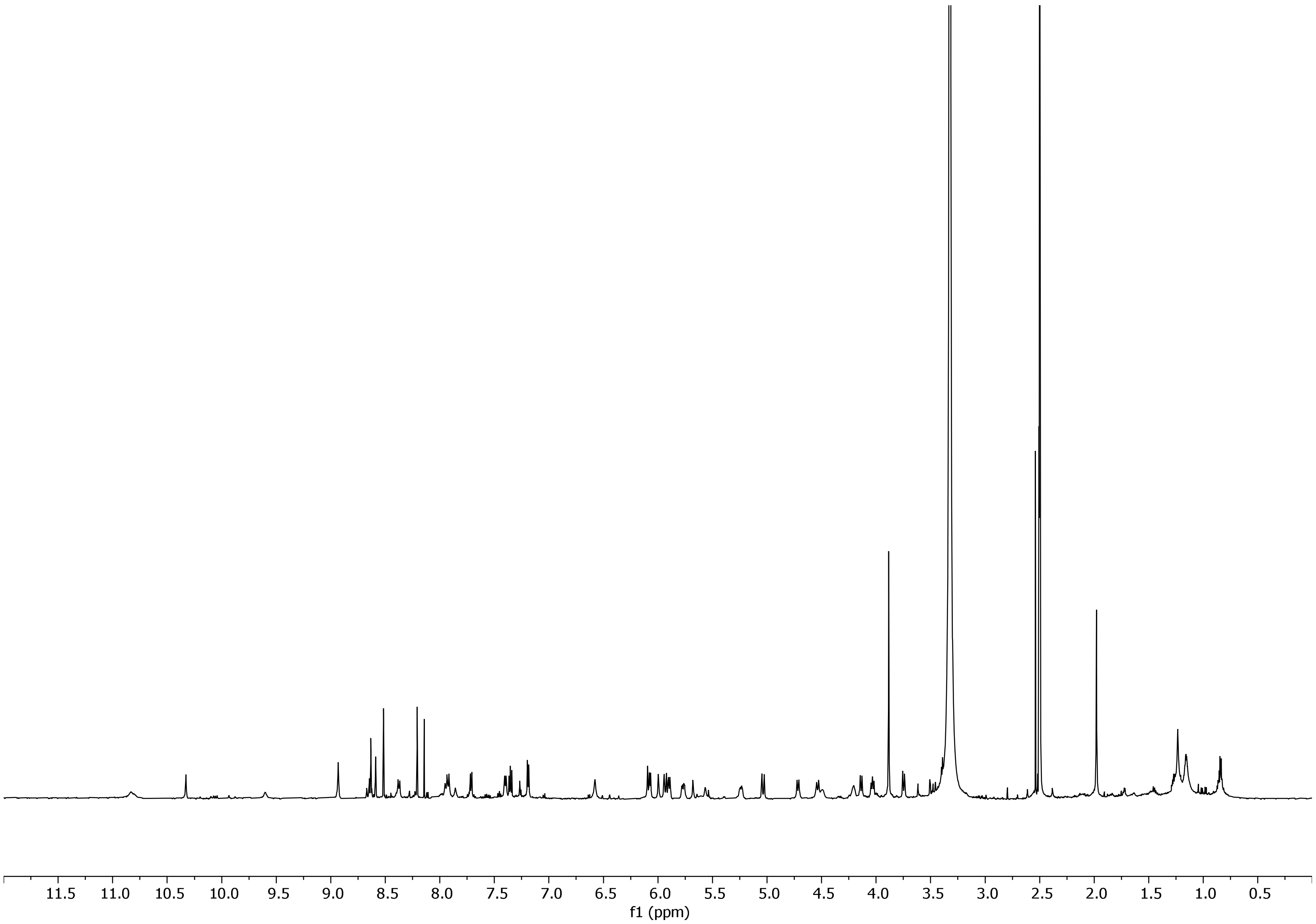


**Figure S19.** Proton spectrum of compound **7** in DMSO-$d_6$.

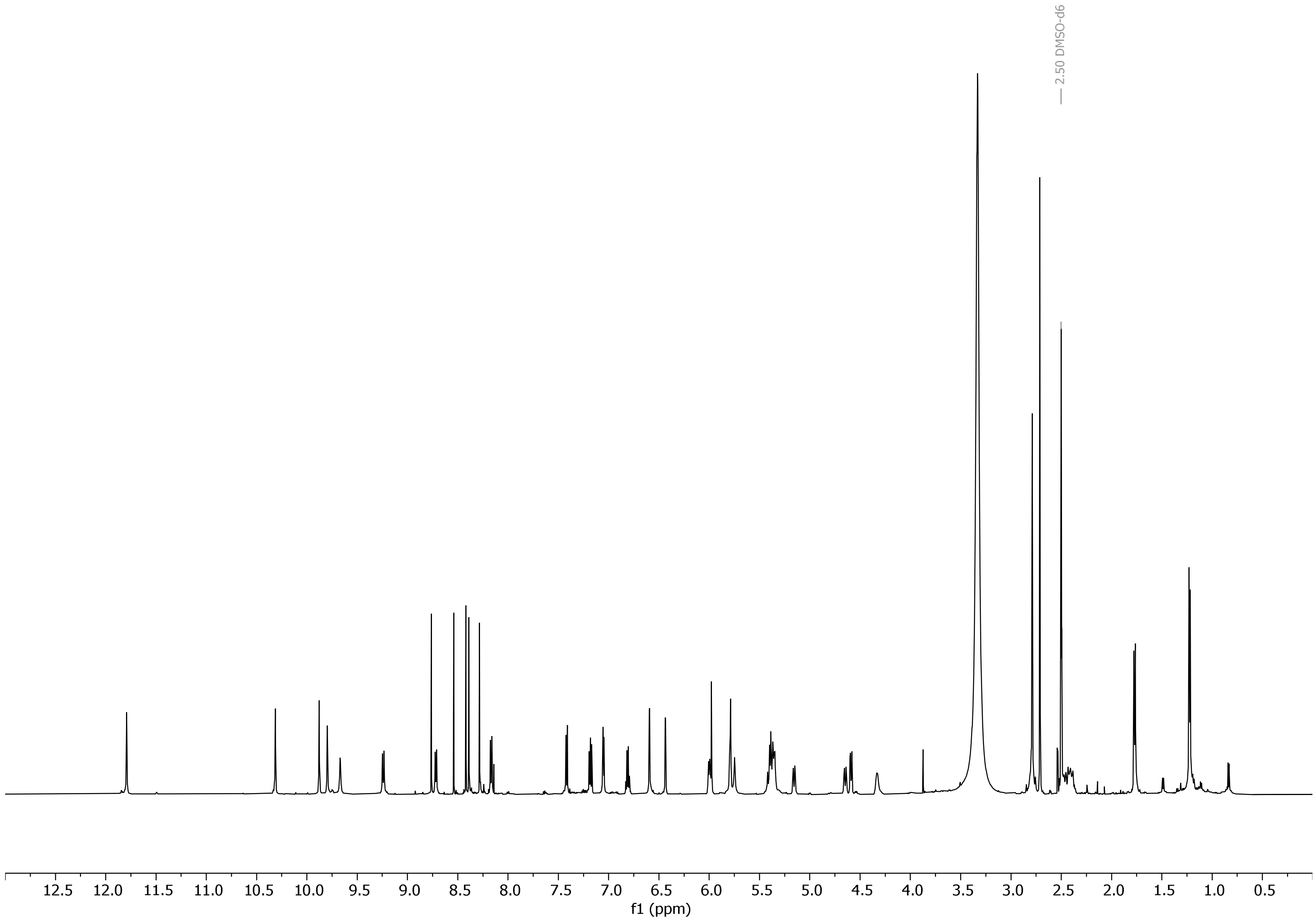


**Figure S20.** Proton spectrum of compound **9** in DMSO-$d_6$.

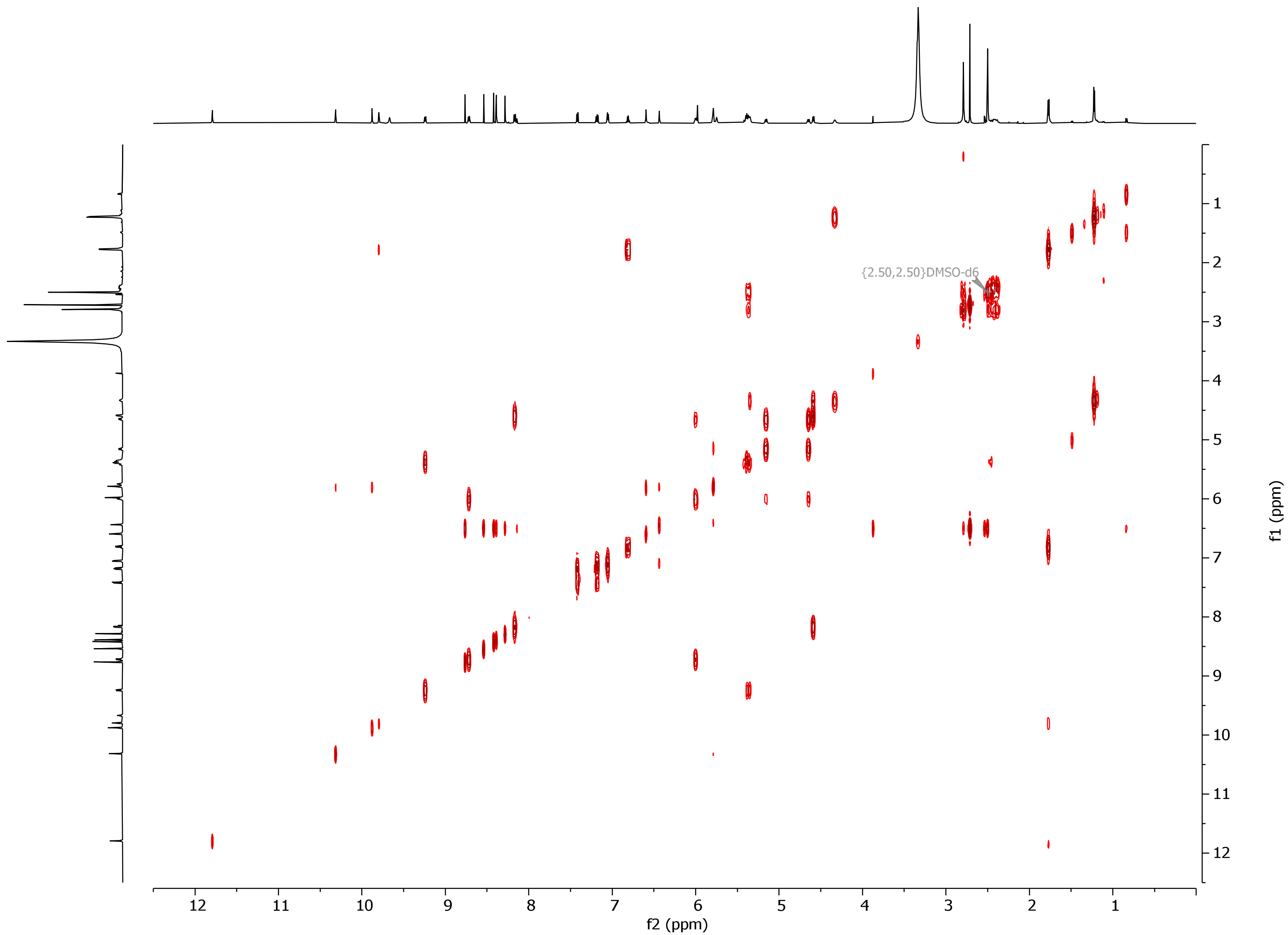


**Figure S21.** COSY spectrum of compound **9** in DMSO-$d_6$.

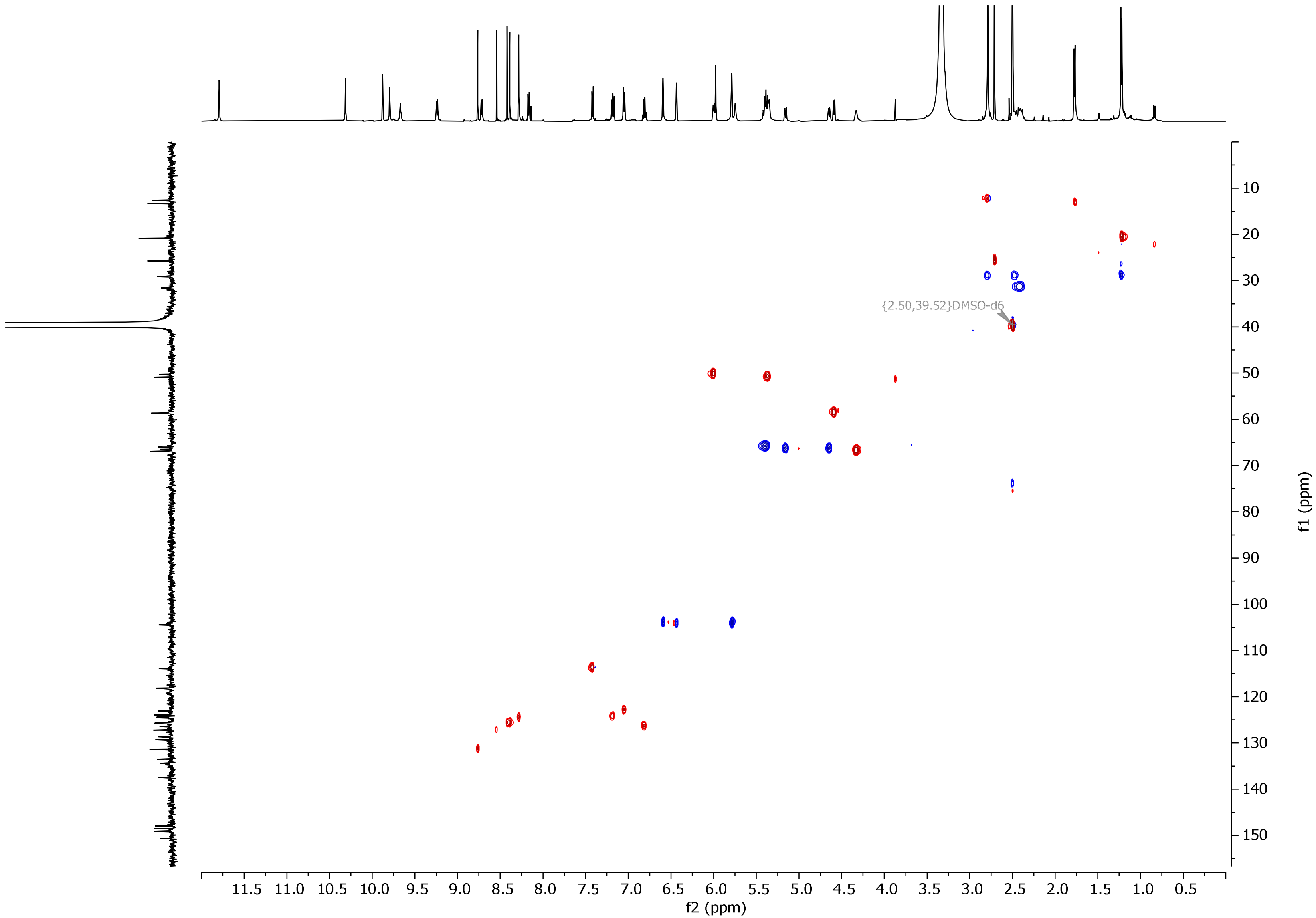


**Figure S22.** HSQC spectrum of compound **9** in DMSO-*d*$_6$.

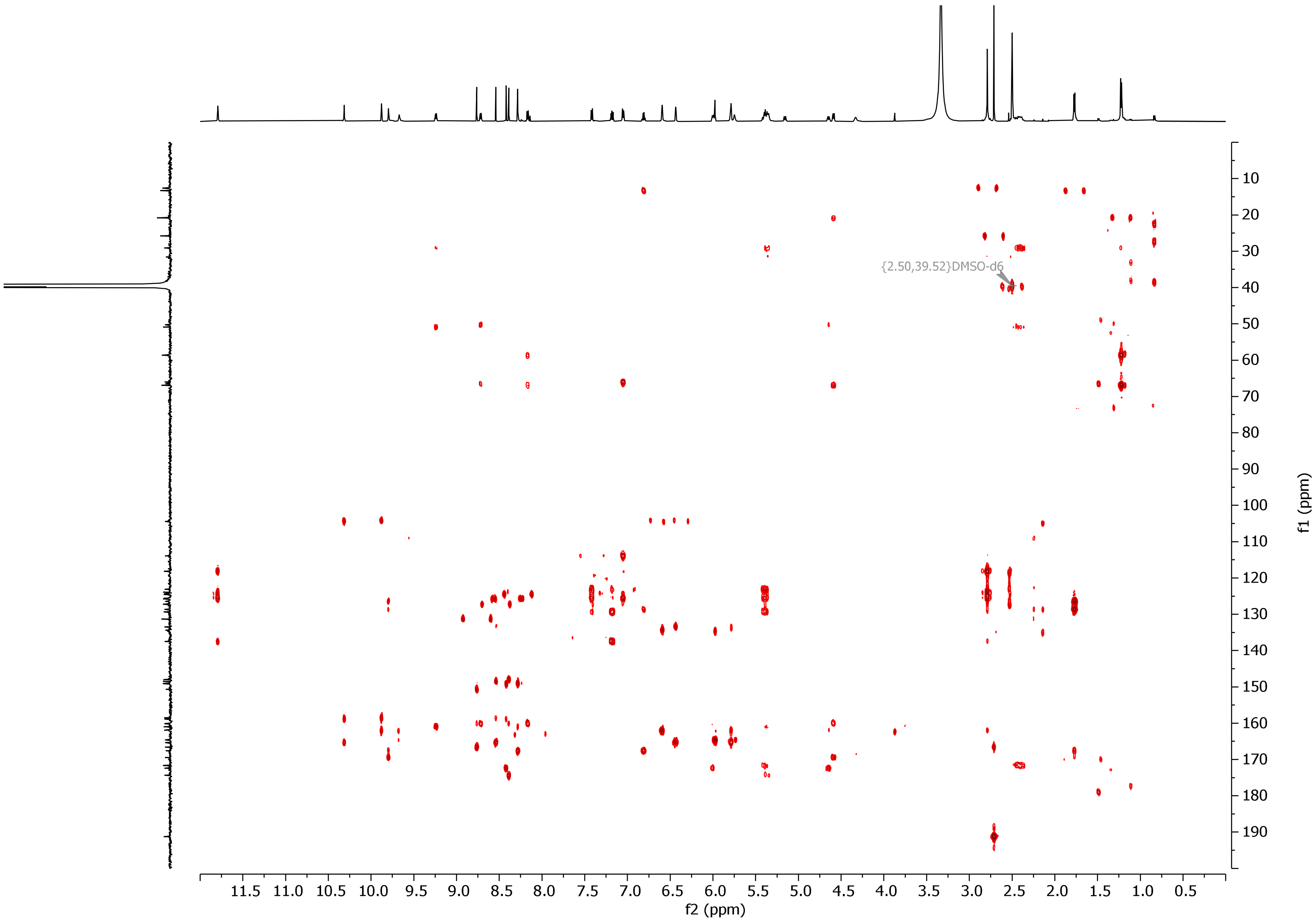


**Figure S23.** HSQC spectrum of compound **9** in DMSO-$d_6$.

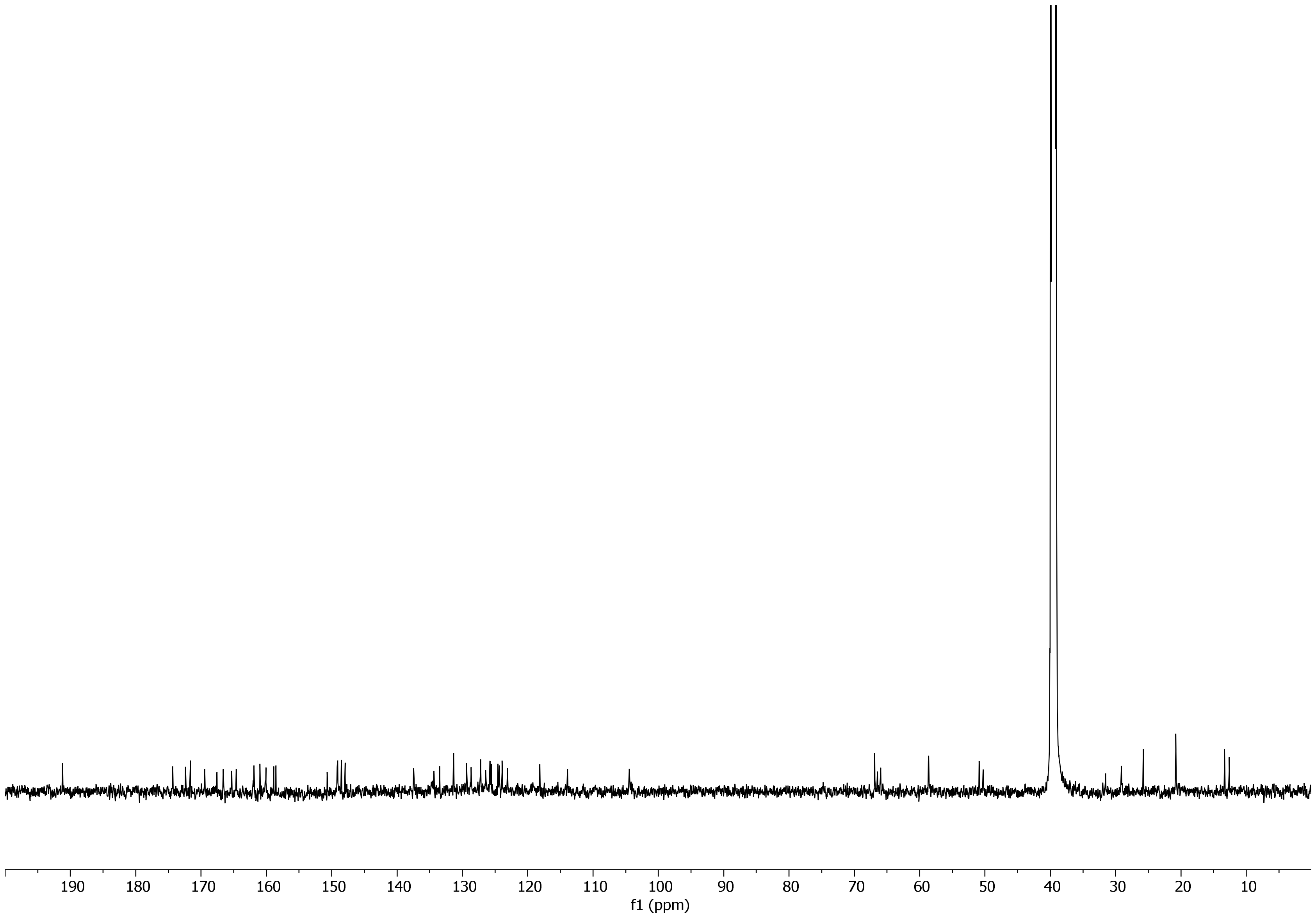


**Figure S24.** $^{13}$C NMR spectrum of compound **9** in DMSO-$d_6$.

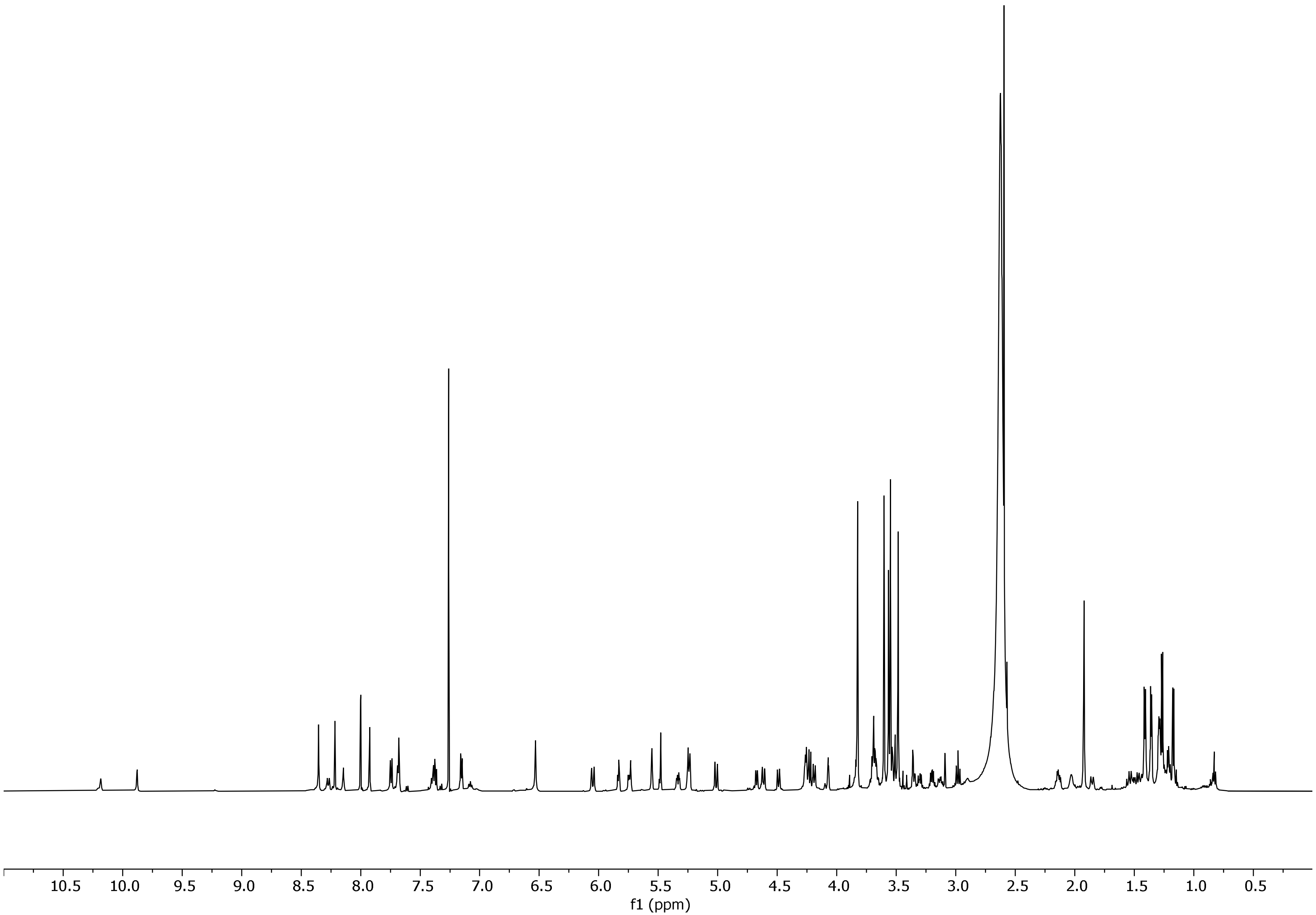


**Figure S25.** Proton spectrum of compound **10** in $CDCl_3$:$CD_3OD$ (9:1).

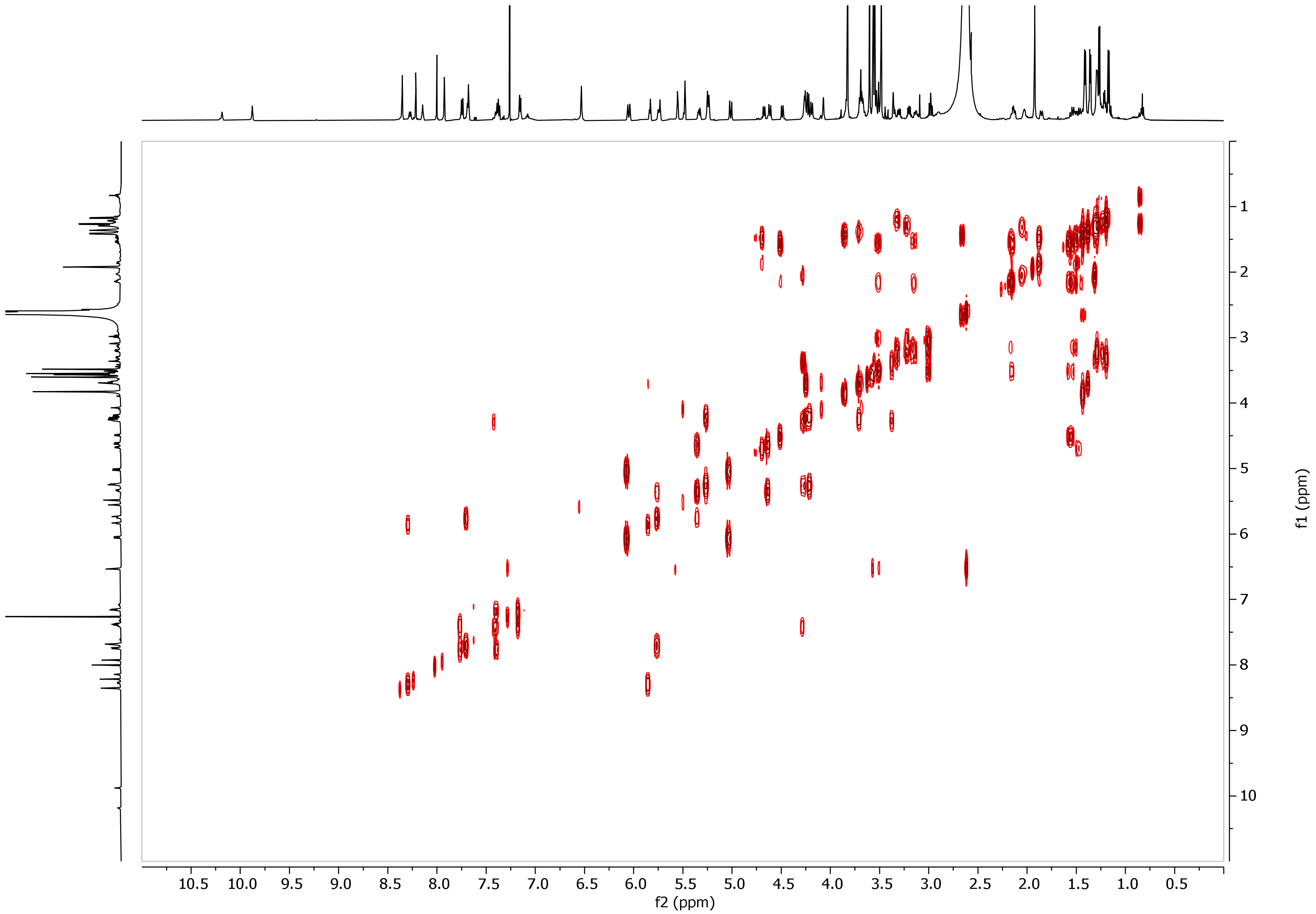


**Figure S26.** COSY spectrum of compound **10** in $CDCl_3$:$CD_3OD$ (9:1).

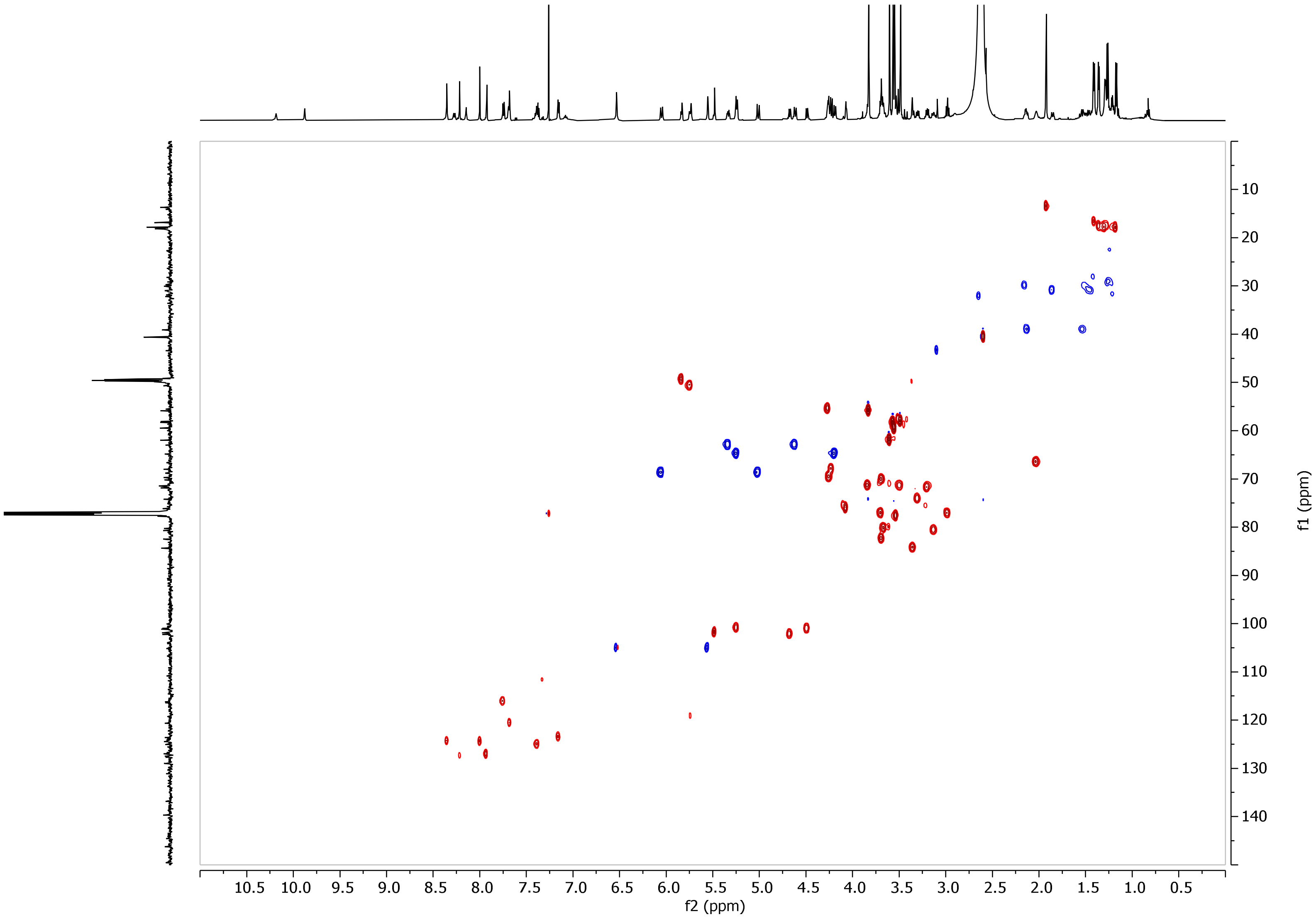


**Figure S27.** HSQC spectrum of compound **10** in $CDCl_3$:$CD_3OD$ (9:1).

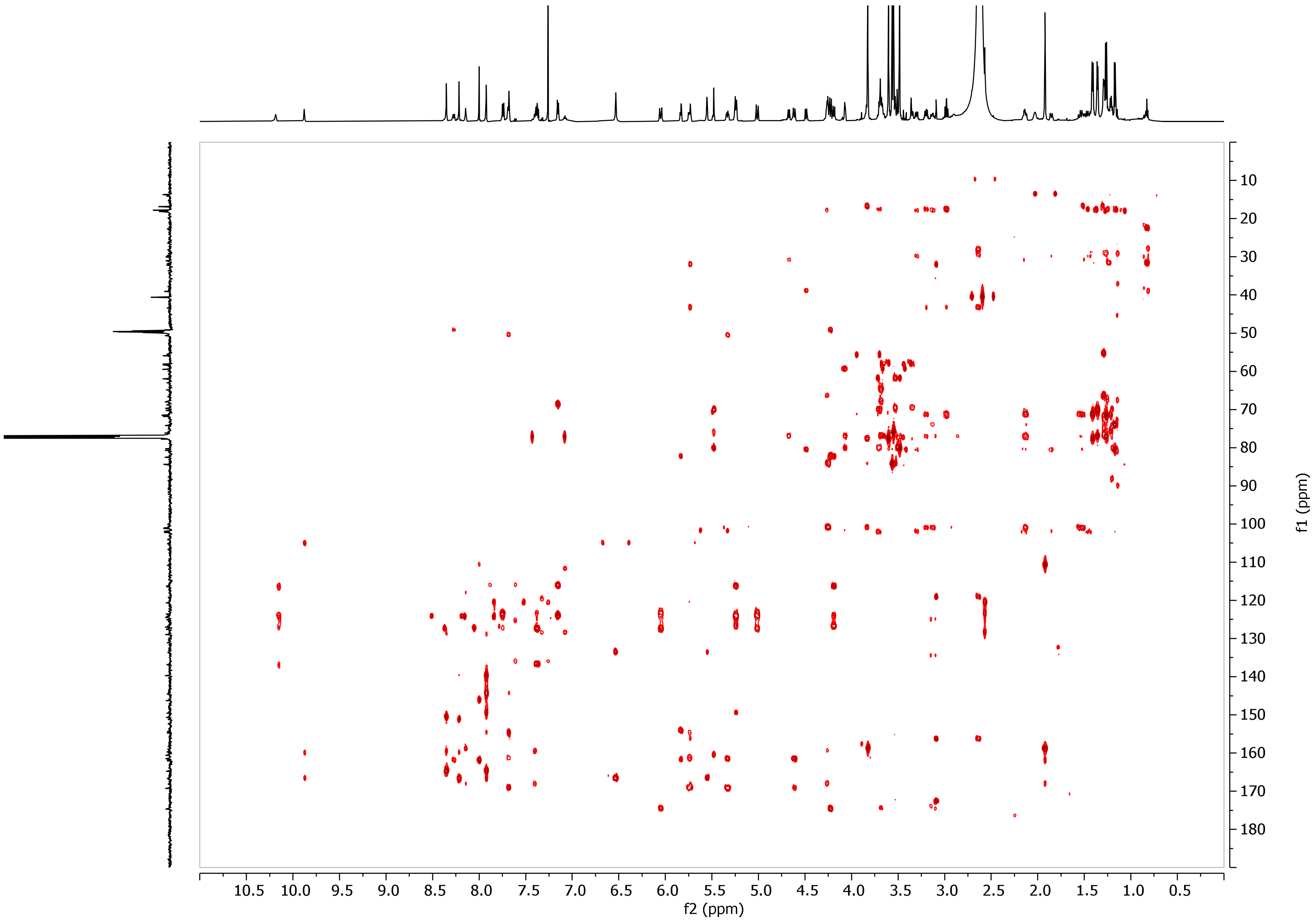


**Figure S28.** HMBC spectrum of compound **10** in $CDCl_3$:$CD_3OD$ (9:1).

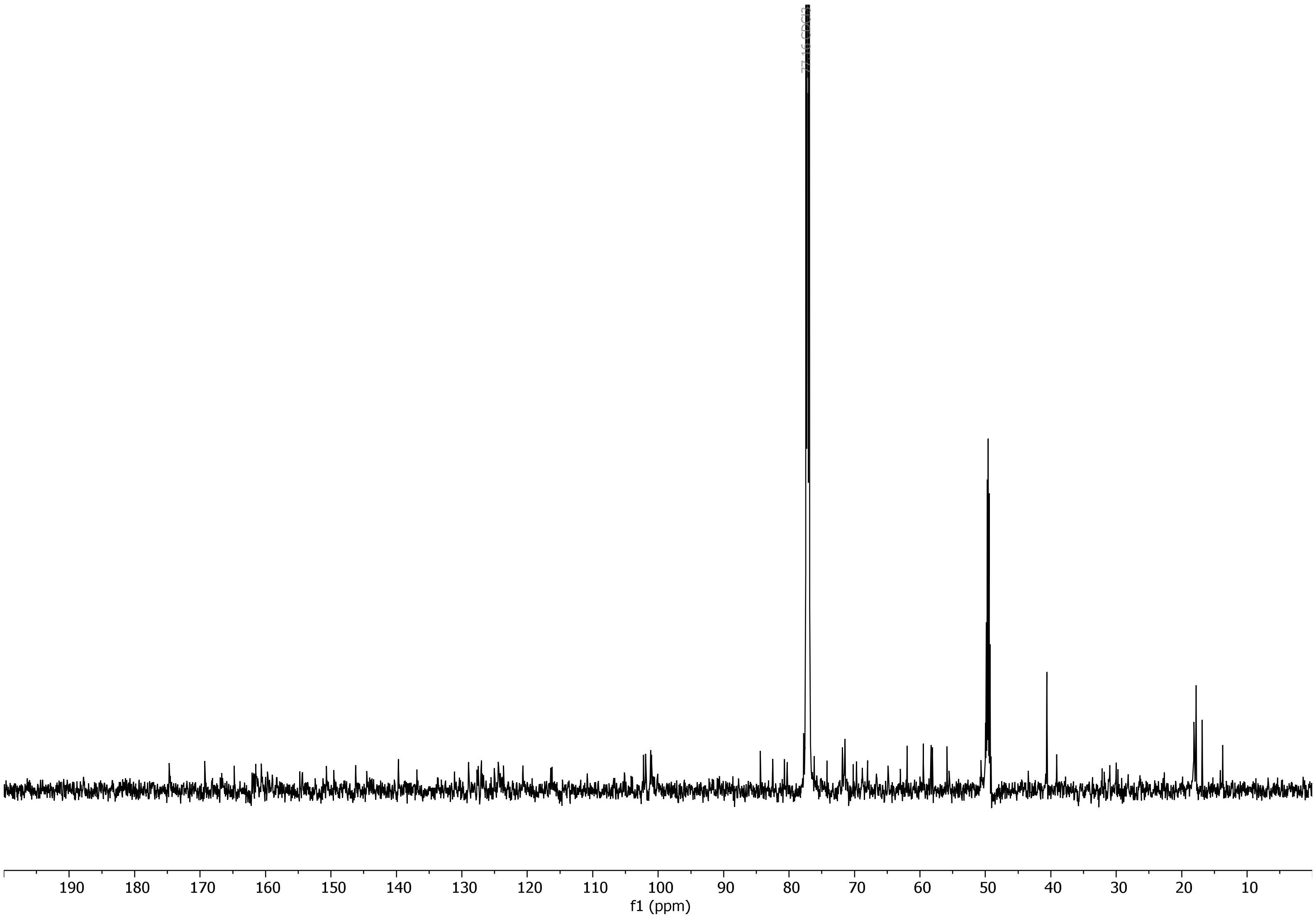

**Figure S29.** Carbon spectrum of compound **10** in $CDCl_3$:$CD_3OD$ (9:1).

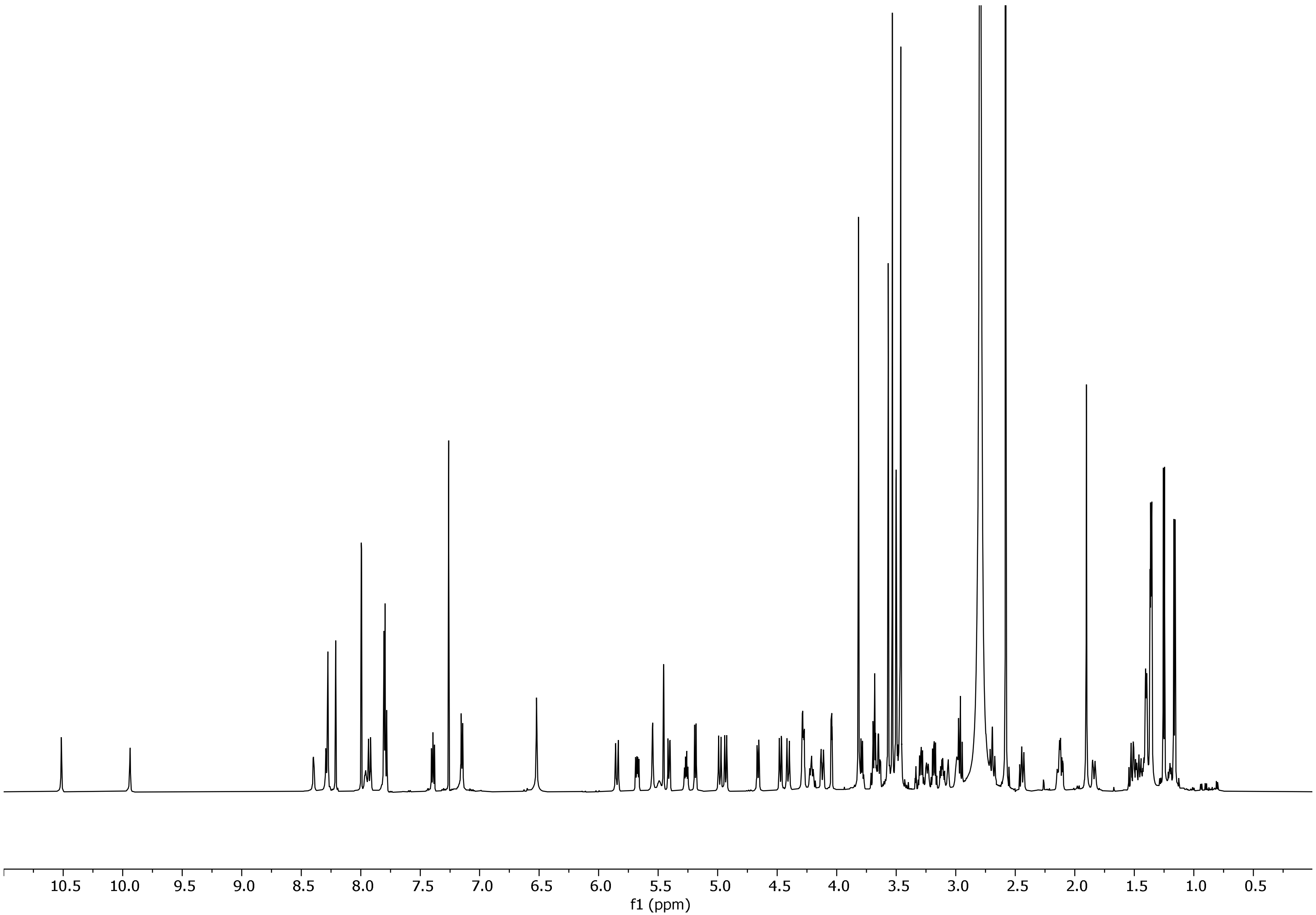


**Figure S30.** Proton spectrum of compound **12** in $CDCl_3$:$CD_3OD$ (9:1).

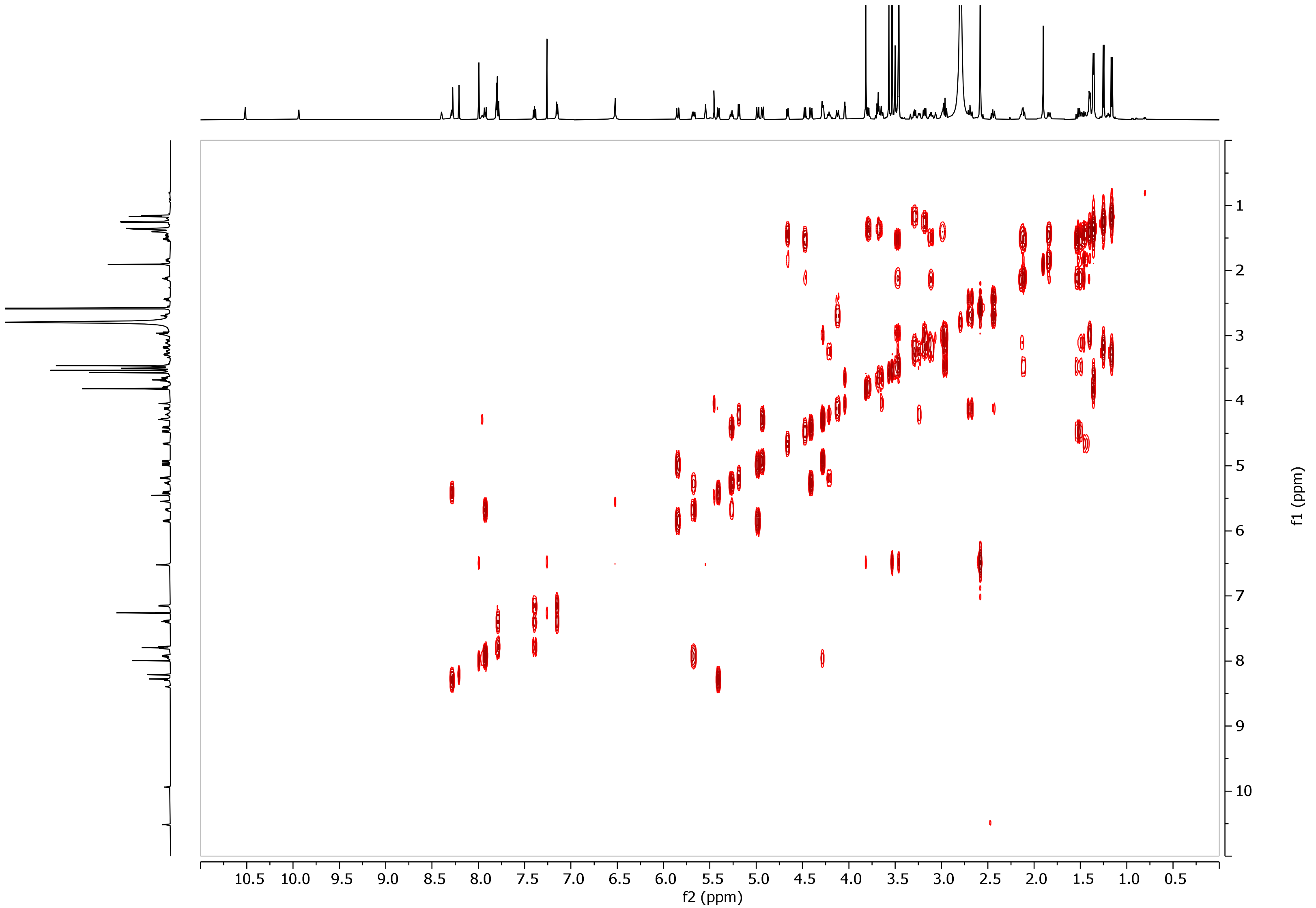


**Figure S31.** COSY spectrum of compound **12** in $CDCl_3$:$CD_3OD$ (9:1).

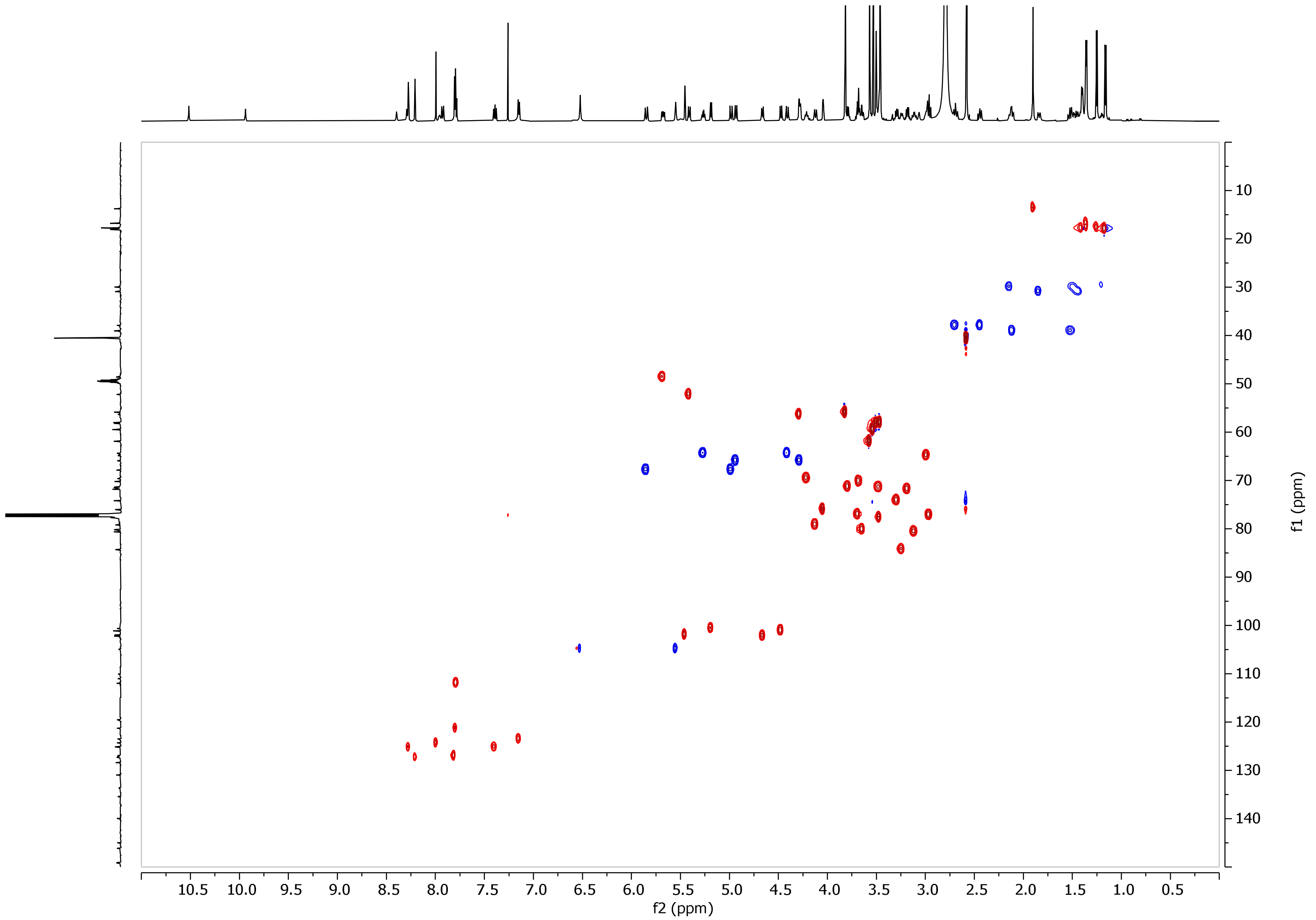


**Figure S32.** HSQC spectrum of compound **12** in $CDCl_3$:$CD_3OD$ (9:1).

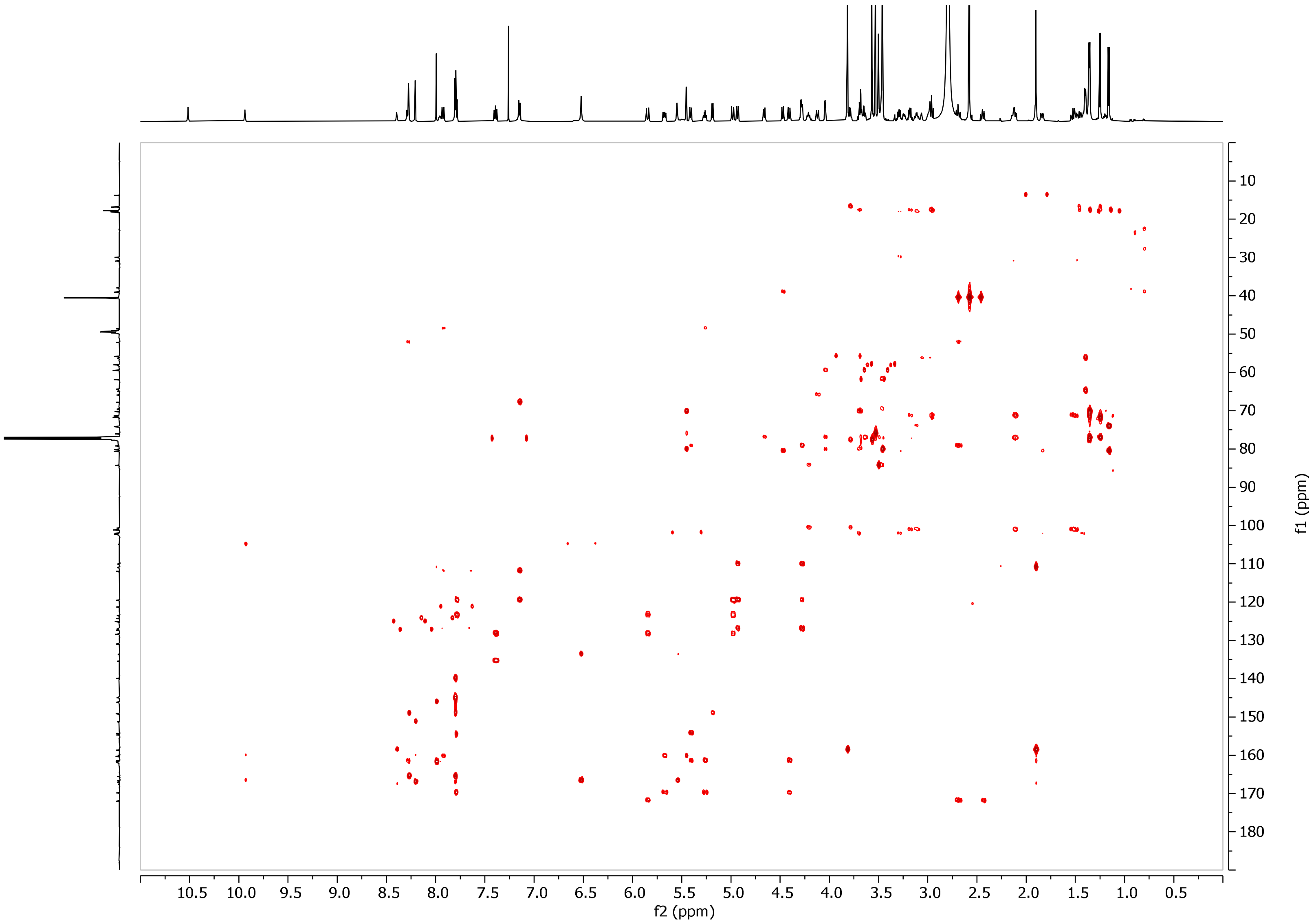


**Figure S33.** HMBC spectrum of compound **12** in $CDCl_3$:$CD_3OD$ (9:1).

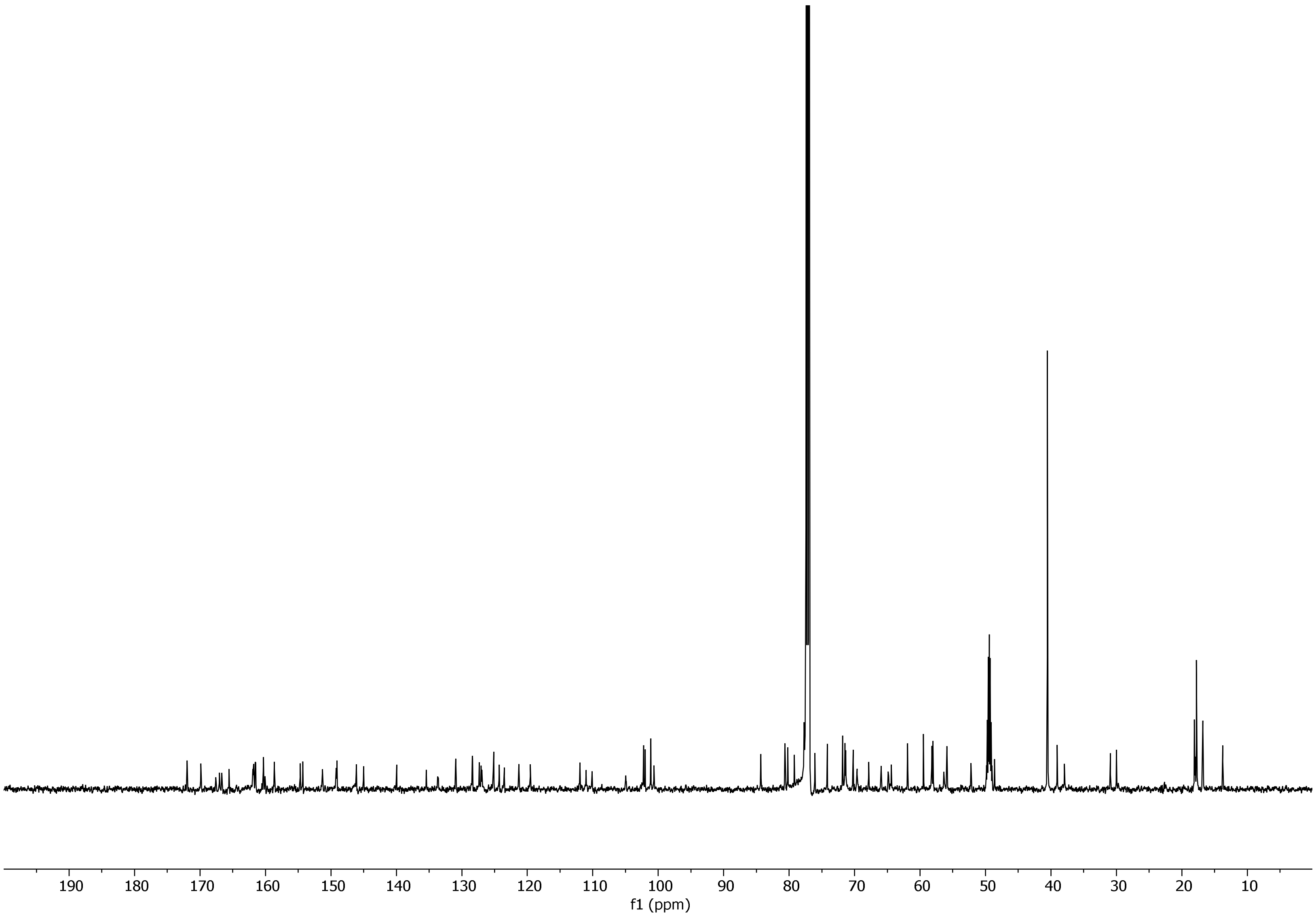

**Figure S34.** Carbon spectrum of compound **12** in $CDCl_3$:$CD_3OD$ (9:1).

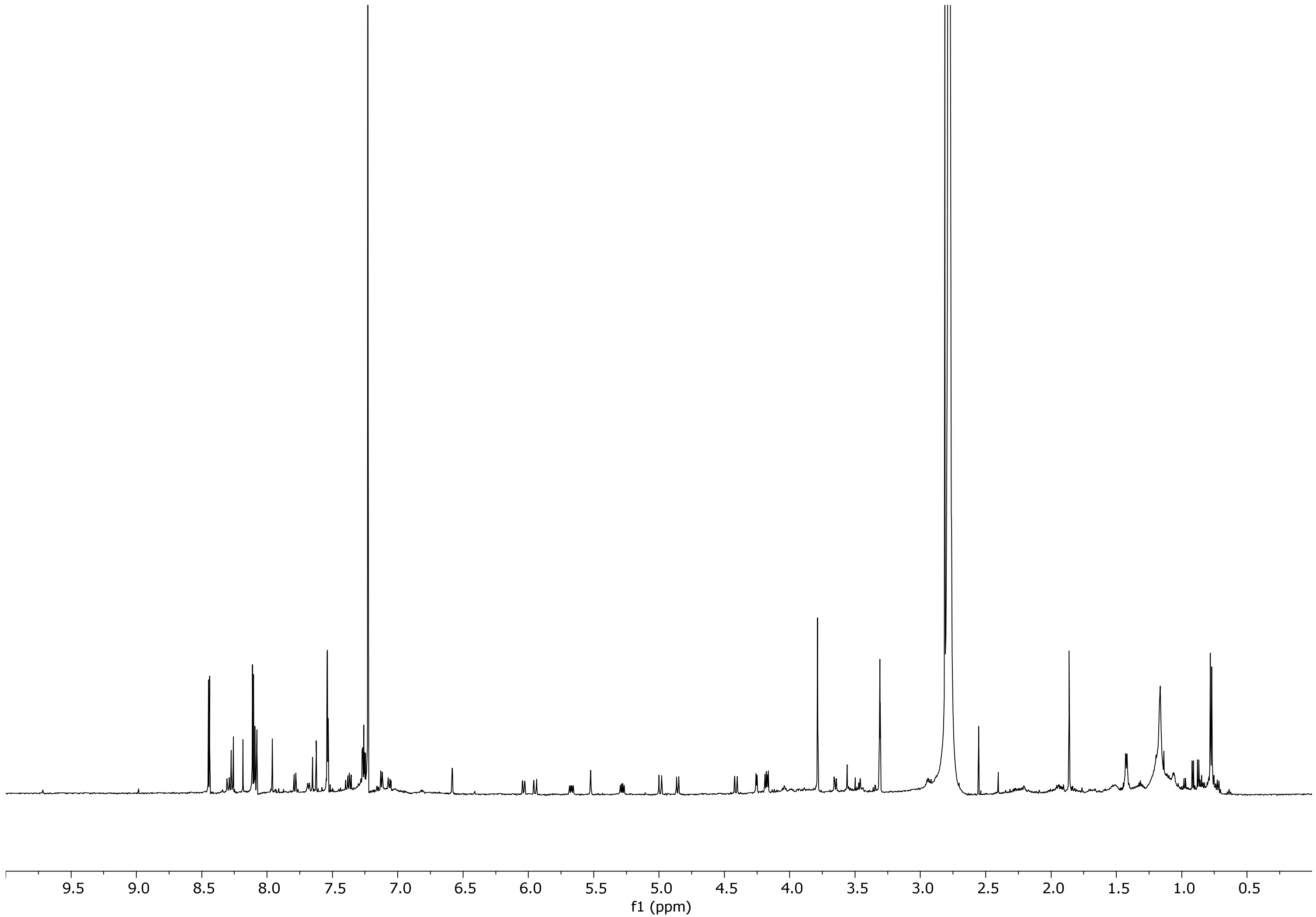


**Figure S35.** Proton spectrum of compound **13** in $CDCl_3$:$CD_3OD$ (9:1).

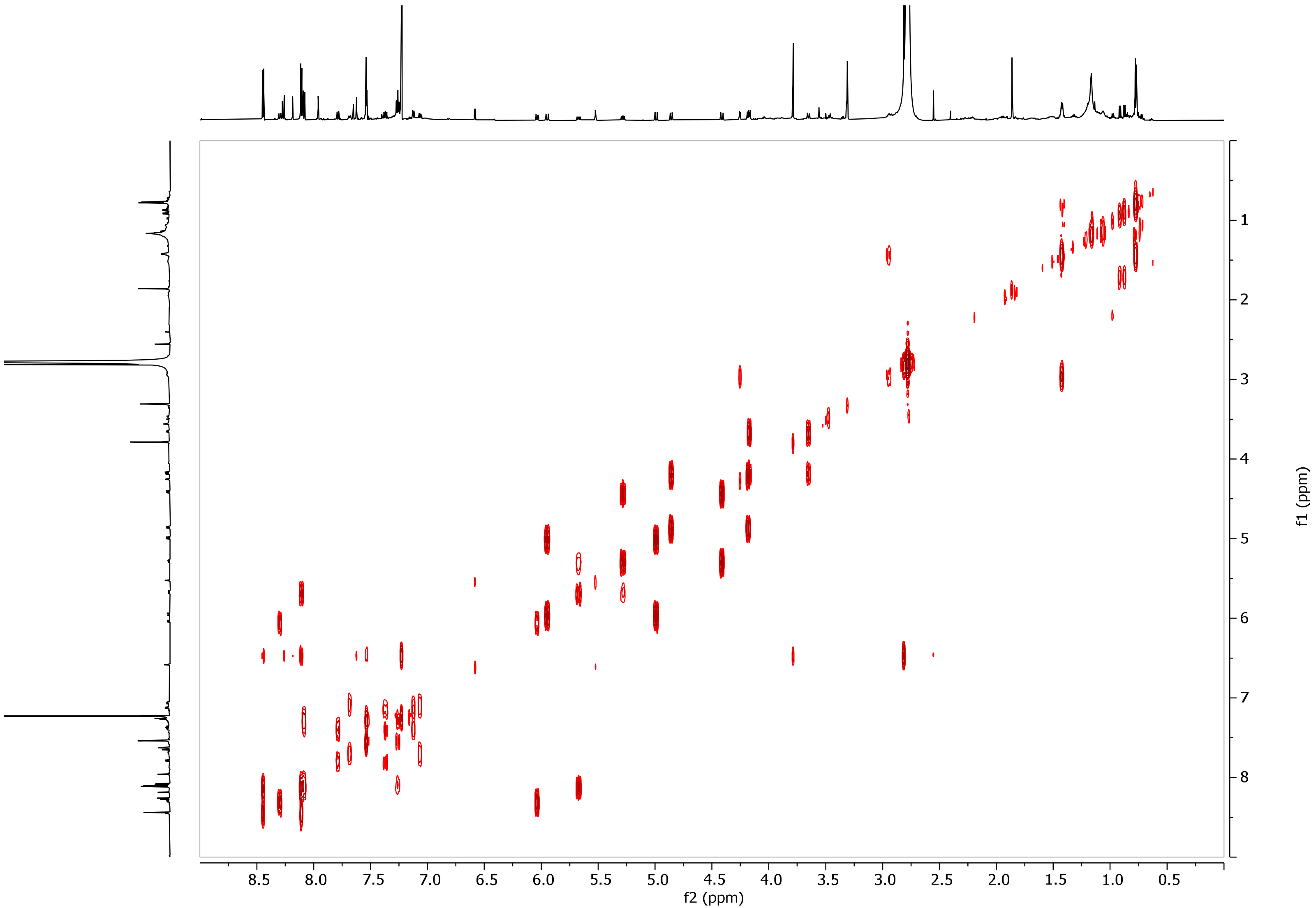


**Figure S36.** COSY spectrum of compound **13** in $CDCl_3$:$CD_3OD$ (9:1).

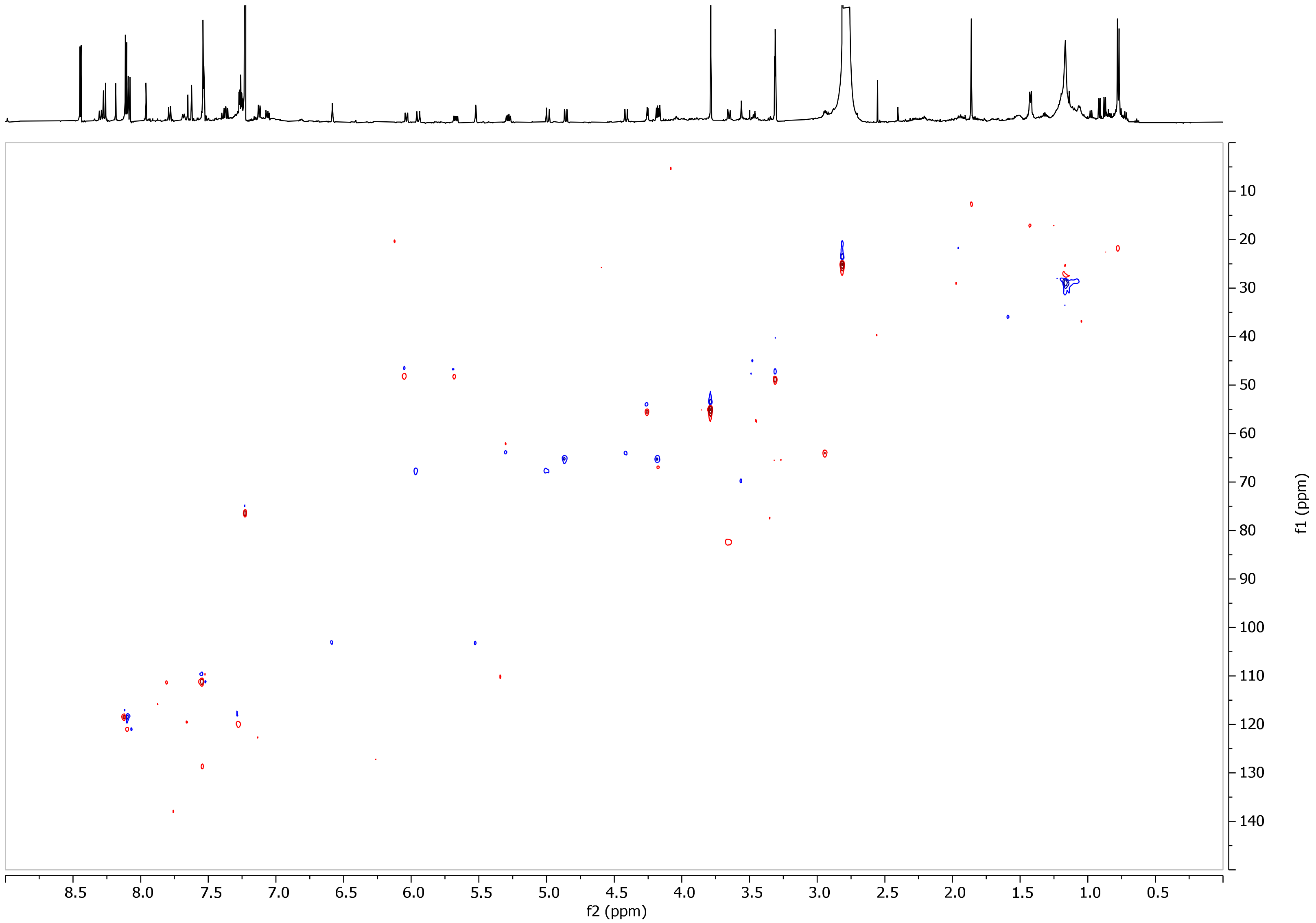


**Figure S37.** HSQC spectrum of compound **13** in $CDCl_3$:$CD_3OD$ (9:1).

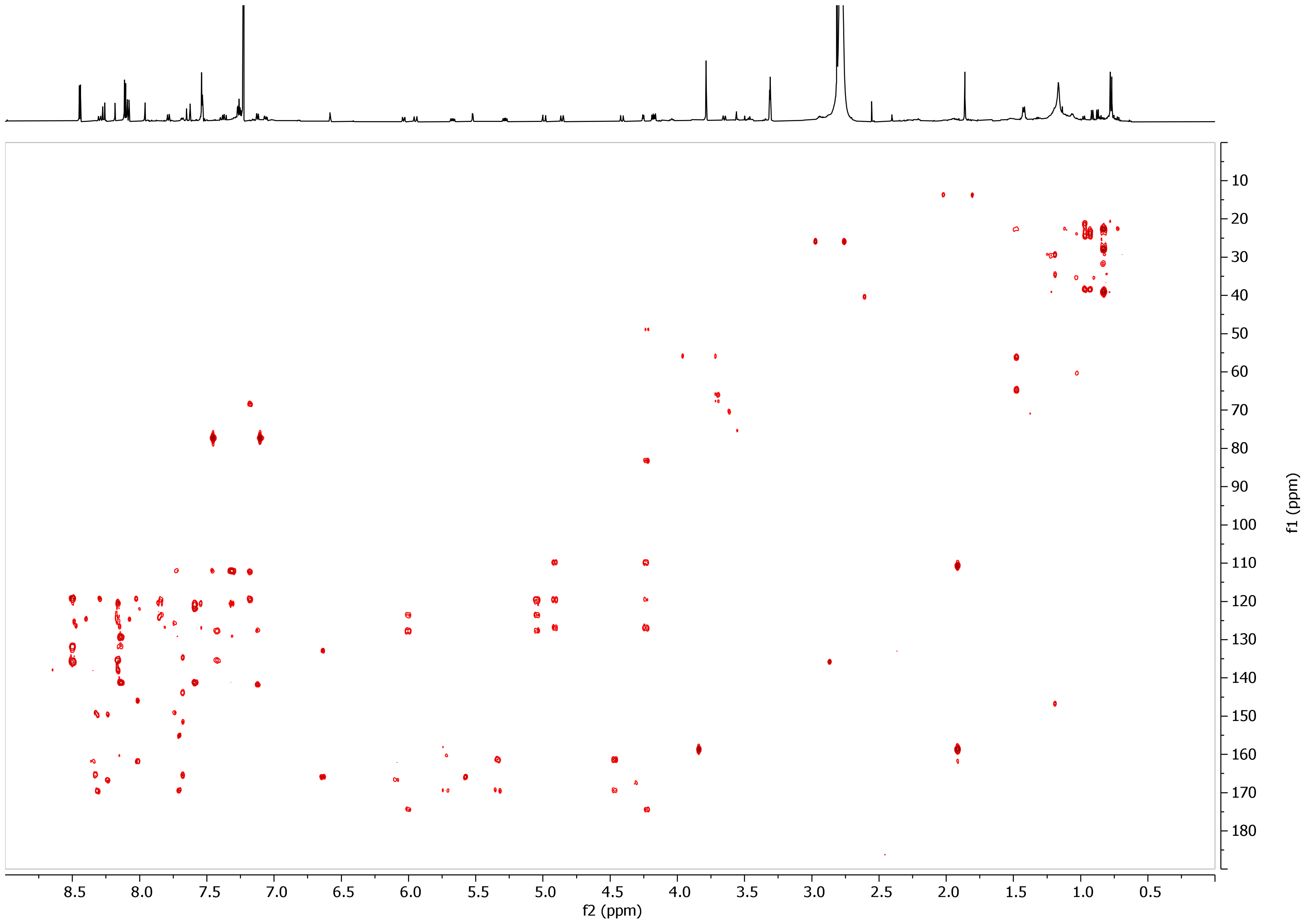


**Figure S38.** HMBC spectrum of compound **13** in $CDCl_3$:$CD_3OD$ (9:1).

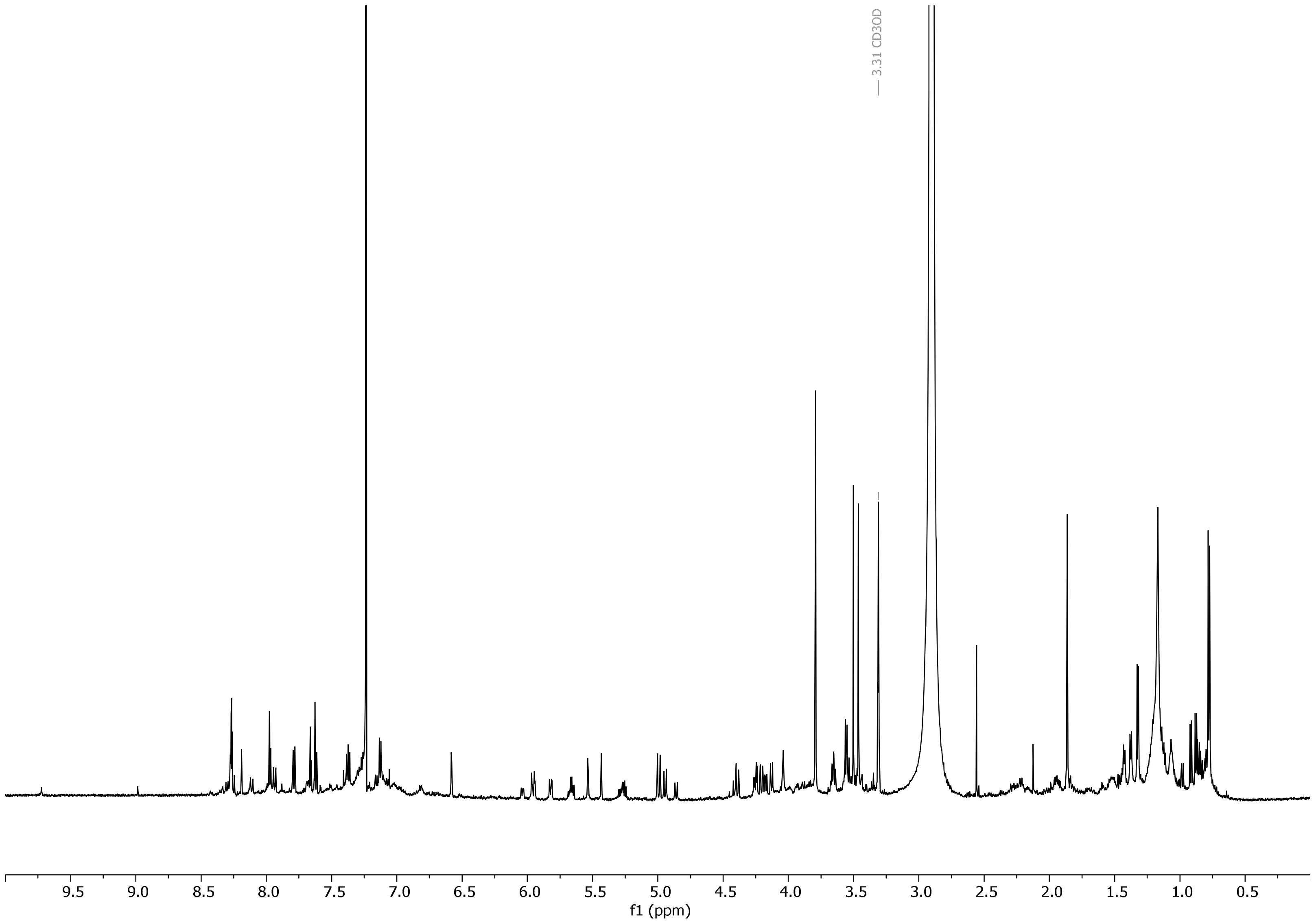


**Figure S39.** Proton spectrum of compound **14** in $CDCl_3$:$CD_3OD$ (9:1).

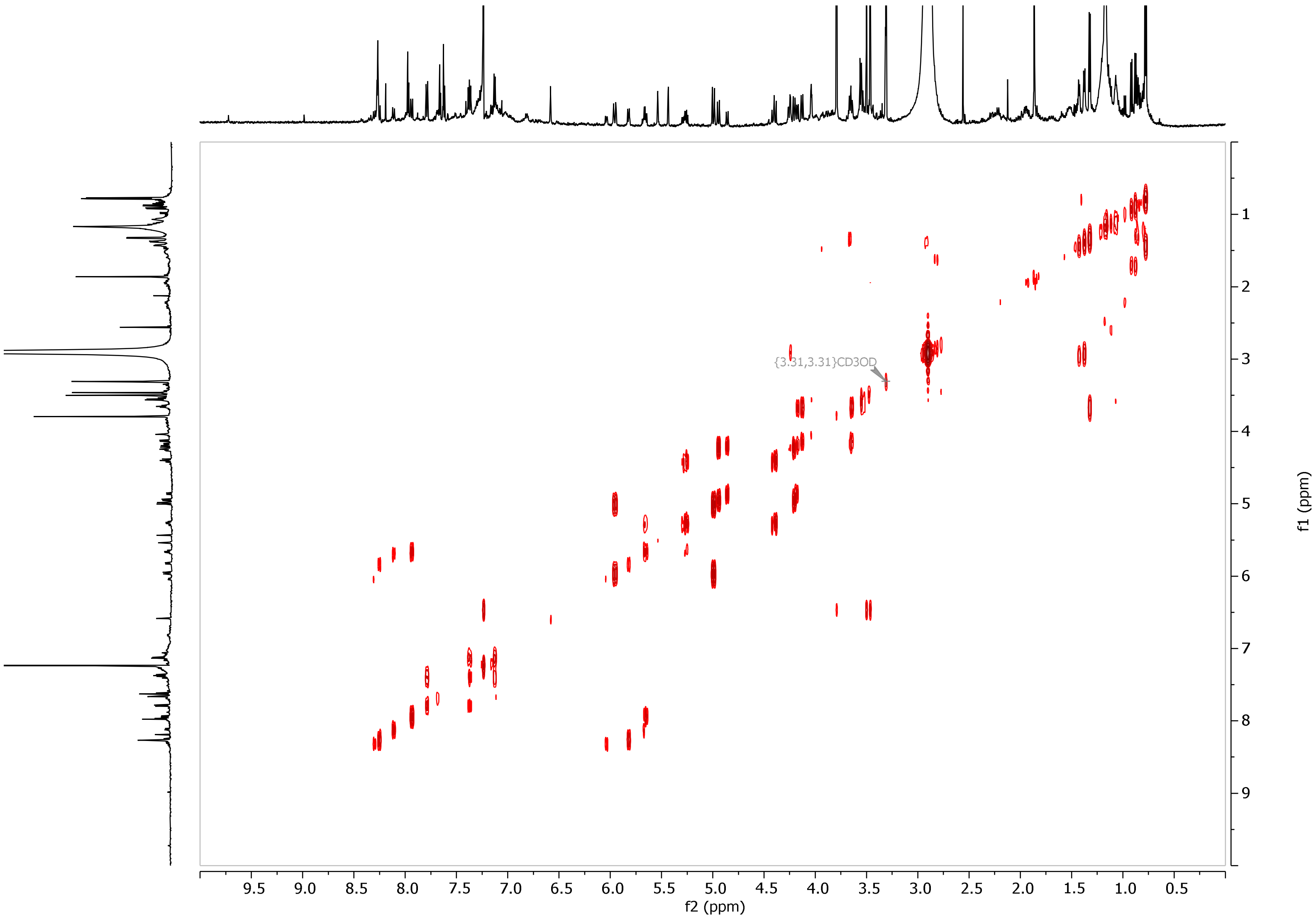


**Figure S40.** COSY spectrum of compound **14** in $CDCl_3$:$CD_3OD$ (9:1).

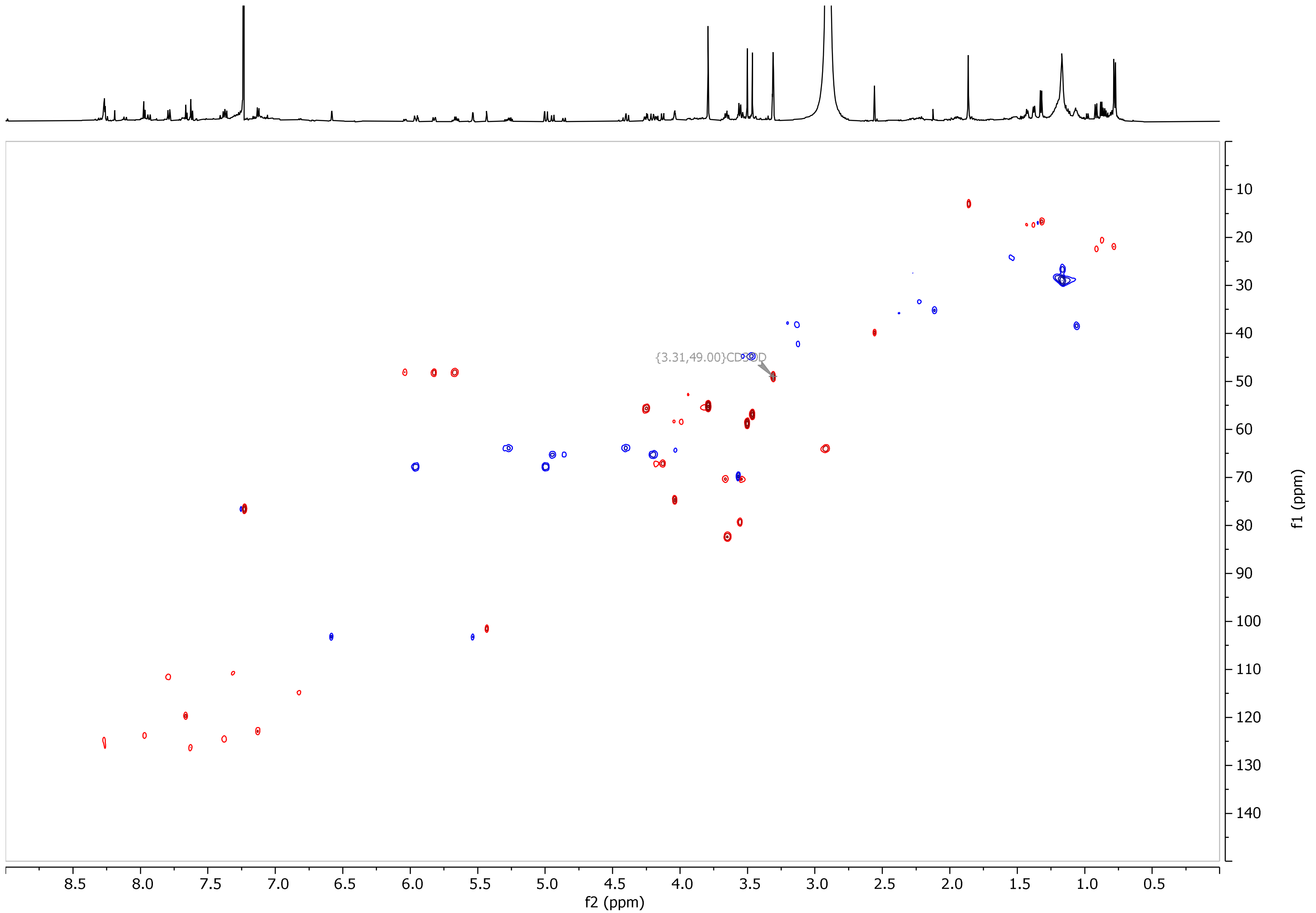


**Figure S41.** HSQC spectrum of compound **14** in $CDCl_3$:$CD_3OD$ (9:1).

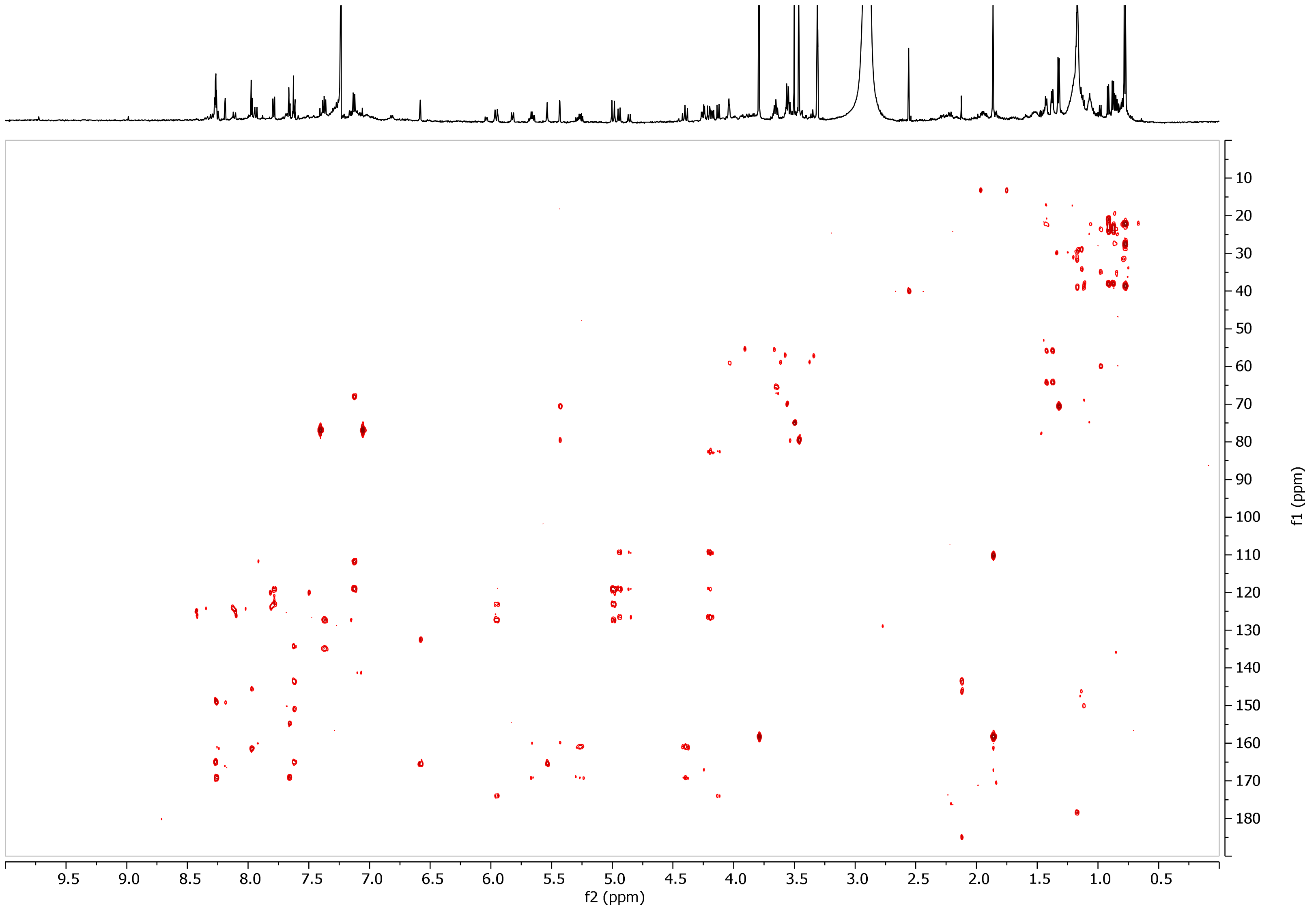


**Figure S42.** HMBC spectrum of compound **14** in $CDCl_3$:$CD_3OD$ (9:1).
